# Compartmentalized dendritic plasticity in the retrosplenial cortex integrates memories across time

**DOI:** 10.1101/2021.10.28.466343

**Authors:** Megha Sehgal, Daniel Almeida Filho, George Kastellakis, Sungsoo Kim, Jinsu Lee, Yang Shen, Shan Huang, Ayal Lavi, Giselle Fernandes, Sunaina Soans Martin, Irene Davila Mejia, Asli Pekcan, Melody Shana Wu, Won Do Heo, Panayiota Poirazi, Joshua T. Trachtenberg, Alcino J. Silva

**Affiliations:** Departments of Neurobiology, Psychiatry and Psychology & Integrative Center for Learning and Memory, UCLA, Los Angeles, CA; Institute of Molecular Biology and Biotechnology (IMBB), Foundation for Research and Technology, Hellas (FORTH), Vassilika Vouton, PO Box 1385, GR 70013 Heraklion, Crete, Greece; Department of Biological Sciences, Korea Advanced Institute of Science and Technology, Daejeon, Republic of Korea; Department of Psychology, The Ohio State University, Columbus, Ohio

**Author notes:** SENAI Institute of Innovation in Advanced Health Systems, University Center SENAI CIMATEC, Salvador, Bahia, Brazil (Daniel Almeida Filho).

## Abstract

Events occurring close in time are often linked in memory, providing an episodic timeline and a framework for those memories. Recent studies suggest that memories acquired close in time are encoded by overlapping neuronal ensembles, but the role of dendritic plasticity mechanisms in linking memories is unknown. Using activity-dependent labeling and manipulation approaches, longitudinal one- and two-photon imaging of somatic and dendritic compartments, and computational modeling, we show that memory linking is not only dependent on ensemble overlap in the retrosplenial cortex, but also on branch-specific dendritic allocation mechanisms. The same dendritic segments are preferentially activated by two linked memories, and spine clusters added after each of the two linked memories are allocated to the same dendritic segments. Our results demonstrate a causal mechanistic role for dendritic plasticity in memory integration and reveal a novel set of rules that govern how linked and independent memories are allocated to dendritic compartments.

## INTRODUCTION

Within the brain, pyramidal neurons use their elaborate dendritic structures to perform computations previously thought impossible for a single cell^1,2^. The molecular and cellular physiology that supports these complex computations within a single cell and how these computations influence ensemble activation, and thus animal behavior, are poorly understood. Active dendritic processes sculpt the flow of information from the synapse to the neuronal soma^3^. *Ex vivo* and *in vitro* studies demonstrate that focal synaptic activity on dendritic segments results in compartmentalized dendritic plasticity, which in turn regulates the integration and propagation of local dendritic signals to the soma, as well as impacts future induction of synaptic plasticity on these dendritic segments^4–8^. Although such localized plasticity within dendritic branches is likely to influence many neural processes, it is unclear whether and how this plasticity modulates memory.

Memory formation is a dynamic process, where single memories are stored, updated, and integrated within the framework of other pre-existing memories to drive adaptive behavior^9–11^. Recent studies in rodents have revealed that the overlap between the neuronal ensembles encoding different memories can link them, such that the recall of one leads to the recall of the other^12–14^. A similar process in humans is believed to mediate inferential reasoning^15,16^ and other forms of memory organization. Transient increases in neuronal excitability drive ensemble overlap^12,14^, but the neuronal locus as well as specific form of cellular plasticity underlying these changes is unknown.

Memory formation and retrieval are mediated by dendritic and synaptic processes^17,18^. Specifically, learning is mediated by input-specific synaptic plasticity^19–21^. In addition, the currents underlying learning-related changes in intrinsic excitability are dendritic in origin^22^. Yet, our understanding of the localized dendritic plasticity mechanisms that regulate the encoding and integration of memories is limited. Since experience-dependent dendritic plasticity is branch-specific^4–6,8^, and potentiation of dendritic spines can affect future plasticity at nearby spines on the same dendritic branch^6,23^, we hypothesized that two memories acquired close in time would be allocated to an overlapping population of dendritic branches, and that this mechanism drives linking of distinct memories. To test this hypothesis, we investigated the role of dendritic allocation mechanisms in contextual memory linking within the retrosplenial cortex (RSC), a brain region important for spatial and contextual memory processing^24,25^. We longitudinally tracked and manipulated RSC somas, dendrites, and spines as mice encoded distinct contexts to demonstrate that linked contextual memories are encoded within overlapping dendritic branches within the RSC.

## RESULTS

### Overlap in RSC ensembles representing linked memories

The overlap between neuronal ensembles encoding two memories (neuronal co-allocation) is critical for linking these memories^12–14^. However, it is unclear if such neuronal overlap is observed within the RSC neuronal ensembles, a brain region critical for encoding contextual memories. Thus, we first investigated whether RSC neuronal ensembles representing memories of two contexts explored close in time (i.e., linked memories)^12^ also display a higher overlap than two ensembles representing memories further apart (i.e., independent memories). We used a customized head-mounted miniature microscope (see Methods for details) to image GCaMP6f-mediated calcium dynamics in RSC neurons (Figures 1a-d and S1, 4599 putative RSC neurons, 132.9 ± 11.6 neurons per session,) while mice explored different contexts. We found a greater overlap between the RSC neuronal ensembles activated during encoding of two contexts explored on the same day (5 hours apart), than between the neuronal ensembles activated during encoding of two contexts explored one week apart (Figures 1e and S1). Notably, the increased overlap between neuronal ensembles for memories acquired 5 hours vs. one week apart was not due to differences in ensemble size (Figure S1), or the criteria used for neuronal cross-registration across days (Supplementary Table 1). Importantly, to establish the stability of our longitudinal imaging, mice underwent exposure to the same or different contexts 7 days or 5 hours apart. We discovered that there is substantial representational drift in the neuronal ensemble representing the same context 7 days apart (Figure S1e). These data are consistent with similar representational drift observed in various brain regions^26–33^. However, the neuronal ensemble representing two contexts seven days apart is significantly more stable when these contexts are the same vs. two distinct contexts (Figure S1f). Therefore, greater overlap between the RSC neuronal ensembles representing two contexts explored on the same day, than between the neuronal ensembles representing two contexts explored one week apart is unlikely to be due to representational drift alone or problems with longitudinal imaging itself. These data indicate that RSC neurons represent temporally proximate contextual memories using overlapping neuronal populations. These results provide strong evidence that the co-allocation of temporally proximate memories to overlapping neuronal ensembles may be a universal mechanism for memory linking.

**Figure 1.**
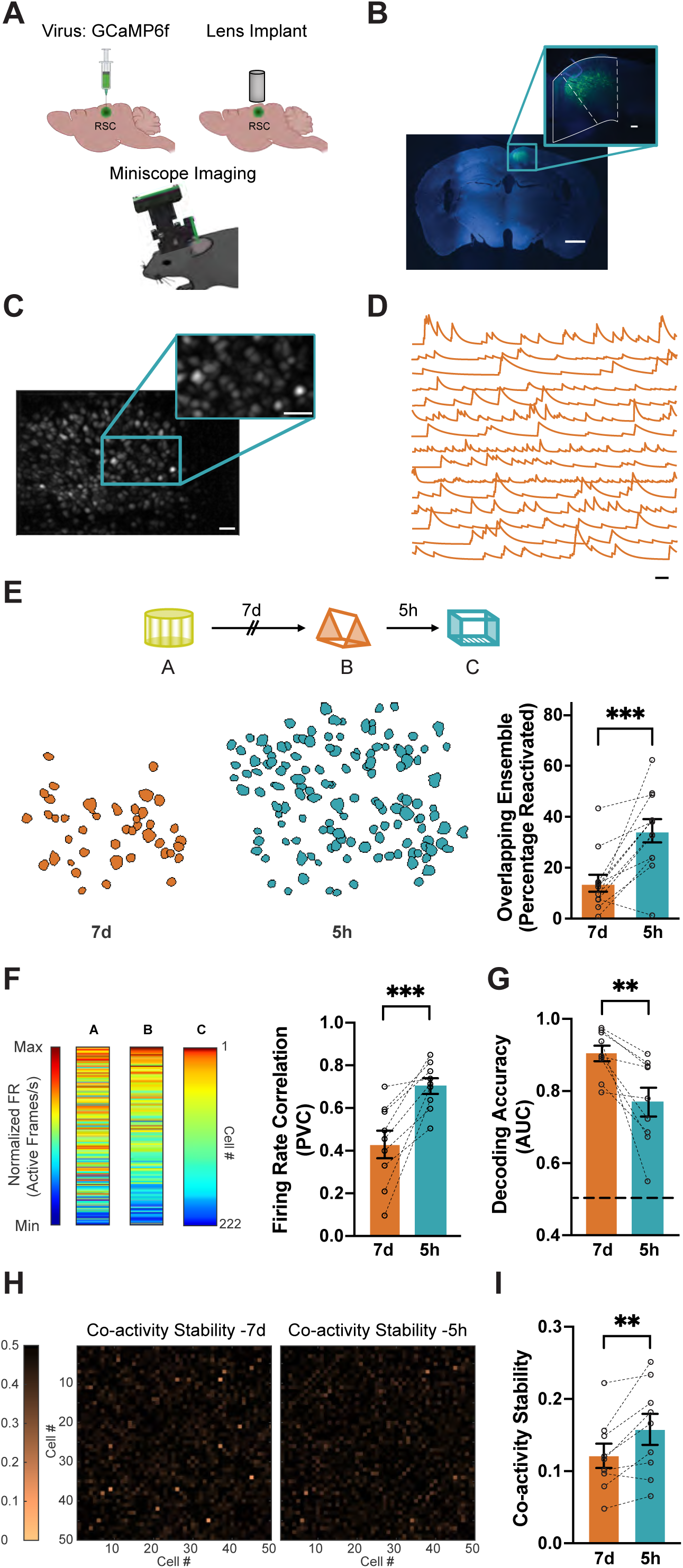
Overlapping RSC ensembles are recruited to encode contextual memories acquired close in time. (a) Miniscope methodology: GCaMP6f virus was injected into the RSC, and a Miniscope was used to image calcium transients in RSC neurons across repeated imaging sessions. (b) GCaMP6f expression within the RSC is limited to specific sub-layers of the RSC. Scale: 1mm; inset, 100 µm. (c) Example of maximum intensity projection of processed calcium signals during context exploration. Scale: 50 µm. (d) Representative calcium traces from 15 putative RSC neurons from one mouse. Scale: 30s. (e) Overlapping RSC ensembles encode distinct memories acquired close in time. Top: Mice were imaged while exploring three novel contexts (A, B, and C) separated by 7 days or 5 hours. Bottom left: Overlapping neurons in RSC ensembles in a representative mouse when contexts were separated by 7 days (orange) and 5 hours (blue). Bottom right: RSC neuronal ensembles display greater overlap when contexts were separated by 5 hours (5h) vs. 7 days (7d; n = 12 mice per group, paired t-test, t = 4.8). The physical contexts presented were counterbalanced to minimize any effect of context similarity. (f) Frequency of calcium transients in the RSC ensemble is more correlated when two contexts are explored close in time. Left: Frequency of calcium transients (active frames/s) for all RSC neurons from one mouse during each context exploration ranked based on their normalized frequency of calcium transients in context C. RSC neurons with a high frequency of calcium transients continue to fire at high rates when contexts are explored close in time (5 hours vs. 7 days apart). Right: Population Vector Correlation (PVC) for normalized firing rates is higher when contexts are explored close in time (i.e., 5 hours vs 7 days apart; n = 9 mice per group; paired t-test, t = 5.1). (g) A Naïve Bayes (NB) classifier is better at distinguishing two contexts explored 7 days apart vs 5 hours apart. The AUC (area under the curve) for the binary NB classification, using neuronal activity between sessions, is higher for sessions recorded 7 days apart than sessions recorded 5 hours apart (n = 9 mice per group; paired t-test, t = 3.5). Chance level performance of the classifier (AUC = 0.5) is represented by a dashed line. Spike probabilities were binned at 10 seconds in non-overlapping intervals. (h) Representative neuronal coactivity across sessions: The stability of neuronal coactivity across sessions is represented as the absolute difference in pairwise correlations between sessions (i.e. Pairwise correlation for Session 2 – Pairwise correlation for Session 1). Absolute difference in coactivity when contexts are explored 7 days (left panel) or 5 hours (right panel) for 50 cell pairs. Higher numbers (darker color) indicate more stable coactivity patterns. (i) Neuronal coactivity is more stable when contexts are explored close in time. The coactivity of neuronal pairs is more stable in two contexts explored 5h vs 7d apart (see Methods for details, n = 9 mice per group; paired t-test, t = 3.4). Data represent mean ± s.e.m. and each data point, * *p* < 0.05, ** *p* < 0.01, *** *p* < 0.001.

We reasoned that if, similar to other memory linking paradigms^12,14^, transient increases in intrinsic excitability in RSC drive neuronal ensemble overlap of linked memories, then the firing rate of RSC neurons should be similar for contexts explored close in time. We found that RSC neurons maintained a similar frequency of calcium transients for contexts explored within 5 hours compared to contexts explored 7 days apart (Figures 1f and S2). Consistent with the role of intrinsic excitability in memory linking, we found that the highly active cells (especially the top 10% of most active neurons) in a context are more likely to be reactivated in a different context 5h later compared to 7d later (Figure S3). Since RSC neurons encode distinct contexts using a firing rate code^25^, it is likely that neuronal firing dynamics within the RSC also impact the ability to decode context identity. Indeed, a Naïve Bayes (NB) classifier performed better at distinguishing sessions recorded a week apart relative to those recorded within the same day (Figures 1g and S2). Finally, we investigated the coactivity patterns of RSC neurons during these context explorations. Theoretical as well as experimental models suggest that groups of neurons with synchronized activity encode task-relevant information in the hippocampus, cortical, and subcortical regions^34–41^. However, the significance of such coactivity patterns during memory formation within the RSC is unclear. Therefore, we calculated the pairwise correlation (PWC) for each pair of RSC neurons within each session (PWC map) and found that these correlations are generally stable across imaging sessions (Figure S5). However, the across-session stability of these PWC maps (co-activity stability) is higher when contexts are explored on the same day compared to one week apart (Figure 1h, i), indicating that RSC neurons maintain patterns of coactivity when contexts are explored within the same day. Such synchronous firing may be key for the linking of contextual memories within the RSC. Together, these data indicate that overlapping RSC ensembles are activated when contextual memories are acquired close in time and that the dynamic activity of these overlapping ensembles may play a critical role in linking different contextual memories.

Although overlap in the underlying neuronal ensemble can serve to link these memories (our current findings as well as ^12–14^), these memories can still remain distinct^12,13^. To address how temporally proximate memories can be distinguished while being behaviorally linked, we calculated the functional connectivity difference (Euclidean distance (ED)) between correlation maps of neuronal activity from different sessions of the same animals when different and the same context were explored across 7d or 5h intervals (Figure S4a). Since the reactivation of neurons is biased towards the cells that were most active in the previous session, especially the top 10% firing rate (FR) cells, we assessed the importance of these groups of cells for computing the ED between correlation maps. We found that the exclusion of 10% of the most active cells from the correlation maps significantly increased the ED between correlation maps when mice explored distinct contexts 5h apart (Figure S4b, c). Similarly defined high FR cells did not affect the ED between correlation maps when the same or distinct contexts were explored 7d apart or the same context was explored 5h apart. The outsized contribution of high FR cells to representational similarity during exploration of two distinct contexts (vs the same context) 5h apart is consistent with their higher probability of reactivation (Figure S3). Overall, these data indicate that high FR cells within the RSC drive overlap as well as representational similarity at an ensemble level between distinct memories that are linked. On the contrary, representation of the same context is driven more equitably by high and low FR cells. Therefore, memory linking may be driven by highly active cells, while less active cells encode different contextual features that are relevant to sustain the independence between contextual experiences^40^.

### Overlap in RSC neuronal ensembles links memories close in time

To investigate the causal role of RSC neuronal co-allocation in linking contextual memories, we used the TetTag system^42^ to tag and manipulate the RSC neuronal ensembles activated during context exposures (Figure 2a, b). The TetTag system can capture neuronal ensembles in an activity-dependent manner, such that neurons with high firing rates during the behavioral epochs are selectively labeled^43^. Following a 10-minute context exposure, we found that 4.7 ± 0.42% of RSC neurons were labeled with ChR2-mCherry after 24 hours. In comparison, 8.5 ± 0.53% of RSC neurons were cFos-immunoreactive 90 minutes after a 10-minute context exposure. We confirmed that optogenetic reactivation of the RSC ensemble underlying a single contextual fear memory (~ 6.05 ± 0.53% of RSC neurons) induces fear expression in an otherwise neutral and novel context (Figures 2c and S6 a-c)^24^. Notably, fear expression following optogenetic reactivation within the RSC is distinct from similar results within the hippocampus ^44^ in that fear expression was sustained throughout the post-stimulation period and not just the Light-On epochs. These results are consistent with previously published findings^24^ and from this point onwards, freezing data during optogenetic reactivation are presented as a comparison between the baseline and post-stimulation period. Overall, our data confirm the critical role of RSC and associated brain circuits in processing contextual information^24,45–47^.

**Figure 2.**
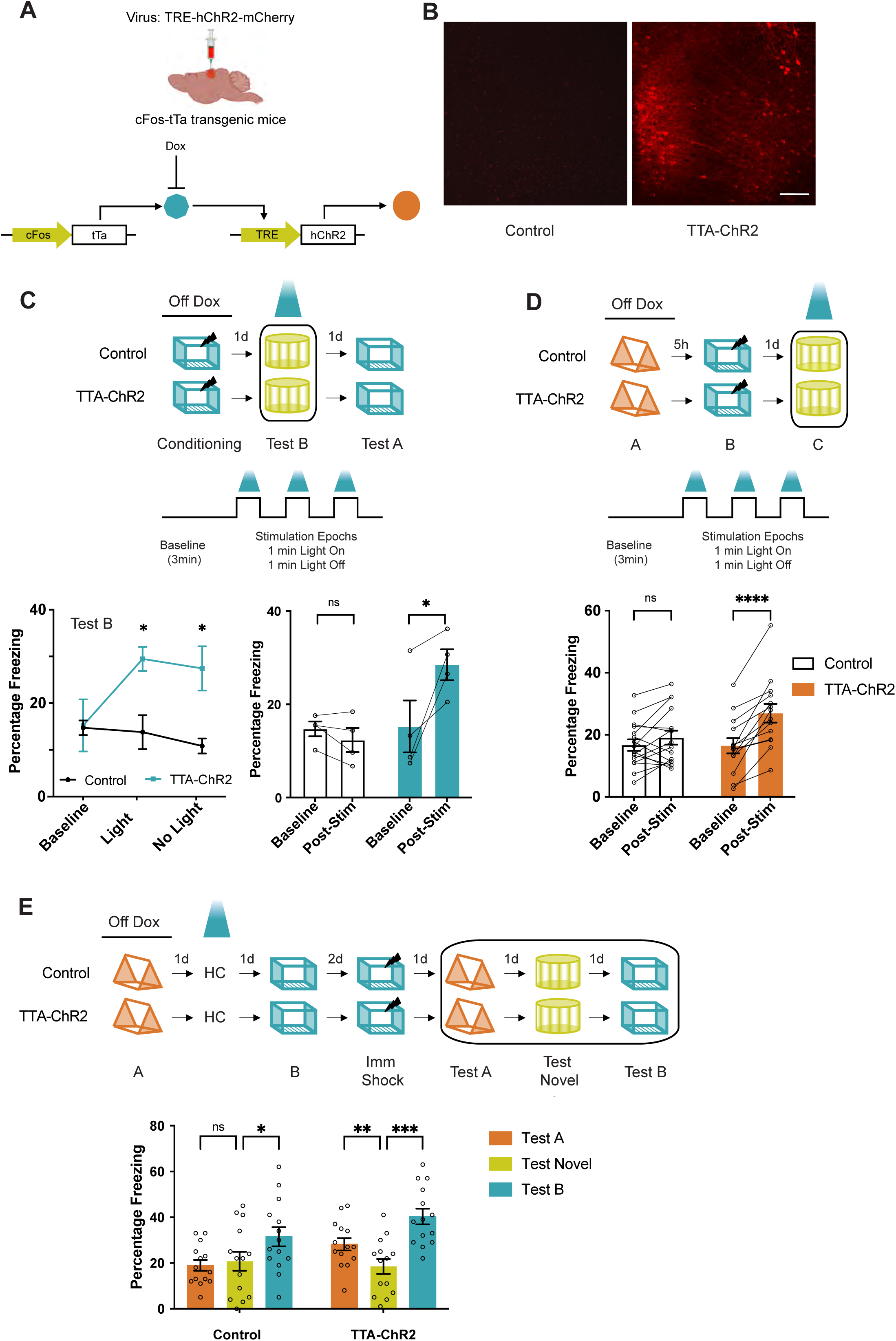
Overlap in RSC neuronal ensembles is sufficient to link contextual memories. (a) Schematic of the TetTag system: cFos-tTa or wildtype littermate mice were injected with the TRE-hChR2-mCherry virus. In the absence of doxycycline, the activation of the cFos promoter allows the expression of Channelrhodopsin (hChR2). (b) ChR2-mCherry expression in the RSC one day after fear learning in the experimental and control groups. Scale: 100 µm. (c) Optogenetic reactivation of an RSC ensemble underlying a fearful context is sufficient for fear expression: Top: Experimental setup: cFos-tTa (TTA-Chr2) mice and their wildtype littermates (Control) underwent bilateral viral injections (TRE-ChR2-mCherry) and optic cannula implants. Mice were taken off doxycycline chow (three days before contextual fear conditioning in context A: 2 footshocks, 2s, 0.7mA) to allow c-fos promoter driven tTA and Channelrhodopsin (ChR2) expression. Following contextual fear conditioning, mice were tested in a novel context (Test B) while the previously tagged neurons were activated. The following day mice were retested without any optogenetic manipulation in the training context (Test A). Middle: Optogenetic stimulation (473nm laser, 5 ms pulses, 5 Hz) was preceded by 3 mins of context acclimation, followed by three 1-minute trials of light pulses, separated by a 1-minute period of light off. Bottom: During Test B, TTA-Chr2 mice and control mice display similar freezing levels during the baseline period. Upon reactivation of the RSC neuronal ensemble tagged during contextual fear conditioning, TTA-Chr2 mice display more freezing compared to the control group during the post-baseline stimulation as well as non-stimulation epochs (n = 4 mice per group; Two-way RM ANOVA, group X time interaction, F (2, 12) = 6.95, *p* < 0.01, Uncorrected Fisher’s LSD, *p* < 0.05). (d) Optogenetic reactivation of an RSC ensemble underlying a linked memory is sufficient for fear expression: Top: RSC ensemble activated during the exploration of the first context (context A) was tagged. Five hours later, the second context (context B) was paired with a footshock. The following day, mice were tested in a third novel context (context C) with optogenetic reactivation of the tagged (Context A) RSC neuronal ensemble. Bottom: Reactivation of the RSC neuronal ensemble tagged during the linked context exploration (context A) increases freezing in cFos-tTa mice during the post-stimulation period, while the freezing in the control group remains unchanged (n = 16 and 14 mice for Control and cFos-tTa groups; Two-way RM ANOVA, F_Interaction_ (1, 28) = 12.5, *p* < 0.001; Sidak’s multiple comparisons test; Baseline freezing (Control vs. TTA-ChR2: p = 0.99); Post-Stim freezing (Control vs. TTA-ChR2: p < 0.05). (e) Reactivation of the RSC ensemble underlying the first context memory (context A) extends the temporal window for memory linking: Mice were allowed to explore 2 contexts (context A and B) separated by 2 days (a time interval when memories are not linked). On the day between the two context exposures, the RSC ensemble tagged during the first context visited (context A) was reactivated optogenetically. While control mice do not link the 2 contexts, reactivation of the first context ensemble leads to robust contextual memory linking: freezing in both previously explored contexts (Context A: linked context and Context B: shock context) is higher than the freezing in a novel context. The control group freeze similarly in context A and novel context, but the freezing in context B (shock context) is higher than freezing in a novel context (n = 14 mice each for Control and cFos-tTa groups; Two-way RM ANOVA, F_GroupXContext_ (2, 52) = 3.3, *p* < 0.04; Dunnett’s multiple comparisons test for contexts). The physical contexts presented were counterbalanced to minimize any effect of context similarity. Data represent mean ± s.e.m. and each data point, * *p* < 0.05, ** *p* < 0.01, *** *p* < 0.001.

When two contextual memories are acquired close in time, and one is paired with a fearful stimulus, the mice also consider the second neutral but ‘linked’ context as fearful (i.e., the two memories are linked)^12^. We asked whether the reactivation of RSC neurons engaged during exploration of the ‘linked’ context after contextual memory linking was sufficient to elicit freezing in mice (Figure 2d). Indeed, optogenetic reactivation of the ‘linked’ context’s neuronal ensemble in RSC alone was sufficient to trigger freezing in mice exploring a novel context (Figures 2d and S7). We confirmed that such fear expression did not result from the labeling of RSC neurons outside of the tagging window (e.g. during exposure to Context B) as doxycycline administration was sufficient to prevent tagging of additional neurons following exposure to another context, as well as following the injection of pentylenetetrazole (PTZ; Figure S7a). Additionally, the differences in optogenetic reactivation of fear between the TetTag mice and the control group did not stem from differences in contextual learning or linking of contextual memories. We tested the same mice in three different contexts on three consecutive days: the ‘linked’ context, another novel context, and the ‘shock’ context to assess their levels of memory linking, memory generalization, and conditioned responses, respectively. Both groups of mice displayed high levels of freezing in ‘linked’ (Context A) as well as ‘shock’ (Context B) contexts in comparison to freezing in the novel context (Figure S7). Thus, both groups of mice learned the context-shock association and linked the fearful context to the previously explored context (‘linked’ context), but only mice where the ‘linked’ context ensemble was reactivated using the TetTag system displayed fear in a novel context. First, these data indicate that reactivating the memory of a ‘linked’, but otherwise neutral context, was sufficient to elicit a conditioned response, a result that supports our hypothesis that the recall of one linked memory results in the recall of the other. Second, these findings also demonstrate that manipulation of neuronal ensembles just within the RSC can drive contextual memory linking.

While two contexts explored within a day are linked, contexts explored 2 or 7 days apart are not allocated to overlapping neuronal ensembles, and therefore are not linked^12,48^. We asked if we could link two distant contextual memories (acquired 2 days apart) by artificially biasing the involvement of a specific RSC neuronal ensemble in the encoding of both memories. With the TetTag system^42^, we tagged the RSC neuronal ensemble activated during a context exploration (Context A) and reactivated this ensemble the next day, one day before exposure to another context (Context B, Figure 2e). We reasoned that this would optogenetically reactivate the first memory, maintain the increase in neuronal excitability, and therefore force the recruitment of this same ensemble^49^ during the exploration of another context a day later. We allowed 24 hours for expression and then reactivated the RSC ensemble to allow sufficient expression of Channelrhodopsin post-tagging^43^. While two contexts explored 2 days apart are normally not linked, this optogenetic reactivation of the first contextual memory was sufficient to bridge this 2-day gap and drive the linking of two otherwise independent contextual memories (Figure 2e). We further confirmed the role of neuronal ensemble overlap in the RSC using a chemogenetic (Lentivirus DREADD^12^) system. We biased neuronal co-allocation of two distinct contextual memories by enhancing the neuronal excitability in a sparse population of RSC neurons before each context exploration (2 days apart, Figure S9). Like the optogenetic manipulation, artificially biasing co-allocation of two distant contextual memories to overlapping RSC ensembles using chemogenetics drives the linking of the two memories, such that the mice showed comparable freezing in both contexts (Context A and B). Additionally, we demonstrate that optogenetically activating a small but random population of RSC neurons between two context exposures (similar to Figure 2d) was not sufficient to link two independent contextual memories (Figure S8). Together, these data demonstrate that neuronal ensemble overlap in RSC is critical to link the memories of two distinct contexts.

### Overlap in dendritic ensembles encoding linked memories

Ours and previous results demonstrate that the allocation of contextual memories to overlapping neuronal ensembles is critical for linking contextual memories^12,48^. However, the intracellular processes that mediate neuronal overlap are poorly understood. Specifically, whether dendritic plasticity mechanisms contribute to neuronal overlap is unclear. Within the overlapping ensembles, linked memories are thought to be encoded by distinct synaptic changes (heterosynaptic) that allow the memories to maintain their distinct identities^50^. There are at least three dendritic hypotheses that could account for contextual memory linking. First, linked memories may be allocated to different dendritic branches within the encoding neurons (dis- allocation). Second, it is possible that linked memories are randomly allocated to the dendritic branches within the encoding neurons. Third, since experience-dependent dendritic plasticity is highly localized and can affect future plasticity at nearby spines on the same dendritic segment under certain conditions^6,23^, it is also conceivable that, following the first context exposure, localized changes in dendritic plasticity temporarily bias the activation of the same dendritic segments during a subsequent context exposure^51^. In the latter scenario, distinct synaptic changes on the same dendritic branches could drive the co-activation, and therefore the linking of the two memories. We propose that localized dendritic plasticity is a key mechanism driving neuronal ensemble overlap, since this plasticity could affect the propagation of synaptic inputs on specific dendritic segments to the soma. To distinguish between these different hypotheses, we used two-photon microscopy to investigate the functional and structural dynamics of the apical dendrites of layer V RSC neurons (Figures 3–5). Specifically, we targeted the apical dendrites of layer V RSC neurons because we have previously demonstrated that dendritic plasticity in the form of spine turnover and clustered spine addition on dendritic hotspots within these compartments facilitates single contextual memory formation^19^. We reasoned that the important role of these dendritic compartments in single contextual memory formation makes them an excellent candidate for co-allocation of dendritic plasticity following memory linking.

**Figure 3.**
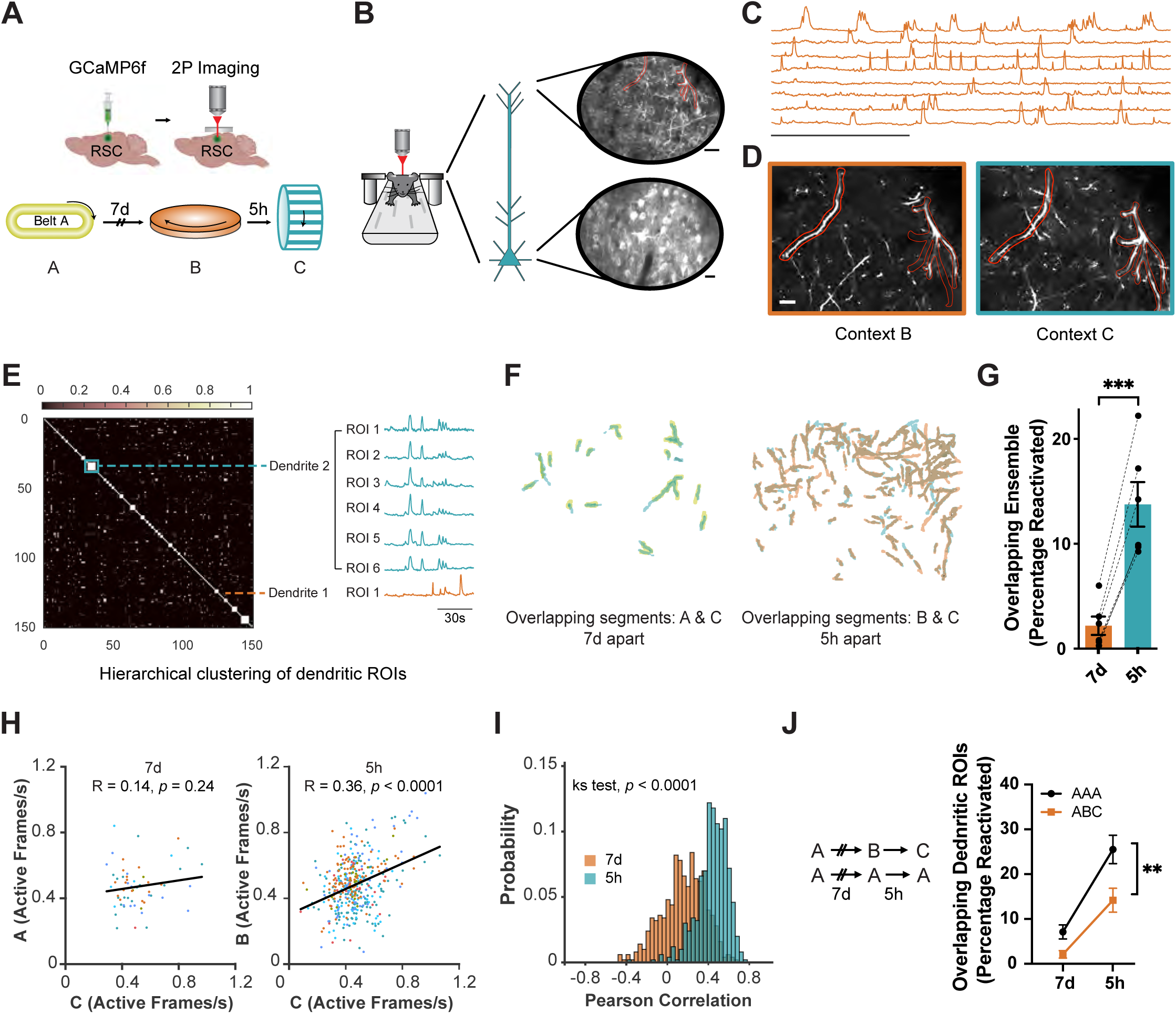
Overlapping dendritic segments encode memories of two contexts explored close in time. (a) Experimental setup: A 3 mm square craniotomy was made over the RSC spanning the midline, and GCaMP6f was injected bilaterally into the RSC. During imaging sessions, mice were head-fixed and experienced three novel contexts either 7 days or 5 hours apart as calcium transients were imaged using a two-photon microscope. Contexts comprised distinct visual, auditory, olfactory, and tactile cues and were counterbalanced (Figure S10). (b) RSC neurons and dendritic segments were tracked across 7 days. Top: Maximum intensity projection from one imaging session showing apical dendritic segments activated during the contextual experience. Two different dendritic segments are outlined in red. Scale: 20 μm. Bottom: Mean frame from one imaging session showing layer V RSC neurons. Scale: 10 μm. (c) Representative calcium traces from 8 putative RSC dendritic segments. Scale: 2 min. (d) Dendritic segments from (b) tracked across two imaging sessions 5 hours apart. Scale: 10 μm (e) Hierarchical clustering of RSC dendritic ROIs: Sorted cosine similarity matrix of 150 ROI pairs from one animal. Blue box and line depict the correlated calcium activity of 6 ROIs clustered as a single dendrite. Orange: Single ROI which was not clustered with ROIs in blue or any other ROI. (f) Overlapping dendritic segments reactivated when contexts are separated by 7 days (left) or 5 hours (right) from one mouse. (g) The same dendritic segments are more likely to be activated when context exposures are 5 hours (5h) apart vs 7 days (7d) apart. (Paired t-test; t = 9.2; *p* < 0.0005; n=6 mice). (h) Dendritic activity (active frames per second) is more correlated when dendrites are reactivated 5h vs 7d apart. Scatter plot of firing rate of all reactivated ROIs in context A (7d, 7 days apart) and context B (5h, 5 hours apart) as a function of firing rate in context C. Lines represent least-squares linear regression. Data from each mouse is represented in a different color. (i) Data from (h) were subsampled (30 ROI pairs, 500x) to generate a probability distribution of Pearson correlations (K-S test, *p* < 0.0001). (j) Dendritic overlap is greater when mice explore the same context vs distinct contexts (Two-way RM ANOVA, ABC vs AAA, *p* < 0.05; Sidak’s post hoc test, ABC, AAA (5h vs 7d) p< 0.001). Data represent mean ± s.e.m. and each data point.

First, we performed longitudinal calcium imaging of the somas as well as apical dendrites of layer V RSC neurons (see Methods for details, Video 1, Figure 3a-b), while mice explored distinct contexts in a head-fixed setting (Figure S10). Consistent with our results with RSC neuronal activation in freely moving mice (Figure 1), head-fixed mice also represent two contexts experienced close in time by recruiting overlapping RSC ensembles (Figure S11). These data reveal the similarities in the neuronal encoding of temporally-proximate memories in head-fixed and freely moving mice indicating the generalizability of our findings across two imaging modalities. We next assessed the degree of overlap among RSC dendritic branches when contexts are explored close in time. We imaged apical dendrites (in layer 1 of the RSC, ~ 30 μm from the pia mater) following GCaMP expression of layer V RSC neurons (see Methods for additional details). We found that the same dendritic ROIs were preferentially reactivated as mice explored two distinct contexts within the same day, but not a week apart (Figures 3e-g, j and S12a). Consistent with the role of NMDA receptor activation in memory linking^12^ as well as clustered spine formation in the RSC^19^, reactivation of dendritic segments required NMDA receptor activation during the first context exposure (Figure S13).

The extent and prevalence of independent dendritic and somatic events during calcium imaging, the causality or direction of their interdependence, and the factors that affect these are poorly established^52–55^. For our experiments, we sought to minimize the effect of backpropagating action potentials and global dendritic transients by imaging RSC apical tuft dendrites, as apical tuft branches display a degree of independence from one another and somatic calcium events^55^. To account for highly correlated calcium transients across ROIs driving our observed effects, we performed a hierarchical clustering analysis to group segmented ROIs into single dendritic units when their calcium dynamics are highly correlated (Figure 3e, see Methods for analysis details). We found similar proportions of clustered ROIs among reactivated and overall segmented ROI populations (Reactivated ROIs: 0.85 ± 0.02; overall ROI population: 0.86 ± 0.02; *p* = 0.3). Clustering segmented ROIs in this way (to account for global dendritic transients or backpropagating action potentials contaminating our results) did not change our observed effects (Figures 3f-g and S12a). The clustered ROIs within reactivated segments maintained high within-cluster correlated activity across sessions, demonstrating the robustness of our clustering algorithm as well as the longitudinal coupling of these ROIs (Figure S12c). It is still possible that there is a one-to-one correspondence between our reactivated neurons and dendritic segments (all clustered ROIs), but we believe this is unlikely given the low levels of clustering (mean cutoff value = 0.13 ± 0.01, 1.15 ± 0.03 ROIs per cluster, Figure S12b) and the large difference between neuronal and dendritic overlap in our head-fixed experiments (Figure 3g and S11). It is possible but unlikely that the differences in neuronal and dendritic overlap are due to a lower signal-to-noise ratio (SNR) during dendritic imaging. Neuronal overlap using one-photon calcium imaging (Figure 1) where SNR is usually markedly lower than two-photon imaging was more similar to neuronal overlap in two-photon imaging conditions (Figure S11) and different than the degree of dendritic overlap. Importantly, these results did not rely on using any particular clustering criteria as clustering cutoffs that consistently resulted in clustered ROIs within shuffled distributions with randomized activity also yielded low cluster sizes in the experimental dataset (Clustering cutoff = 0.3, 1.39 ± .06 ROIs per cluster, see Methods) as well as similar overlap results (*p* <0.001).

Next, we analyzed calcium transient frequencies within the reactivated dendrites during two contexts explored 5 hours apart and found that these were highly correlated (Figure 3h). We did not observe a significant correlation between calcium transient frequencies in reactivated dendrites when contexts were explored 7 days apart (Figure 3h). To confirm that this analysis was not biased by differences in the number of reactivated dendrites for two context exposures 7 days or 5 hours apart, we repeatedly (500X) subsampled 30 reactivated ROIs (see Methods) from each condition to generate a probability distribution of correlation coefficients (Figure 3i). This subsampling analysis confirmed that calcium transient frequencies are more correlated when contexts are explored close in time. These data indicate that the synaptic drive and the local excitability mechanisms driving dendritic activity are maintained during the encoding of linked memories.

Finally, we asked if dendritic overlap was different when animals explored the same or different contexts at 7 days or 5 hours apart (Figure 3j). Similar to our data demonstrating neuronal overlap (Figure S1), we found that dendritic ROIs were more likely to be reactivated closer in time (5h vs 7d) whether animals experienced the same or different contexts (Figure 3j). In addition to this effect of time, we confirmed that dendritic representations are more similar for the same context vs different contexts irrespective of time. Our data here (Figure 3) are consistent with the hypothesis that local dendritic mechanisms govern the allocation of two contextual memories encoded close in time to the same dendritic segments^51^. Next, we used structural imaging of RSC apical dendrites to confirm and extend these findings.

### Linked memories bias spine remodeling to overlapping dendritic segments

Given that overlapping dendritic segments are activated when encoding contexts that are experienced close in time, we next investigated whether learning-related spine dynamics were also evident on the same dendritic segments when contextual memories are acquired close in time. Within the RSC, formation of a contextual memory is accompanied by structural plasticity at apical dendritic branches of layer V RSC neurons, such that behavioral performance is positively correlated with clustered spine addition on small stretches (~5μm) of a dendritic segment^19^. These data are consistent with the clustered plasticity hypothesis, and indicated that experience-dependent spine remodeling is spatially restricted in a branch-specific manner^5,6,19^. We used *in vivo* two-photon microscopy to image spines on the RSC apical dendrites of Thy1-YFP-H mice following multiple context exposures (Figure 4a-d). As structural imaging involved mice being imaged under anesthesia, we confirmed that mice were still able to link the memories for distinct contexts under these conditions (i.e., ~40mins of anesthesia following context exposure: ~ length of the spine imaging session, Figure S14). We quantified the spines added or lost following each context exposure and found that relative to spine dynamics during a baseline period, novel context exposure does not change overall spine addition, spine loss, or spine turnover (Figure S15a, b). However, following context exposure, new spines were added in clusters (i.e., within 5μm of each other); and a shuffling analysis showed that the number of these clustered spines was significantly above chance levels (*p* = 0.009; Figure S15c). In contrast, clustered spine addition in the control group - that went through all the same imaging procedures, but that was never exposed to a novel context - was at chance levels (p = 0.14; Figure S15c). Finally, spine formation or clustering was not correlated with pre-learning turnover (HC, ρ = 0.1 and 0.09 respectively; Context exposure group, ρ = 0.07 and 0.02; all *p* values = ns). We found that new spine formation following learning was correlated with spine density in control conditions (HC, ρ = 0.43, *p* = 0.007) but not following a context exposure (ρ = 0.23, ns). Hence, consistent with previous findings^19^, novel context exploration results in clustered plasticity in RSC dendrites.

**Figure 4.**
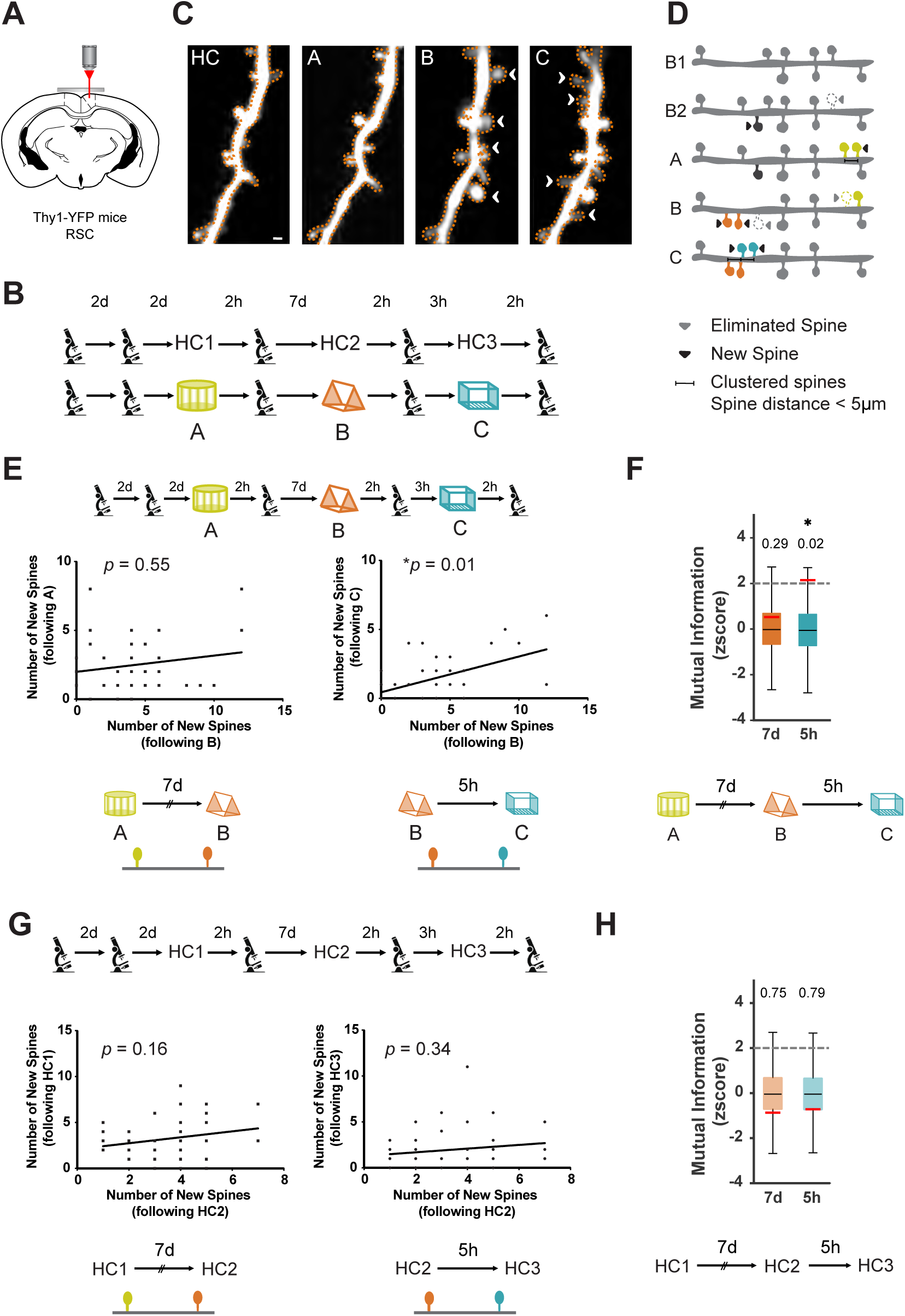
Spines are added to overlapping dendritic segments following memory linking. (a) Experimental setup: A 3 mm square craniotomy was made over the RSC of Thy1-YFP mice. Apical RSC dendrites were imaged to measure context exposure and memory linking related spine dynamics. (b) Mice received contextual exposure to three distinct contexts 7 days or 5 hours apart. Control mice were imaged at time-matched intervals without context exposure. (c) Representative example of spine dynamics during longitudinal imaging showing clustering of new spines following linked memory formation. Gained spine: white arrowhead. HC: Last baseline imaging session; A, B and C: Exposure to contexts A, B and C respectively. Scale: 1 μm. (d) Schematic of various spine dynamics (spine addition, elimination, and clustering) measured. (e) New spines are likely to be added to the same dendritic segments when contexts are explored close in time. Left: Number of new spines added to a dendritic segment following Context A and B exposure 7 days apart are not correlated (ρ = 0.09, *p* = 0.55). Right: Number of new spines added to a dendritic segment following Context B and C exposure 5 hours apart are correlated (n = 6 mice, ρ = 0.37, *p* = 0.01). Spearman’s correlation was used. Alpha level was adjusted to 0.025 to account for multiple comparisons. (f) Mutual information between new spines added at 7 days or 5 hours apart is higher for spines added following context exposures 5 hours vs 7 days apart. Observed values (red line) were compared to the z-score of a chance distribution. (g) For home cage (HC) controls, the numbers of new spines added to a dendritic segment are not correlated whether imaging sessions are separated by either 7 days (right, ρ = 0.22, *p* = 0.16) or 5 hours (left, ρ = 0.15, *p* = 0.34; n = 5 mice). (h) Mutual information between new spines added at 7 days or 5 hours is unchanged in control mice. Observed values (red line) were compared to the z-score of a chance distribution. The physical contexts presented were counterbalanced to minimize any effect of context similarity.

Since our results showed reactivation of the same dendrites during context exposures close in time, we next investigated the possibility that spines added following these context exposures also tend to be added to the same dendritic segments (Figure 4e-h). We found a positive correlation between the number of spines added to the same dendritic segments following two context exposures experienced 5 hours apart (Figure 4e). In contrast, the number of spines added to the same dendritic segments following context exposures one week apart were not correlated (Figure 4e). Spine addition was also not correlated in the control group that experienced the same imaging procedures without context exposures, at either the 5 hours or 7 days time intervals (Figure 4g). Next, we shuffled the number of new spines added to a dendritic segment following imaging sessions 5 hours or 7 days apart to generate a shuffled distribution. This procedure was repeated (10,000X) to generate a probability distribution of correlation coefficient values from this shuffled data. Our observed correlation coefficient between the number of spines added to a dendritic segment following two context exposures 5 hours apart (ρ = 0.37) was unlikely to be observed in this shuffled distribution (*p* = 0.006). In contrast, the number of spines added to a dendritic segment when two distinct contexts were explored 7 days as well as under home cage conditions are not correlated (also see Figure 4e, g) and not statistically different from correlation coefficients observed following the shuffling procedure (AB, 7d apart, *p* = 0.3; HC, 5h apart, *p* = 0.2; HC, 7d apart, *p* = 0.1).

We also calculated the Mutual Information (see Methods) contained in the number of spines added following two context explorations 5 hours apart and found that spine addition following the encoding of one context is predictive of the number of spines added following the encoding of a linked context (Figure 4f). This was not true for other imaging conditions (i.e., when contexts are explored 7 days apart; Figure 4f) or in the home cage controls (Figure 4h). Furthermore, the number of spines lost was not correlated whether the imaging sessions were conducted 7 days or 5 hours apart in both the control and experimental groups (Figure S16). Finally, we compared the distribution of newly added spines across groups. The correlation coefficient generated by distributions of newly added spines following two context exposures 5 hours apart was statistically different than the correlation values generated in other conditions (Fisher transformation; Experimental 5h vs HC 5h, Z = 1.8; *p* = 0.03; Experimental 5h vs Experimental 7d, Z = 1.67; *p* = 0.047; HC 7d vs HC 5h, Z = −0.42; *p* = 0.33). To control for the differences in the number of dendritic branches imaged under different conditions, we subsampled 40 dendritic branches from each condition (10,000X) to obtain a distribution of Spearman Correlations and Mutual Information for each condition. We found that mean Spearman Correlation, as well as Mutual Information values, are higher when mice explore two novel contexts 5h apart compared to all the other conditions (Figure 5e, f; *p* < 0.001). These data indicate that spine addition is biased to the same RSC dendritic segments when contextual memories are linked (acquired 5 hours apart), but not when these memories are independent (acquired 7 days apart).

**Figure 5.**
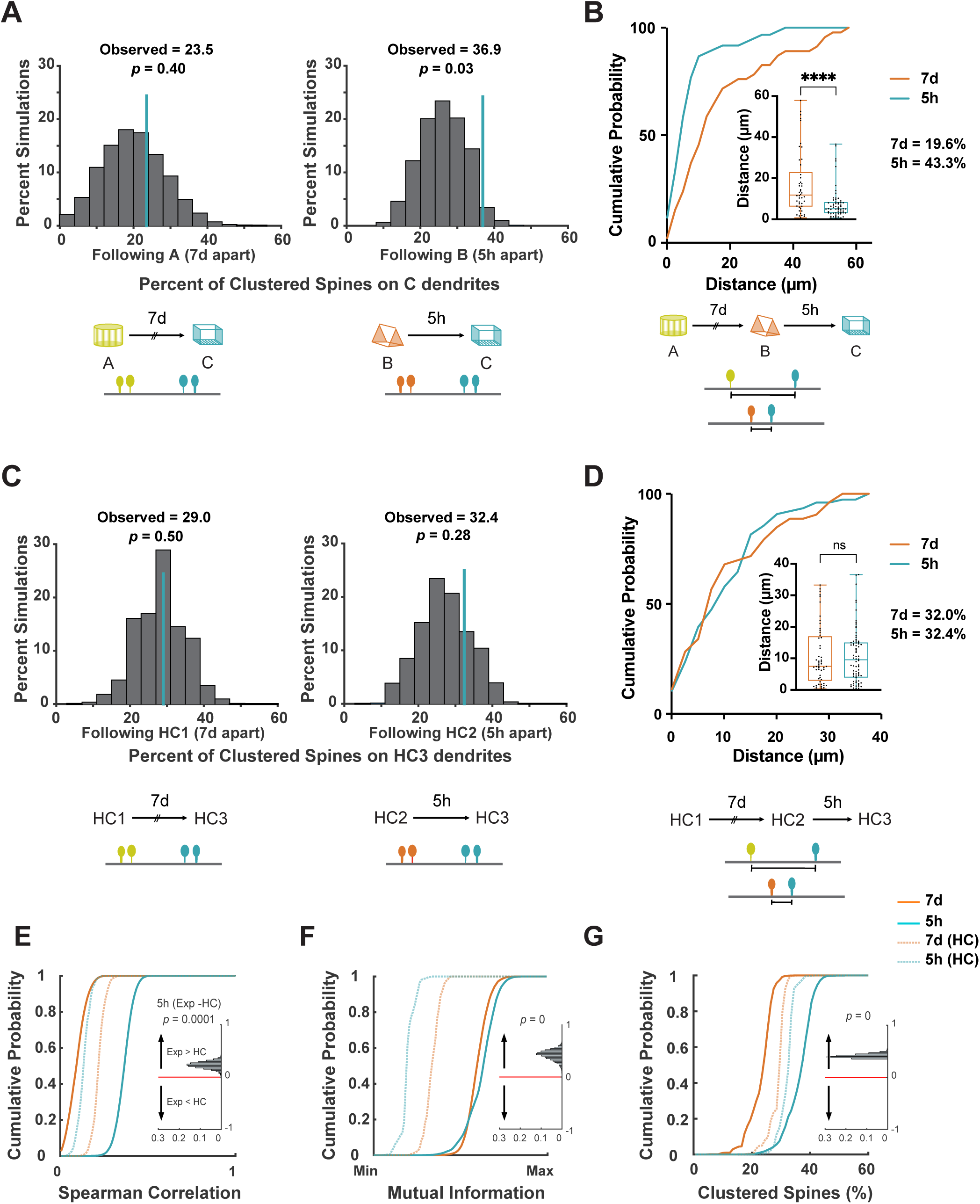
Overlapping dendritic segments gain spine clusters following memory linking. (a) Co-allocation of clustered spines to dendritic segments: The probability that clustered spines are co-allocated to the same dendritic segment by chance following two imaging sessions was calculated by randomly distributing clustered spine positions following one imaging session. Left: The percentage of clustered spines co-allocated to the same dendritic segments following contexts A and C (explored 7 days apart) are at chance levels. Right: In contrast, the percentage of clustered spines co-allocated to the same dendritic segments following contexts B and C (explored 5 hours apart) is higher than expected by chance. n = 6 mice; 10000 permutations. (b) New spines following context exposures 5 hours, but not 7 days, apart are formed close to one another (KS test, *p* <0.0001). Inset: Average distance between newly formed spines following exposure to contexts A and C (7d) or contexts B and C (5h). (Mann Whitney, *p* <0.0001) (c) For home cage (HC) controls, the percentage of clustered spines that were added to segments also containing clustered spines in a previous imaging session (7 days or 5 hours prior) were at chance levels; n = 5 mice; 10000 permutations. (d) New spines formed in home cage control mice do not co-cluster for imaging sessions 5 hours or 7 days apart (KS test). Inset: Average distance between newly formed spines in home cage controls 7-days or 5-hours apart. (Mann Whitney test). (e) Cumulative distribution of Spearman’s rho (ρ) values calculated by randomly subsampling 40 dendritic branches 10,000x for each condition. ρ values quantify the rank correlation between the number of new spines added to the same branch between two time points 5h or 7d apart. Inset demonstrates that the Spearman’s rho (ρ) is higher for resampled experimental vs HC group at 5h (*p* < 0.001). (f) Cumulative distribution of mutual information (MI) values calculated by randomly subsampling 40 dendritic branches 10,000x for each condition. MI values quantify the amount of information obtained about the new spines added to a branch by observing the number of new spines added to the same branch in a previous session 5h or 7d before. Inset demonstrates that the Mutual information is higher for resampled experimental vs HC group at 5h (*p* < 0.0001). (g) Cumulative distribution of values of the percentage of clustered spines among 40 dendritic branches randomly subsampled 10,000x for each condition. The percentage of clustered spines values quantify, among all clustered spines added after a session, the percentage added to the same branches that received clustered spines in a previous session 5h or 7d before. Inset demonstrates that the probability of gaining a clustered spine on a segment already containing a clustered spine during a previous session is higher for resampled experimental vs HC group at 5h (*p* < 0.0001). The physical contexts presented were counterbalanced to minimize any effect of context similarity.

Next, we analyzed spine dynamics looking for co-allocation of clustered spines (i.e., whether dendritic segments that gained clustered spines during a context exposure were also the ones that gained clustered spines during a previous context exposure). We found that the probability of the same dendritic segments gaining spine clusters following exposures to two contexts was at chance levels when these contexts were explored 7 days apart (*p* = 0.4, Figure 5a). Importantly, spine clusters were more likely to be added to the same dendritic segments when contexts were explored within the same day (5 hours apart; *p* = 0.03, Figure 5a). In contrast, the addition of clustered spines in the control group was random: the probability of the same dendritic segments gaining spine clusters during two imaging sessions was at chance levels whether the imaging sessions were 5 hours or 7 days apart (*p*= 0.28 and 0.5 respectively; Figure 5c). Using a similar analysis as for Figures 5e and f, we found that the probability that clustered spines were added to a dendritic branch already containing clustered spines was higher for an imaging session when two contexts were explored 5h apart in comparison to an imaging session at the same time interval under HC condition (Figure 5g, *p* < 0.0001). Thus, synaptic plasticity in the form of clustered spine addition following the encoding of linked memories is biased to overlapping dendritic segments.

Finally, we asked whether new spines added following exposure to two linked contexts were added close to one another, or if they cluster with each other. While 43.3% of newly formed spines following the last context exposure were clustered with the spines added following the context exposure 5 hours before (average distance between nearest neighbors = 7.7 ± 1.0 μm), only 19.6% of the new spines were clustered with new spines from the context exposure a week before (average distance between nearest neighbors = 17 ± 2.3 μm; Figure 5b). In control mice that did not receive context exposures, newly formed spines following the last imaging session clustered with new spines from the previous imaging session at similar rates whether the last session imaging was 5h or 7d before (5h: 32.4%, average distance between nearest neighbors = 10.7 ± 1 μm; 7d: 32.0%, average distance between nearest neighbors = 10.7 ± 1.3 μm, Figure 5d). Importantly, the cumulative frequency distribution as well as the average distance between nearest neighboring spines are statistically different between the experimental and HC group even when the imaging sessions were performed 5 hours apart (KS test, *p* <0.005 and Mann Whitney, *p* <0.05 respectively). Thus, new spines and spine clusters are added to overlapping dendritic segments following the formation of linked memories, and these newly formed spines cluster with each other. It is likely that such synaptic clustering can facilitate non-linear summation of dendritic inputs^56^, which would result in more robust propagation of inputs to the soma resulting in increased somatic firing^1,57–60^. Indeed, clustered spines are more effective at influencing neuronal spiking and, thus, the tuning properties of a neuron^56,61,62^. We demonstrate that following the encoding of two linked memories, spine clusters are added to the same dendritic branches. The addition of new spines in clusters to these dendritic branches could facilitate future ensemble activation. Together, the structural, as well as functional imaging data from RSC dendrites, indicate that the same dendritic branches are recruited to encode contextual memories formed close in time (i.e., linked memories).

### Optogenetic reactivation of a tagged dendritic ensemble links two independent memories

Next, we tested whether such dendritic co-allocation is sufficient for linking contextual memories. To investigate the behavioral significance of RSC dendritic co-allocation, we modified an optogenetic tool for tagging and manipulating previously activated dendritic segments: We leveraged the activity-dependent labeling of the cFos-tTA system, and combined it with the dendritic targeting element (DTE) of Arc mRNA, which is selectively targeted and locally translated in activated dendritic segments following learning (Figure 6a,b)^63,64^. This novel approach allowed us to manipulate dendritic activity by expressing a light-sensitive ion channel (Channelrhodopsin) in recently activated dendritic segments^18,65,66^ of RSC neurons that underlie the contextual memory trace. Since our tagging strategy was designed to mimic the well-documented activity-dependent increase of Arc mRNA expression in recently activated dendrites^67–70^, we first verified whether Arc mRNA is co-localized to the tagged dendritic segments. Following DTE-based labeling, mRNA encoding the fluorescent tag 5h after exposure to a novel context is more likely to colocalize near Arc mRNA in dendritic compartments (Figure S17a) indicating that our strategy can target activated dendrites. Next, we confirmed that dendritic segments labeled using a DTE-based method were also more likely to be reactivated upon re-exposure to the original tagging stimuli, i.e. exposure to the original context. Several lines of evidence have established that synaptic activity results in rapid phosphorylation (2-7 mins) of Cofilin protein in the synapse ^71,72^. We found that PSD-95 puncta on dendritic segments labeled using the DTE-based tagging method displayed an increase in phosphorylated form of Cofilin protein in comparison to PSD-95 puncta that were not present on labeled dendrites (Figure S17b). These data are consistent with previous reports^18,65,66^ and support that the DTE-based labeling allowed us to tag dendrites in an activity-dependent manner.

**Figure 6.**
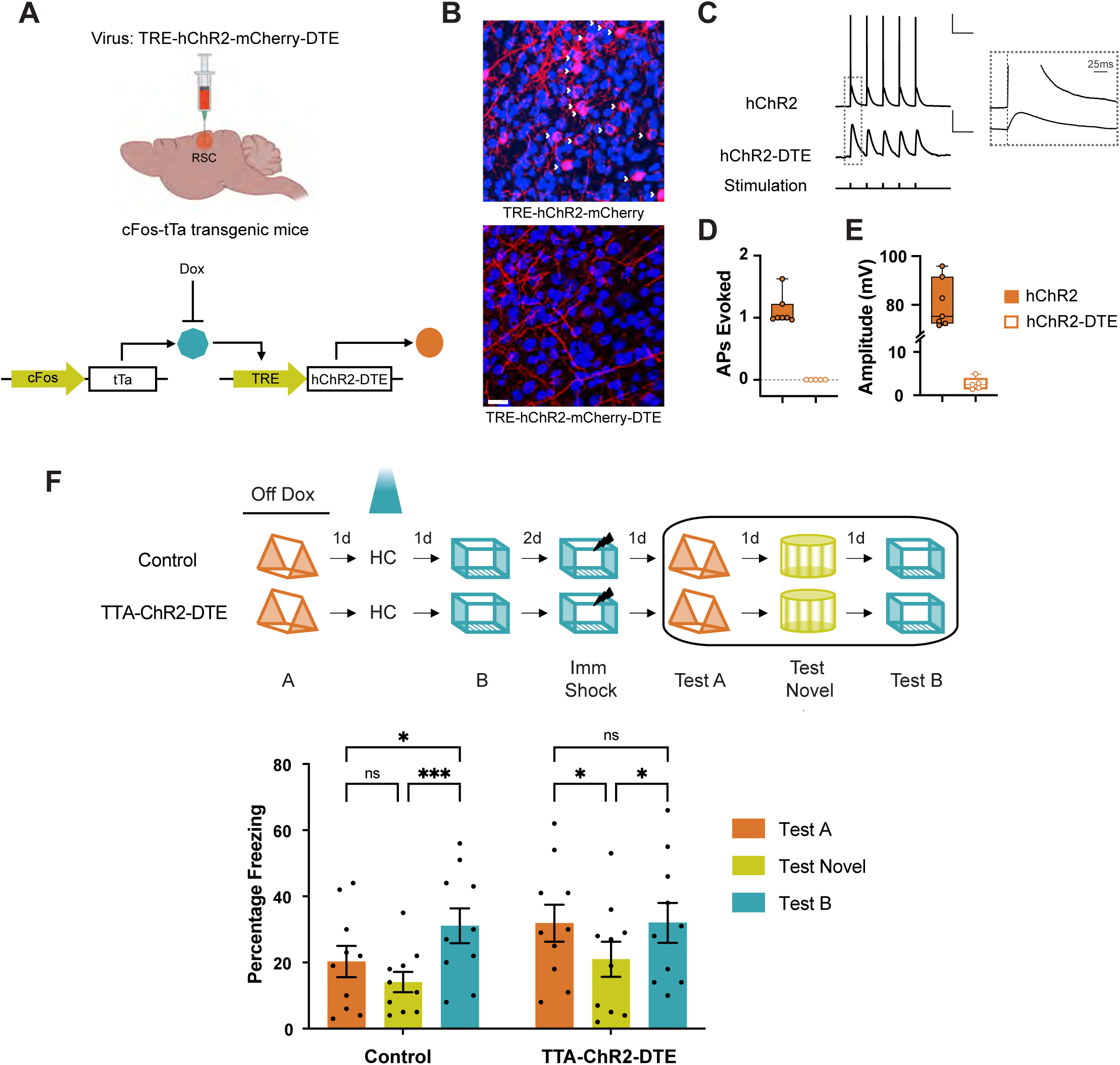
Optogenetic reactivation of RSC dendritic ensembles links contextual memories. (a) The TRE-hChr2-mCherry-DTE virus was designed to optogenetically reactivate previously activated dendritic segments. The key genetic elements include 1) TRE promoter and 2) Dendritic Targeting Element (DTE) at the 5’ end: The DTE element targets mRNA to activated dendrites and has been used to target photoactivatable molecules to activated dendrites/spines (*8*). TRE-hChr2-mCherry-DTE virus was injected into cFos-tTa mice to express Channelrhodopsin in the recently activated dendritic segments of cFos-expressing neurons. (b) Representative RSC images of cFos-tTa mice injected with TRE-hChR2-mCherry-DTE (left) and TRE-hChR2-mCherry (right) showing selective expression of Channelrhodopsin in dendritic segments in the presence of DTE. White arrowheads: somatic expression of hChR2 in the absence of DTE. Scale: 20 µm (c) Whole-cell patch-clamp recordings from RSC neurons of cFos-tTa mice tagged using TRE-Chr2 or TRE-Chr2-DTE constructs. Representative waveforms showing optogenetic stimulation of RSC neurons from TTA-ChR2 and TTA-ChR2-DTE mice results in somatic responses in the form of action potentials and transient depolarizations respectively. Scale, 20mV (top), 1mV (bottom); 250ms. Inset: Magnified view of the first optogenetic stimulation showing response latencies of the stimulus onset. Scale, 25ms. (d) Average number of action potentials elicited and (e) response amplitudes in TTA-ChR2 and TTA-ChR2-DTE mice. (Mann Whitney test, p < 0.005; TTA-ChR2: n = 7 cells, N =3 mice and TTA-ChR2-DTE: n = 5 cells, N =3 mice). Only neurons with detectable responses were plotted. (e) Experimental setup: Top: Mice explored two contexts 2 days apart (a time interval when memories are not linked). The physical contexts presented were counterbalanced to minimize any effect of context similarity. On the day between the 2 context exposures, the dendrites activated during the first context exposure were reactivated optogenetically. Bottom: Reactivation of context A dendrites, on the day between the two context exposures (contexts A and B), results in high freezing in both the previously explored contexts (context A: linked context and context B: shock context) relative to freezing in a novel context. The control mice freeze similarly in context A and novel context, but the freezing in context B (shock context) is higher than freezing in context A or a novel context (n = 10 mice each for Control and cFos-tTa groups; Two-way RM ANOVA, F_time_ (2, 36) = 14.11, P< 0.001; Dunnett’s multiple comparisons test). Data represent mean ± s.e.m. and each data point, * *p* < 0.05, ** *p* < 0.01, **** *p* < 0.0001.

Next, we assessed whether activation of dendritic segments tagged in this manner results in somatic activation. While optogenetic stimulation of RSC neurons from TTA-ChR2 mice resulted in somatic responses in the form of action potentials, the same stimulation only elicits transient small amplitude depolarizations in the TTA-ChR2-DTE mice (Figure 6c-e). Moreover, we also tagged and reactivated RSC dendrites activated during contextual memory formation in a novel context (Figure S6d, e). Unlike our somatic manipulations, tagging and activation of dendritic compartments alone were not sufficient to elicit fear expression in a novel context. Together these data indicate that Channelrhodopsin-mediated dendritic activation using the TTA-ChR2-DTE system has limited effects on the depolarization of somas and therefore fails to elicit an acute behavioral response. Hence, combining the cFos-tTA system with DTE allowed us to study the role of previously active dendrites (while limiting somatic involvement) in memory linking.

Next, we asked whether artificially biasing dendritic allocation, similar to our neuronal manipulations (Figures 2e and S9), is sufficient to link two contextual memories, which would otherwise be independent. We tagged active dendrites during the first context exploration (Context A) and reactivated these dendrites the next day while the mice were in their home cage. One day after this reactivation (i.e., two days after the exploration of Context A), we exposed the mice to another novel context (Context B). As we have shown above, two contexts explored 2 days apart are normally not linked (Figure 2e and S9)^12^. However, similar to the reactivation of Context A neurons (Figure 2e), the reactivation of dendrites first activated in Context A is sufficient to link Context A and Context B (the shock context): freezing in the ‘linked context’ (Context A) was higher than freezing in a ‘novel’ context and similar to freezing in the shock context (Context B). Hence, the reactivation of dendrites tagged during the exploration of one context is sufficient to link that context to another independent context (Figure 6f). These results demonstrate a causal role for RSC dendritic mechanisms in the allocation as well as linking of contextual memories and reveal a novel set of rules that govern how linked, and independent memories are allocated to various dendritic compartments.

### Biophysical dendritic modeling: dendritic plasticity mechanisms are necessary for neuronal overlap and co-recall of memories

Our data thus far suggests that synergism between somatic and dendritic mechanisms sculpts memory allocation within the RSC to regulate the linking of memories. To explore whether linking of memories that are acquired close in time is even possible in the absence of dendritic mechanisms, we adapted a network model of memory allocation^19,51^ and used it to investigate how two independent memories can first become linked in a brain region. The model incorporates somatic as well as dendritic allocation mechanisms that rely on intrinsic excitability modulation (see Methods, Figure 7a, b), as suggested by prior studies^4,8,73^. As with our experimental data, the network model shows that neuronal (Figure 7c), as well as dendritic overlap (Figure 7d), is higher when two memories are acquired close in time (5 hours vs 2 or 7 days apart). Moreover, high dendritic overlap is also characterized by increased synapse clustering (Figure 7e), in line with our spine imaging experiments. Importantly, our model predicts that when linked memories are recalled (i.e., memories acquired 5 hours apart), they maintain higher neuronal overlap indicating co-recall and thus stable linking of these memories (Figure 7f). To dissect the relative contributions of somatic vs. dendritic mechanisms in memory linking, we asked how these neuronal and dendritic overlap measures change in the absence of dendritic allocation and plasticity mechanisms (see Methods). Remarkably, both neuronal and dendritic overlap during encoding is reduced when the model lacks dendritic mechanisms (Figure 7c-e, orange). More importantly, the lack of dendritic mechanisms in the model abolished co-recall or linking of memories, suggesting that dendritic mechanisms are crucial for stable memory linking (Figure 7f, orange: neuronal overlap during recall is the same whether memories are acquired 5 hours, 2 days or 7 days apart).

**Figure 7.**
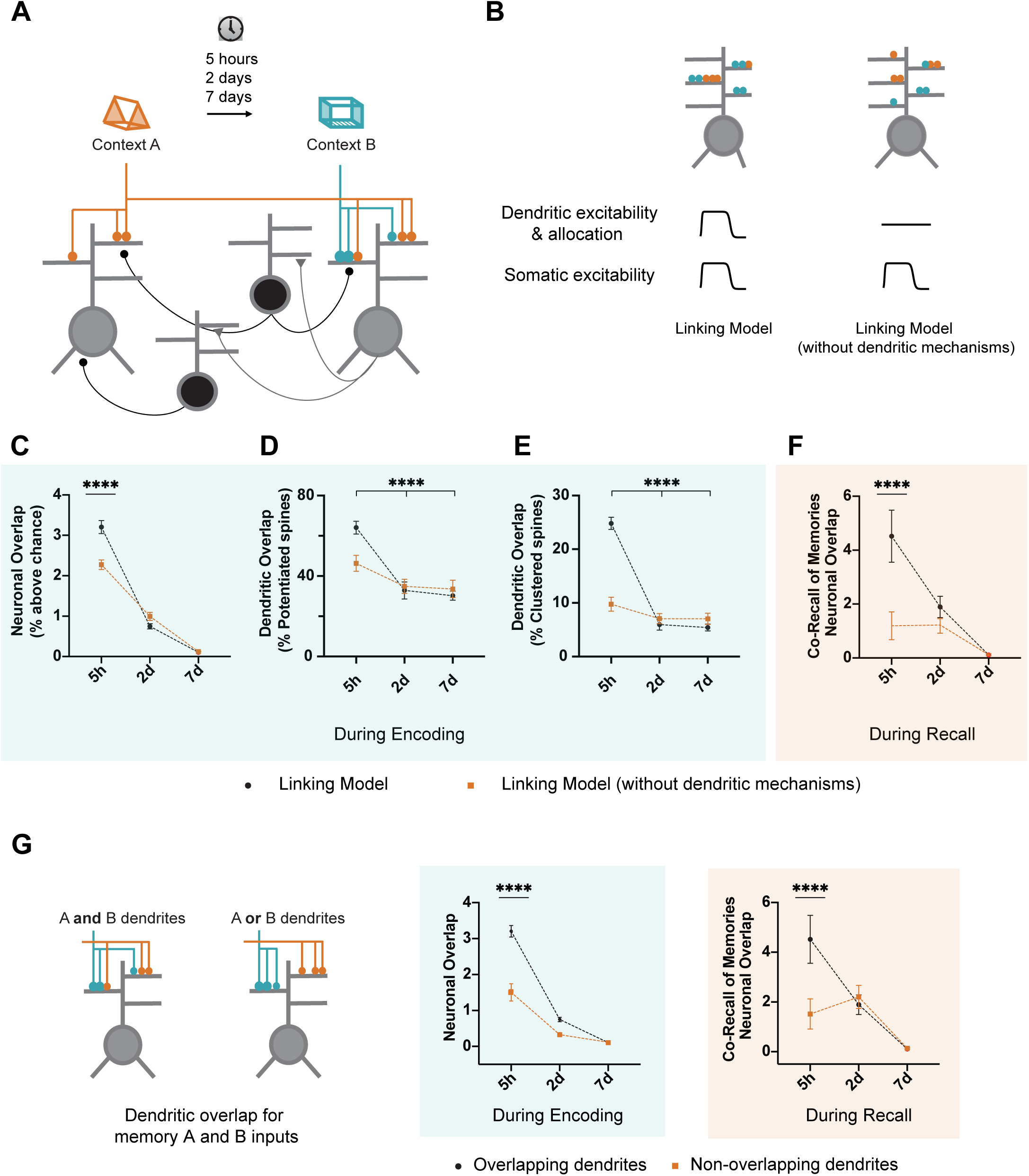
Dendritic mechanisms are necessary for linking memories acquired close in time in a spiking network model. (a) Schematic of the spiking network model. The network consists of 2-layer excitatory neurons (gray) with dendritic subunits, and subpopulations of dendrite-targeting and soma-targeting interneurons (black). See Methods for details. (b) Details of the learning-related plasticity mechanisms within the two network models: The linking model contains somatic and dendritic allocation mechanisms such that memory formation results in increases in somatic and dendritic excitability, and synapses are more likely to be potentiated in the presence of preexisting potentiated synapses on the same dendrite (see Methods). To assess the contribution of dendritic mechanisms to memory linking, learning-related changes in dendritic excitability and probability of synaptic potentiation are eliminated in the linking model without dendritic mechanisms. (c) Neuronal overlap during the encoding of two memories acquired 5 hours, 2 days or 7 days apart. When dendritic mechanisms (dendritic allocation mechanisms and increased dendritic excitability) are removed from the model, neuronal overlap during encoding is reduced when memories are acquired 5 hours (5h) apart. Neuronal overlap represents percentage above chance overlap. (d) Overlap between dendritic branches containing potentiated synapses following two memories acquired 5 hours, 2 days or 7 days apart. When dendritic mechanisms are removed from the model, overlap between dendritic branches with potentiated spines is reduced when memories are acquired 5 hours (5h) apart. (e) Overlap between dendritic branches containing newly added clustered spines following two memories acquired 5 hours, 2 days or 7 days apart. When dendritic mechanisms are removed from the model, overlap between dendritic branches with newly added clustered spines is reduced when memories are acquired 5 hours (5h) apart. (f) Co-recall of two memories as measured by neuronal overlap during recall. When dendritic mechanisms are removed from the model, co-recall of two memories (as measured by neuronal overlap during recall) is reduced when memories are acquired 5 hours (5h) apart. In addition, when dendritic mechanisms are removed from the model, neuronal overlap during recall is not different whether memories are acquired 5 hours, 2 days or 7 days apart indicating that dendritic mechanisms are necessary for linking of memories acquired close in time. Two-way RM ANOVA. Dunnett’s multiple comparisons for Figure 7c-e: linking model and linking model (without dendritic mechanisms): 5h vs 2d and 5h vs 7d, all *p* values < 0.0001. For Figure 7f, linking model: 5h vs 2d and 5h vs 7d, all *p* values < 0.0001 but for linking model (without dendritic mechanisms): 5h vs 2d and 5h vs 7d, all *p* values = non significant. Sidak’s multiple comparisons for Figure 7c-e, overlaps at 5h for linking model vs linking model (without dendritic mechanisms): all *p* values < 0.0001. For simplicity only comparisons within the linking model without dendritic mechanisms are presented (5h vs 2d or 5h vs 7d, **** *p* < 0.0001). (g) Dendritic overlap allows somatic overlap and co-recall of memories. Inputs representing context A and B impinge on overlapping or completely separate dendrites (dendritic overlap is eliminated). During encoding and recall, neuronal overlap is reduced between groups at 5h but not 2d and 7d time intervals (Sidak’s posthoc, p < 0.0001). When memories are encoded by non-overlapping dendrites, neuronal overlap is similar between 5h, 2d and 7d group (for overlapping dendrites group, Dunnett’s posthoc, p < 0.0001). Neuronal overlap represents percentage above chance overlap (see Methods). Data represent mean ± s.e.m. of 10 simulation trials.

To assess the importance of converging synaptic input onto the same dendritic compartments for memory linking, we modeled synaptic inputs representing the two contexts (A and B) on separate (exclusively non-overlapping) dendritic branches. The model predicts impaired neuronal overlap (during encoding and recall) when two memories encoded 5 hours apart recruit non-overlapping dendritic populations, suggesting that to effectively link separate inputs within a neuron, these inputs need to overlap onto the same dendritic compartments (Figure 7g). Together, our data indicate that dendritic allocation mechanisms may be necessary (Figure 7) and sufficient (Figure 6) for linking memories acquired close in time.

## DISCUSSION

Our findings demonstrate that localized dendritic mechanisms play a causal role in mediating neuronal ensemble overlap and thus, linking of contextual memories. We demonstrate that in addition to neuronal ensemble overlap, local dendritic rules further sculpt the allocation of memories to dendritic segments, such that temporally proximate (i.e., linked) memories are likely to be allocated to the same dendritic segments, while temporally distant (i.e., independent) memories are not. We leveraged activity-dependent targeting of dendritic segments to demonstrate that biasing memory allocation to the same dendritic segments is sufficient to link these memories. Accordingly, computational modeling supports the key role of dendritic mechanisms in memory linking. Altogether, the findings presented here demonstrate that localized dendritic mechanisms are critical for linking memories. Furthermore, since RSC is one of the first brain regions to demonstrate Alzheimer’s-related dysfunction^74,75^, and memory linking is affected early during aging^12^, our findings have clinical implications for understanding age-related cognitive decline and associated pathological conditions.

### RSC neuronal ensemble activation during encoding of linked memories

Within the RSC, neuronal ensemble activation encodes contextual and spatial information^24,25,55,76,77^. We asked whether RSC representations of two contextual memories acquired close in time (5h apart), such that they are behaviorally linked, are more similar than those of two memories acquired a week apart (independent memories). We discovered that linked memories activated an overlapping neuronal ensemble in the RSC (Figure 1e). Furthermore, the inferred activity and co-activity dynamics of these RSC ensembles were also similar for linked, but not independent memories (Figure 1g-i). Accordingly, a classifier showed lower accuracy for decoding distinct contexts seen closer in time (i.e., linked contexts) compared to contexts seen further apart in time (i.e., independent contexts). Our data are consistent with the important role of RSC neural dynamics in contextual memory encoding and demonstrate that the neuronal representations of two linked contexts encoded close in time are more similar than the neuronal representations of independent contexts. In addition, these data indicate that linked memories are encoded by overlapping neuronal ensembles in multiple brain regions, including hippocampal CA1^12,48^ and RSC (Figure 1).

### RSC neuronal mechanisms underlie the linking of contextual memories

Several studies have demonstrated that plasticity within the RSC is necessary for contextual fear expression^76,78^, and RSC-associated brain circuits have been described that underlie the encoding of single contextual fear memories^45–47^. Consistent with previous findings^24^, we demonstrate that optogenetic reactivation of RSC ensembles (as captured by the TetTag system) is sufficient for contextual fear expression (Figure 2c). While similar experiments in the hippocampus lead to fear expression specifically during the stimulation (Light-On epochs), we found that RSC ensemble reactivation results in a sustained increase in freezing during the post-stimulation period. These differences could be a result of experimental conditions (context size and similarity etc.)^79^ which can affect the nature of memory expression upon optogenetic stimulation. It is just as likely that RSC ensemble reactivation which by itself is sufficient to elicit fear expression in the absence of hippocampal activity elicits a distinct pattern of activation in downstream target regions resulting in a sustained fear expression^24^.

Importantly, we show that optogenetic reactivation of the RSC ensemble underlying a linked neutral memory is sufficient to induce fear expression associated with a second fearful memory (Figure 2d). Thus, the activation of a neural ensemble underlying one of the linked memories is sufficient for the co-recall of another linked memory. Moreover, these effects on contextually linked memories can be mediated by RSC ensembles alone. These data demonstrate the critical role of the RSC in mediating the expression of linked contextual memories.

Linking of contextual memories is robust when these memories are acquired within the same day, but with intervals of two or more days, these memories are behaviorally independent of one another (Figures 2e and S9)^48^. Here, we demonstrate that optogenetic or chemogenetic manipulation of RSC ensemble overlap alone is sufficient to link two otherwise independent memories (Figure 2e and S9). These data are consistent with amygdala-dependent linking of aversive memories, where the retrieval of a previously encoded memory resets the time window for linking a new memory^14^. Similarly, persistent reactivation of neural ensembles underlying two aversive amygdala-dependent memories by co-recall links them behaviorally^13^. It is likely that optogenetic activation of RSC ensembles underlying the first memory in the home cage (Figure 2c) results in recall of this memory, which is likely to re-engage the same plasticity mechanisms that were important for encoding the original contextual memory, thereby extending the window for memory linking. Using a similar but mechanistically different strategy, we chemogenetically increased the excitability of the same sparse RSC ensemble prior to two contextual memory episodes (Figure S9). Similar strategies have been used to bias memory allocation to the underlying ensemble^12,14,49^. Forcing neuronal overlap within RSC ensembles in this way was sufficient to link two independent contextual memories (Figure S9). In addition, we show that transiently manipulating the activity of RSC neurons before a single context exposure does not affect memory linking (Figure S9). Finally, we found that activating a sparse but randomly labeled neuronal ensemble in a similar manner did not result in memory linking (Figure S8) demonstrating that effects are specific to the ensembles representing a novel context. Together, our data demonstrate that neuronal overlap within the RSC ensemble alone is sufficient to link distinct contextual memories (Figures 2 and S9), and that this is a key mechanism for linking contextual memories under various behavioral conditions.

We find that neuronal overlap in the RSC, similar to overlap in the CA1 ensembles, can affect the linking of contextual memories. Activity within the hippocampus as well as the RSC is important for contextual as well as spatial tasks^24,25,44,78,80–82^. For example, activity and plasticity within CA1 as well as RSC ensembles during contextual fear memory formation affects retrieval of contextual fear memories^78,83^. Similarly, the reactivation of RSC^24^, CA1^17^, or dentate gyrus^44^ ensembles tagged during context fear memory formation is sufficient to induce fear expression in a novel neutral context. While the precise role and the interaction of hippocampal and retrosplenial sub-regions in contextual memory processing is not well understood, optogenetically reactivation of tagged RSC ensembles can result in fear expression even when hippocampal activity is inhibited^24^. These data indicate that both hippocampus and RSC support contextual memory formation and RSC ensembles can support fear expression independently. Subiculum to RSC projections arising from vGlut1 and vGlut2-expressing pyramidal neurons play a differential role in recent and remote contextual memory formation^46^. These data and RSC’s well-established projections to the important nodes in the fear circuit^84^ support the hypothesis that RSC may be downstream of the hippocampus in the contextual fear memory circuit and plays an important role in information processing within this circuit. RSC activity is especially critical in more complicated associative memory paradigms^85,86^, tasks where discrimination between similar stimuli is required^87^ as well as generalization^88^ and remote memory recall^46^. The precise contribution of hippocampal and RSC processing to contextual memory formation, as well as the linking of contextual memories, is unknown but important to develop a clearer understanding of these phenomena and will be an exciting new area of research.

### Localized dendritic plasticity mechanisms mediate the encoding of temporally proximate memories

Experience-dependent localized dendritic plasticity has been assessed in *ex vivo* settings^4,6–8,23,89^. However, it was unclear whether new learning induces compartmentalized dendritic plasticity. Moreover, the function of such localized dendritic plasticity in memory processes is unknown. We show that linked contextual memories result in the activation of overlapping dendritic branches within RSC apical dendrites (Figure 3). The inferred firing rate, a measure in part dependent on the excitability of dendritic segments, was more correlated between segments reactivated following the encoding of linked, but not independent memories. In addition, we demonstrated that following the formation of linked memories, structural synaptic plasticity (addition of single spines as well as spine clusters) is also biased to overlapping dendritic segments (Figures 4 and 5). Although our dendritic calcium imaging results (Figure 3) did not conclusively rule out a neuron-wide pan-dendritic reactivation, we believe this possibility is unlikely given low-level dendritic overlap and the wealth of evidence that learning-related plasticity is input-specific and only observable on some but not all dendritic/synaptic loci within a neuron^21,90–92^. Our finding that linked contextual memories are co-allocated to overlapping dendritic branches is consistent with the hypothesis that experience-dependent localized dendritic plasticity is a metaplasticity mechanism that influences future plasticity on these dendritic branches. Indeed, our computational model predicts that a localized increase in dendritic excitability, and the associated facilitation of structural plasticity on these dendritic segments, are necessary for neuronal overlap during recall (i.e., a measure of co-recall in our biophysical model, Figure 7).

We targeted apical dendrites from layer V RSC neurons as we have previously shown that these dendritic compartments undergo clustered plasticity (i.e., spine addition) following contextual memory formation. Such clustered spine addition is correlated with improved learning, and spines added in clusters are more stable^19^. Manipulations that increase clustered spine addition on these dendritic branches also facilitate contextual memory performance^19^. Apical dendrites from layer V RSC neurons receive projections from various thalamic nuclei^45,93^ including anterodorsal and anteroventral subdivisions of the anterior thalamic nuclei (ATN)^45,47^ which regulate contextual memory encoding and specificity, respectively^47^. Apical dendrites from layer V RSC neurons also receive long-range projections from a sparse population of GABAergic CA1 neurons that co-regulate RSC activity along with ATN inputs^45^. It is currently unclear how these various inputs to RSC dendrites interact to facilitate clustered synaptic plasticity as well as the linking of contextual memories. Since, the inactivation of hippocampal as well as thalamic inputs to the RSC impairs fear expression^45–47^, these inputs likely also regulate the linking of contextual information within the RSC.

In addition to the circuit mechanisms that underlie memory linking, the intracellular mechanisms that mediate clustered plasticity and reactivation of the same dendritic compartments are also unknown, but biophysical computational models (Figure 7) should provide testable hypotheses for future investigations. Ex vivo investigations of localized dendritic plasticity have revealed several underlying mechanisms. These include the plasticity of dendritic spikes^8,56^, changes in ion channel function^4^, signaling pathways^7^ as well as protein synthesis-dependent mechanisms^6^. For example, an increase in local dendritic excitability could facilitate dendritic depolarization, which in turn would promote the addition of spine clusters to the same dendritic segment during memory linking.

To demonstrate that the dendritic overlap we discovered underlies the linking of contextual memories within RSC ensembles, we modified the TetTag approach to tag and manipulate specific dendritic segments within tagged RSC cells (Figures 6, S6 and S17). Our tagging and manipulation approach involved the addition of a DTE sequence to Channelrhodopsin to restrict Channelrhodopsin expression to recently active dendritic segments. DTE had previously been used to restrict photoactivatable molecules to dendritic segments in an activity-dependent manner^18,65,66^.

Similar to the reactivation of RSC neuronal ensembles, the reactivation of RSC dendritic ensembles alone allowed us to extend the temporal window during which contextual memories are linked. Importantly, optogenetic activation with TTA-Chr2 (Figure 2e) results in the activation of somas as well as all entire dendritic tree, whereas TTA-Chr2-DTE restricts activation to a smaller subset of dendrites that were previously active (Figure 6f). These results demonstrate that dendritic ensemble overlap plays a key role in memory linking.

In the current study we use an established behavioral paradigm to demonstrate that neuronal and dendritic mechanisms in the RSC regulate the linking of two memories experienced five hours apart. We assume that discrete memories that share common elements can be linked and organized together across various dimensions ^11^ including temporal proximity (such as in our current study), contextual similarity^94^ and other common elements among memories^95^. Organizing memories based on these common elements can facilitate various higher-order cognitive processes such as inference, causal reasoning, etc. Therefore, given the obvious evolutionary advantage of linking memories that share common elements to the organism’s success, we hypothesize that the mechanisms that allow memory organization are tightly regulated and depend on the conserved features across brain regions. Changes in intrinsic neuronal and dendritic excitability are likely to mediate the linking of memories across various brain regions and various dimensions. Furthermore, such intrinsic processes are ideally positioned to use neuromodulatory systems to determine which memories are linked and which remain independent based on their salience^96^.

Finally, within the dendritic arbor, inputs from distinct pathways are organized on distinct dendritic domains ^97^. This organization along with our current findings suggests that only inputs impinging on the same dendritic compartments might be linked while other inputs remain independent (Figure 7). Therefore, dendritic plasticity mechanisms may facilitate the linking of similar memories that share similar modalities (i.e. two contextual memories) that share similar inputs that synapse onto overlapping dendritic segments. On the other hand, memories encoded by inputs synapsing onto non-overlapping dendritic domains do not benefit from dendritic plasticity mechanisms and thus remain unlinked.

Understanding the roles of branch-specific plasticity mechanisms in differentially modulating inputs within a neuron, and by extension, the larger neuronal circuit, is important to understand information processing within a distributed circuit. Importantly, aberrant dendritic and synaptic plasticity mechanisms are also a hallmark of many pathological states, making the findings presented here clinically relevant^98,99^.

## Supplementary figures

**Figure S1.**
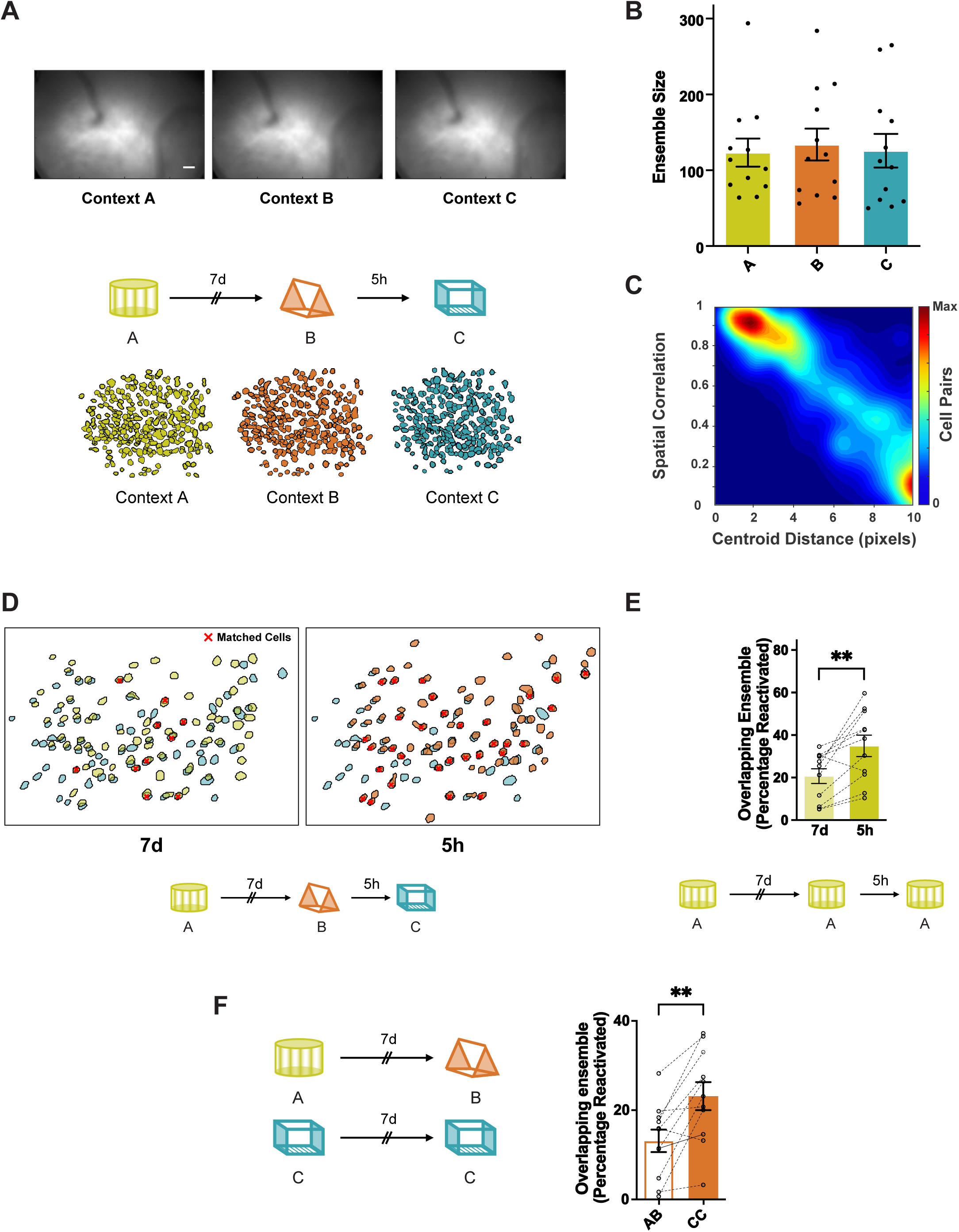
Stability of imaging and neuronal registration across days. (a) Top: Images of mean fluorescence from each session from a representative mouse. Scale: 50 µm. These images from each session were cross-registered with each other (see Methods for details). Bottom: Description of imaging paradigm and RSC ensemble segmented from a mouse. (b) RSC ensemble size captured by Miniscope imaging remains stable across hours and days when different contexts are imaged (4599 putative RSC neurons, 132.9 ± 11.6 neurons per session, One-way repeated measures ANOVA, F (1.08, 11.9) = 0.52, *p* = 0.5, n = 12 mice per group). Please note that although the size of these ensembles remains unchanged, the neurons participating in these ensembles may change. Data represent mean ± s.e.m. and each data point. (c) Spatial correlation and centroid distance were calculated for all cell pairs from all mice. Ensemble overlap using a range of criteria from spatial correlation ≥ 0.6-0.95 and centroid distance ≤ 3-9 pixels is shown in Supplementary Table 1. (d) Example cross-registration of neurons in a mouse from sessions 7 days and 5 hours apart. Red cross indicates matched cells. Cross-registration criteria: spatial correlation = 0.9 and centroid distance = 4 pixels. (e) Representational drift over a week: Mice were exposed to the same context (AAA) five hours or seven days apart. RSC neuronal ensembles display greater overlap when mice experience the same context 5 hours vs 7 days apart. (n = 12 mice per group; paired t-test, t = 3.7, *p* < 0.005). Data represent mean ± s.e.m. (f) Neuronal ensemble stability over a week: Mice were exposed to two different (AB) or the same context (CC) seven days apart. All context presentations were counterbalanced. RSC neuronal ensembles display greater overlap when mice experience the same context 7 days apart vs distinct contexts. (n = 11 mice per group, paired t-test, t = 4.07, p < 0.01). Data represent mean ± s.e.m. and each data point.

**Figure S2.**
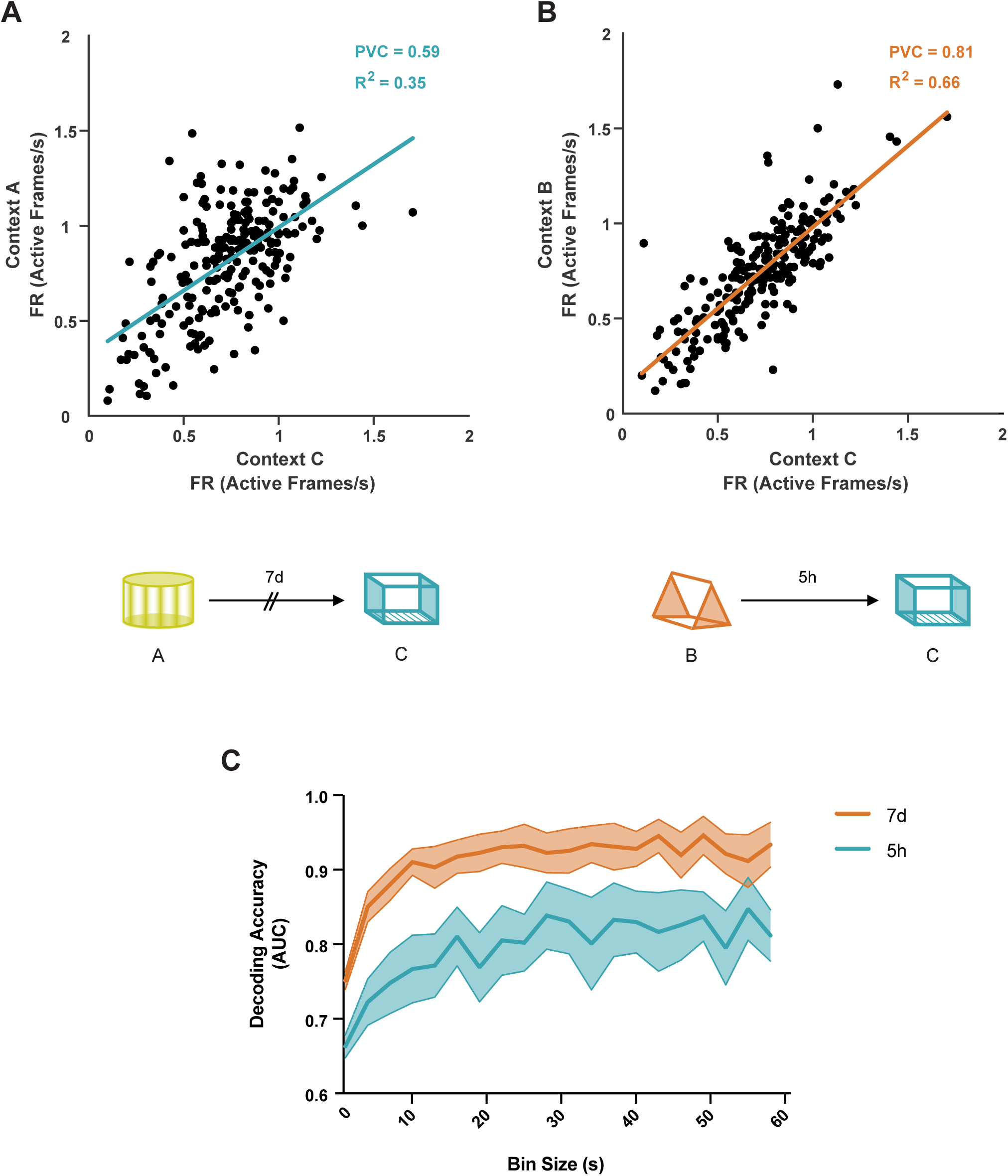
Neuronal activity is more correlated when two contexts are explored closer in time. (a, b) Scatter plot of the firing rate of all neurons from one mouse in context A (a, 7 days apart) and context B (b, 5 hours apart) as a function of firing rate in context C highlights that neuronal firing rate is maintained when two contexts are explored close in time. Lines represent least-squares linear regression. (c) Naïve Bayes (NB) classifier is more accurate at distinguishing imaging sessions recorded 7 days vs 5 hours apart irrespective of bin size. AUC (area under the curve) for the binary NB classification between sessions recorded 7 days (7d) apart or 5 hours (5h) apart using neuronal activity indicates that neuronal activity can be used to distinguish between contexts explored 7 days apart more accurately than contexts explored 5 hours apart. Spike probabilities were binned for non-overlapping intervals ranging from 0.5 to 60 seconds (step size 0.5s; Two-way Repeated Measures ANOVA for AUC by bin size; F_Group_ (1, 16) = 6.2, *p* < 0.05, n = 9 mice per group). Data represent mean ± s.e.m. Chance Levels performance of the AUC = 0.5.

**Figure S3.**
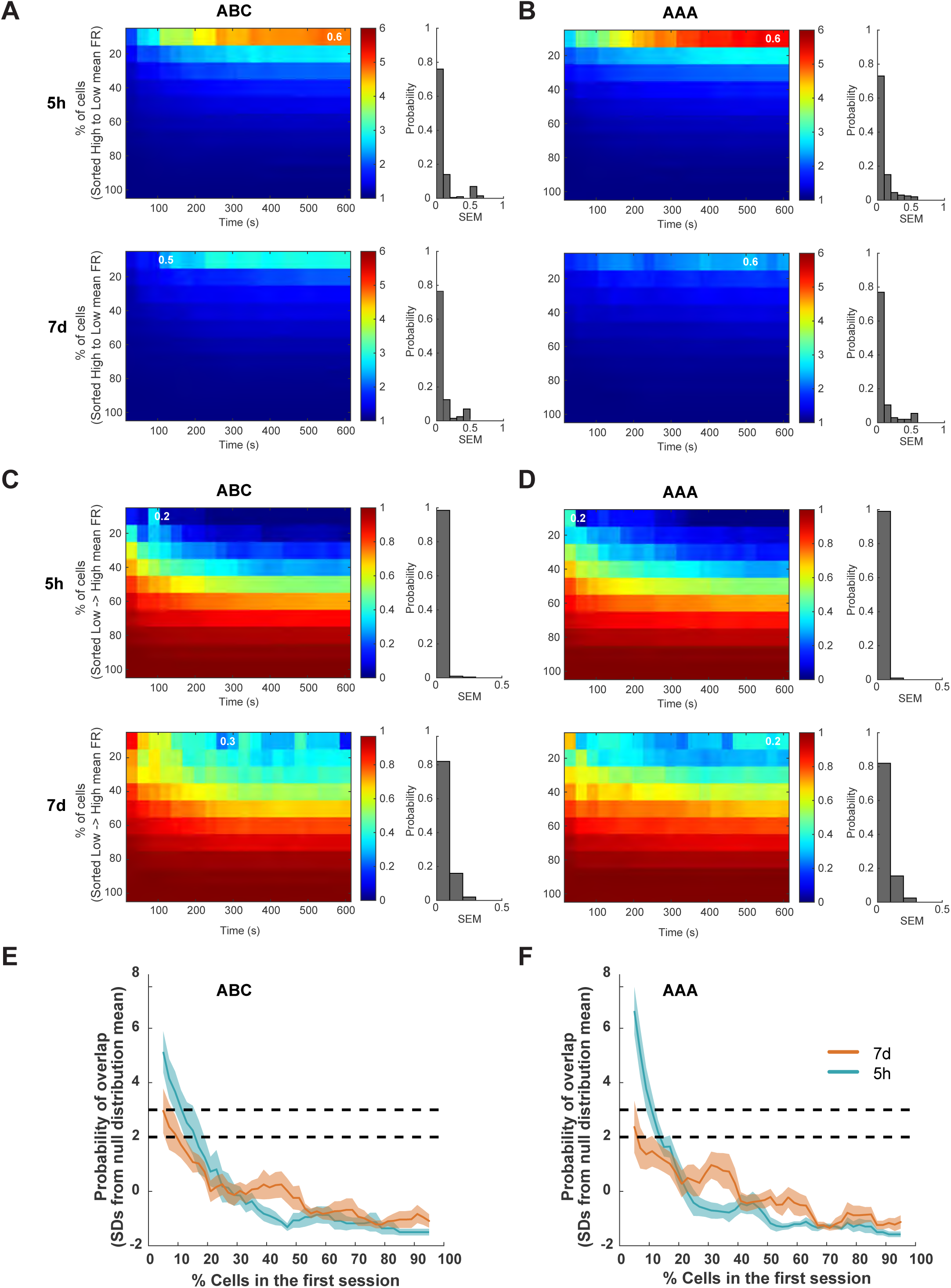
Neuronal activity is correlated with the probability of neuronal overlap. (a) The top 10% high-activity cells in context B are very likely to be the top 10% high activity cells in context C 5h later. Probability of overlap between high activity cells in context A (7d) or context B (5h) and high activity cells in context C. Left: Probability of overlap between subsets of cells with different levels of activity during previous context exploration session (A or B) across time in C. Color bars refer to normalized probabilities (chance = 1). Cumulative values were used for x and y axis (e.g., for x-axis, 300s means 0–300 s; for y-axis, 40 refers to the neurons within the top 40% of high activity). Values represent average across mice. Right: the distribution of SEM across mice for the figures on the left. Numbers (in the probability of overlap figures) represent the maximum SEM from each plot. (b) Similar to figure (a) but comparisons are performed between the same context experienced 5 hours or 7 days apart. (c,d) Similar to figures (a) and (b) respectively but the probability of overlap between low activity cells in contexts experienced 7 days or 5 hours before and high activity cells in the third context is presented under ABC (c) and AAA (d) conditions. (e) Cells were sorted from high to low activity in contexts A or B with a 10% sliding window and 2% steps. Plots show the probability of overlap between subsets of cells (10% ensemble size) from context A (7d) or B (5h) and the top 10% high FR cells from context C. The probability values were z-scored with respect to a null distribution created by randomly subsampling 10% of cells from contexts A or B 10,000x (i.e., results are represented as standard deviation (SD) from the mean of the null distribution). The 2SD and 3SD thresholds are labeled with a dashed line. (f) Same comparison as (e) but for the AAA condition. (ABC: n = 9, AAA: n = 9 mice).

**Figure S4.**
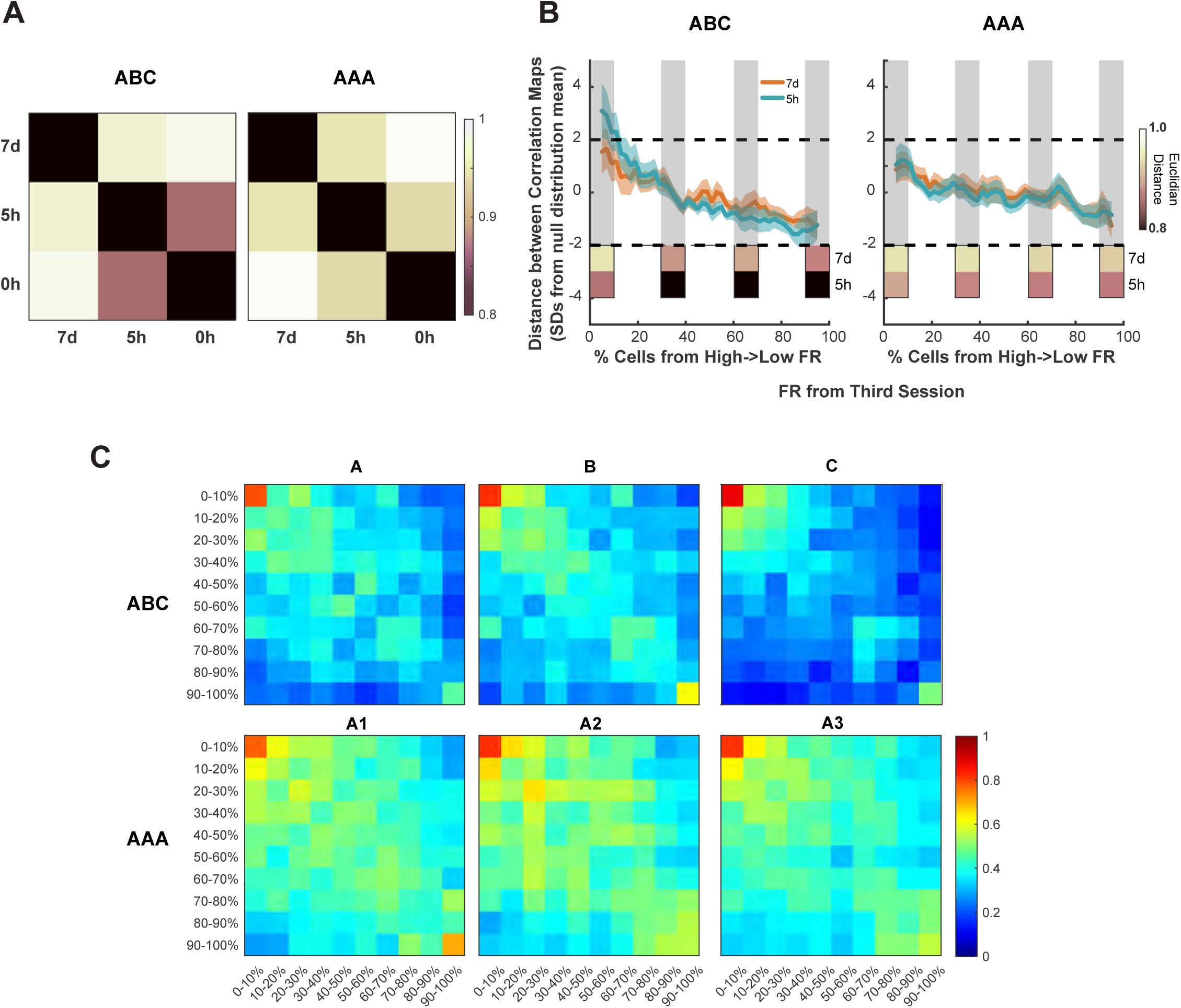
High and low firing rate neurons make differential contributions to representational similarity to regulate memory linking and discrimination. (a) Euclidean distance between correlation maps from different sessions. For each animal, the Euclidean distance was calculated for all possible combinations to create a Euclidean distance map. For each map, all the distances were normalized by the maximum distance. Normalized Euclidean distance maps were then averaged across animals to produce the plots (ABC, right, n = 9; AAA, left, n = 9). Note that the Euclidean distance between correlation maps in the 5h interval is lower than for the 7d interval for the ABC (right) or the AAA (left) contexts conditions. (b) Cells were sorted from high to low activity in context C (x-axis) with a 10% sliding window and 2% steps. Correlation maps were calculated by excluding 10% of cells belonging to each of these sliding windows and the Euclidean distance (y-axis) between contexts explored 7d or 5h apart under ABC (right) or AAA (left) condition was calculated. The Euclidean distance values were normalized with respect to a null distribution created by randomly subsampling 10% of cells from context C 10,000x (i.e., results are represented as standard deviation (SD) from the mean of the null distribution). The 2SD threshold is labeled with a dashed line. Plots on the bottom of each image show the average Euclidean Distance across animals for the 7d and 5h intervals when the following groups of cells are excluded: 0-10%, 30-40%, 60-70%, and 90-100%. Note that the Euclidean distance for the 5h interval is always lower than for the 7d interval. For AAA condition, the exclusion of any batch of 10% cells does not significantly affect the Euclidean distance. However, for the 5h interval in ABC condition, the top 10% FR cells, when excluded, significantly change the Euclidean Distance. Therefore, the top 10% FR cells are critical for the similarity between correlation maps when different contexts are explored but the contribution of these top 10% FR cells is not significant when the same context is explored at the same time intervals. (c) Plots show the average normalized correlation maps across animals during the exploration of contexts in ABC (top) or AAA (bottom) conditions. For each animal, cells were sorted from high to low firing rate (based on the last context exploration). The neuronal population was then split into 10% non-overlapping groups. Average Pearson correlation between groups was calculated. A correlation map of the average correlation between groups was constructed and normalized to the maximum value of average correlation for each animal. Plots show the average of these normalized correlation maps across all animals. For all conditions and sessions, the top FR cells have the highest correlation values. Importantly, for the AAA condition, all correlation maps are quite similar despite the session. However, for the ABC condition, the maps show larger differences and the similarity between correlation values between contexts B and C, which are 5h apart, seem to be higher for the high FR cells, as shown previously.

**Figure S5.**
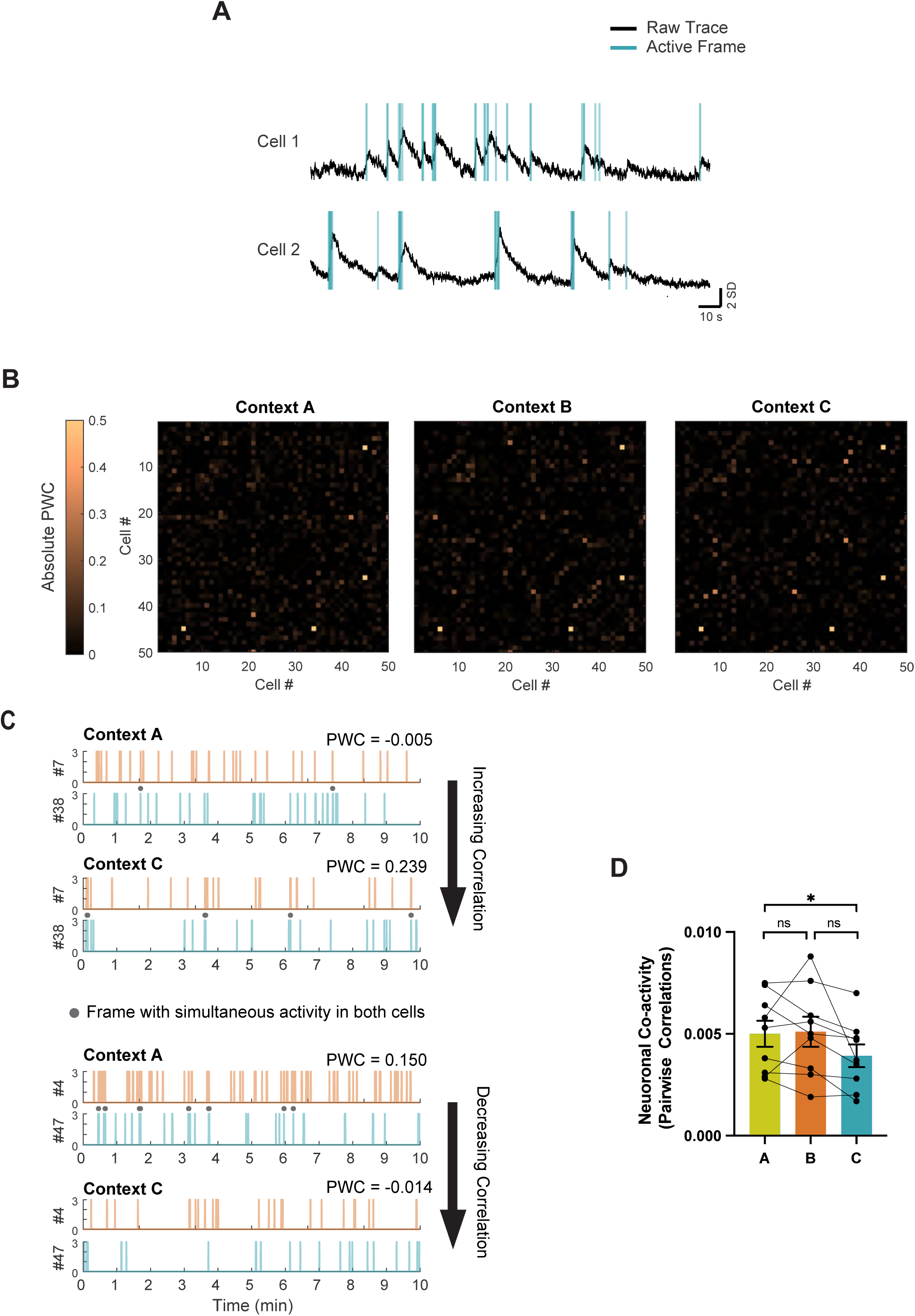
Coactivity among RSC neurons during multiple contextual exposures. (a) Example of two co-active neurons: For each imaging session, deconvolved neuronal activity was binned into 100ms periods, and the Pairwise correlation (PWC: Pearson correlation of the number of active frames) for each neuron pair was calculated. Black: raw calcium trace and Blue: frames identified as active following deconvolution of the calcium trace. (b) Pairwise Correlation (PWC) map for each context for 50 cell pairs from one mouse. Absolute Pearson’s correlation coefficients were plotted. (c) Top Panel: Example of a neuronal pair with increasing PWC between imaging sessions: Cell pair (7 and 38: arbitrary cell IDs) display more coactivity (number of simultaneous active frames) during Context C exploration than Context A exploration. Bottom Panel: Example of a neuronal pair with decreasing PWC between imaging sessions: Cell pair (4 and 47: arbitrary cell IDs) display less coactivity during Context C exploration than Context A exploration. (d) Average PWCs across the three context exposures display a small decrease (One-way repeated measure ANOVA; F (1.6, 12.9) = 4.05, *p* = 0.05; Tukey’s post-hoc test, n = 9 mice per group). Data represent mean ± s.e.m. and each data point, * *p* < 0.05.

**Figure S6.**
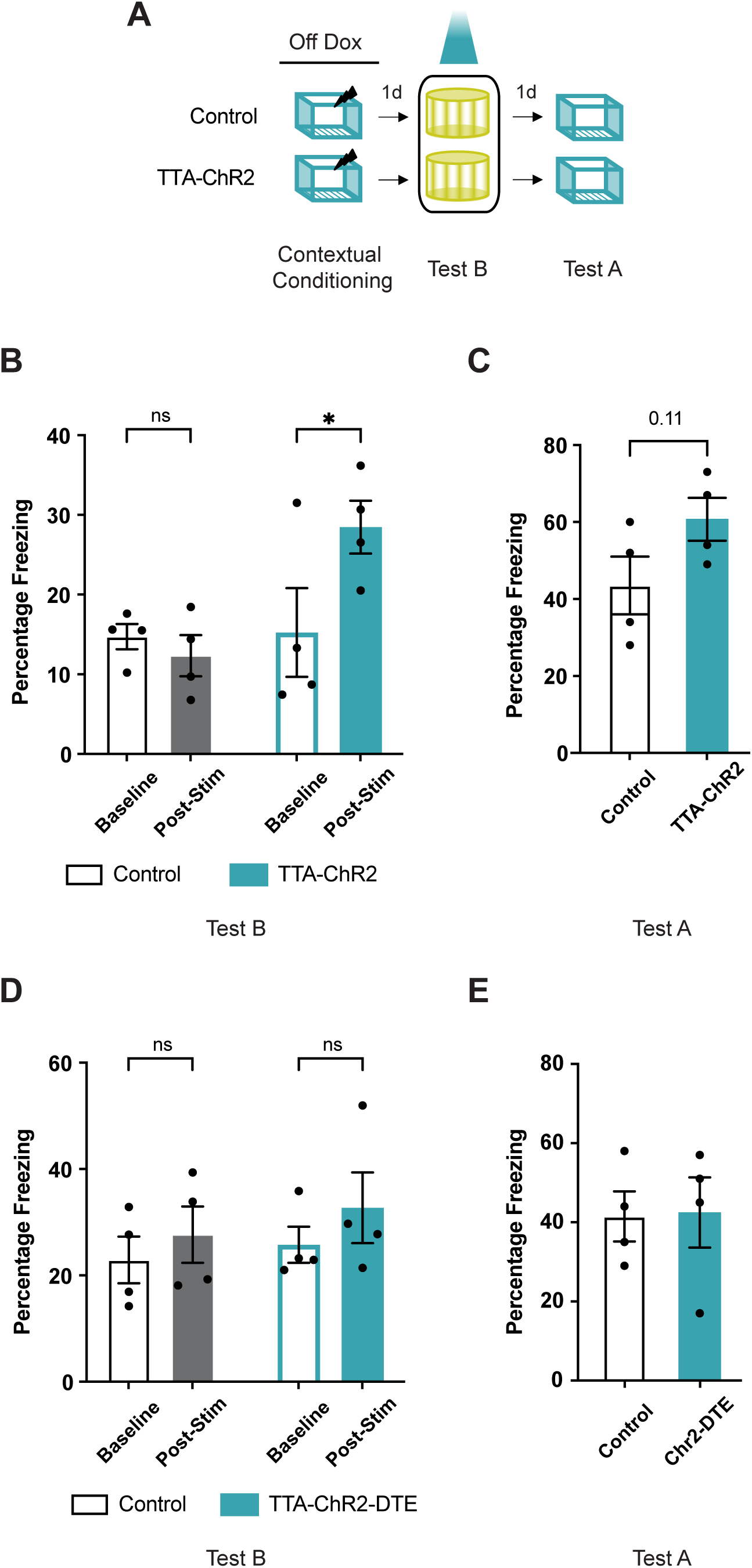
RSC neuronal ensemble (and not dendritic ensemble reactivation) following contextual fear conditioning results in fear expression. (a) Experimental setup: cFos-tTa (TTA-Chr2) mice and their wildtype littermates (Control) underwent bilateral viral injections (TRE-ChR2-mCherry) and optic cannula implants. Mice were taken off doxycycline chow (three days before contextual fear conditioning in context A: 2 footshocks, 2s, 0.7mA) to allow c-fos promoter-driven tTA and Channelrhodopsin (ChR2) expression. Following contextual fear conditioning, mice were tested in a novel context (Test B) while the previously tagged neurons were activated. The following day mice were retested without any optogenetic manipulation in the training context (Test A). (b) During Test B, freezing during the two post-stimulation conditions (with laser and without laser stimulation) was not different. Therefore, freezing during this period is presented together as post-stimulation freezing (Two-way RM ANOVA, group X time interaction, F (1, 6) = 11.93, *p* < 0.01, Sidak’s post hoc tests). (c) During Test A, the TTA-Chr2 mice display comparable freezing to the control mice (t=1.85, df=6, *p* = 0.11). (d) Reactivation of previously activated dendrites is not sufficient for fear memory expression: Experimental set up is the same as (a), but animals were injected with TRE-hChR2-mCherry-DTE or TRE-mCherry-DTE virus in the RSC to reactivate dendritic segments active during contextual fear conditioning. Both groups display similar freezing during baseline and post-stimulation epochs while testing in a novel context (Two-way RM ANOVA, group X time interaction, F (1, 6) = 0.26, *p* = 0.6, Sidak’s post hoc tests). (e) Both groups (injected with TRE-hChR2-mCherry-DTE or TRE-mCherry-DTE virus in the RSC) display similar freezing during test in Context A (t=0.09, df=6, *p* = 0.9). Data represent mean ± s.e.m. and each data point, * *p* < 0.05, ns = not significant.

**Figure S7.**
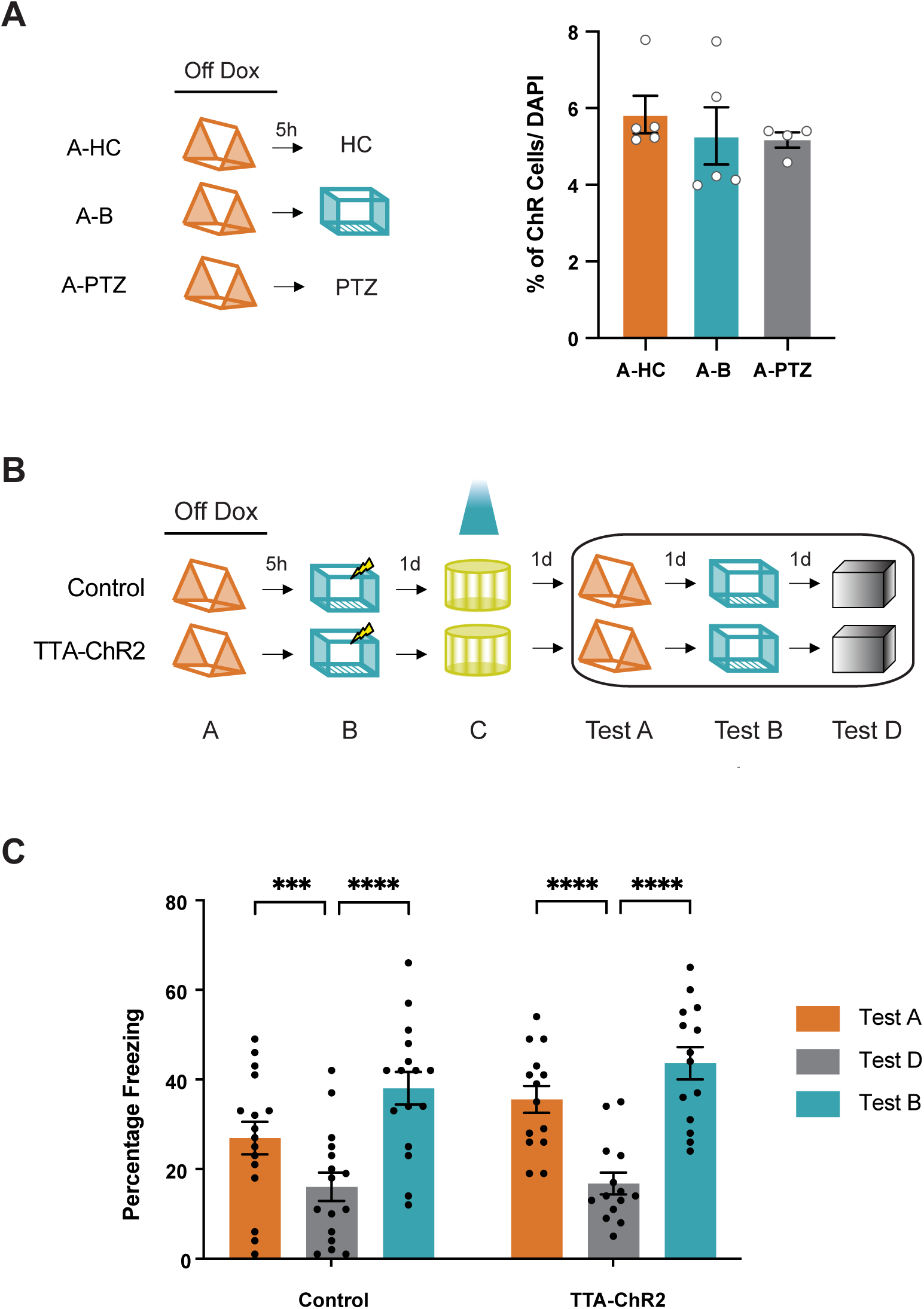
Fear expression due to reactivation of the linked memory ensemble is not driven by changes in the strength of memory linking or tagging of cells outside of the first behavioral episode. (a) Five hours after context exposure, the tagging window is closed by doxycycline. RSC ensemble activated during the exploration of the first context (context A) was tagged. The tagging window was closed by placing the mice on a high concentration of doxycycline chow (200mg/kg) as well as a doxycycline injection (i.p., 50 μg/ gram of body weight) two hours post exposure to context A. Five hours following the context exposure mice were either left in home cage (A-HC), explored another context (A-B) or received an injection of pentylenetetrazole (PTZ, 30mg/kg, A-PTZ). Animals were perfused and brains collected 24 hours post-tagging (same timepoint as optogenetic activation for experiment in Figure 2d). No differences in Channelrhodopsin/mCherry expression were observed between groups (One-way ANOVA, F (2, 11) = 0.4, *p* = 0.7; Dunnett’s multiple comparisons test, ns). (b) Following optogenetic activation of the first memory ensemble (Figure 2d), mice were tested in the linked context (Context A), shock context (Context B), and in another novel context. (c) Both groups of mice display robust memory linking, such that the freezing in both context A (linked) and context B (shock) is higher than freezing in a novel context (Two-way RM ANOVA, F_time_ (1.9, 54) = 86.9, *p* < 0.0001; Dunnett’s multiple comparisons test). Data represent mean ± s.e.m. and each data point, *** *p* < 0.001, **** *p* < 0.0001.

**Figure S8.**
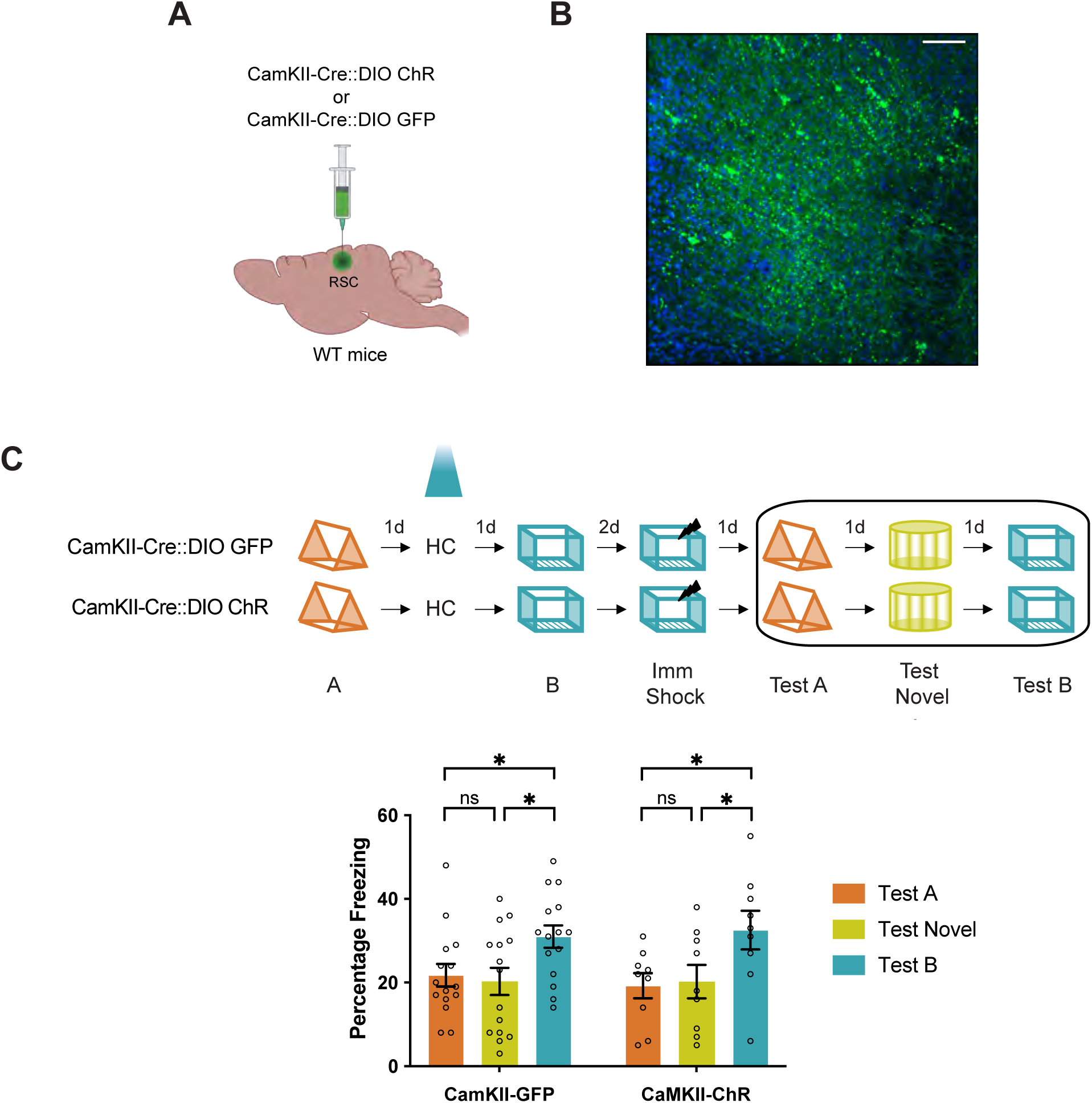
Optogenetic activation of a randomly labeled ensemble does not result in memory linking. (a) Mice received a bilateral injection CamKII-Cre::DIO-ChR or CamKII-Cre::DIO-GFP to label a small subset of RSC ensemble. (b) Representative image of WT mice injected with CamKII-Cre::DIO-ChR-GFP in the RSC. Scale: 20 µm (c) Control (CamKII-Cre::DIO-GFP) as well as experimental (CamKII-Cre::DIO-ChR) mice display low levels of freezing in a novel as well as the previously explored neutral (Context A) context but freeze more in the training context (Context B). (Two-way RM ANOVA, F_time_ (1.9, 42.4) = 9.8, P < 0.0005, Tukey’s multiple comparisons test). Data represent mean ± s.e.m. and each data point, * *p* < 0.05.

**Figure S9.**
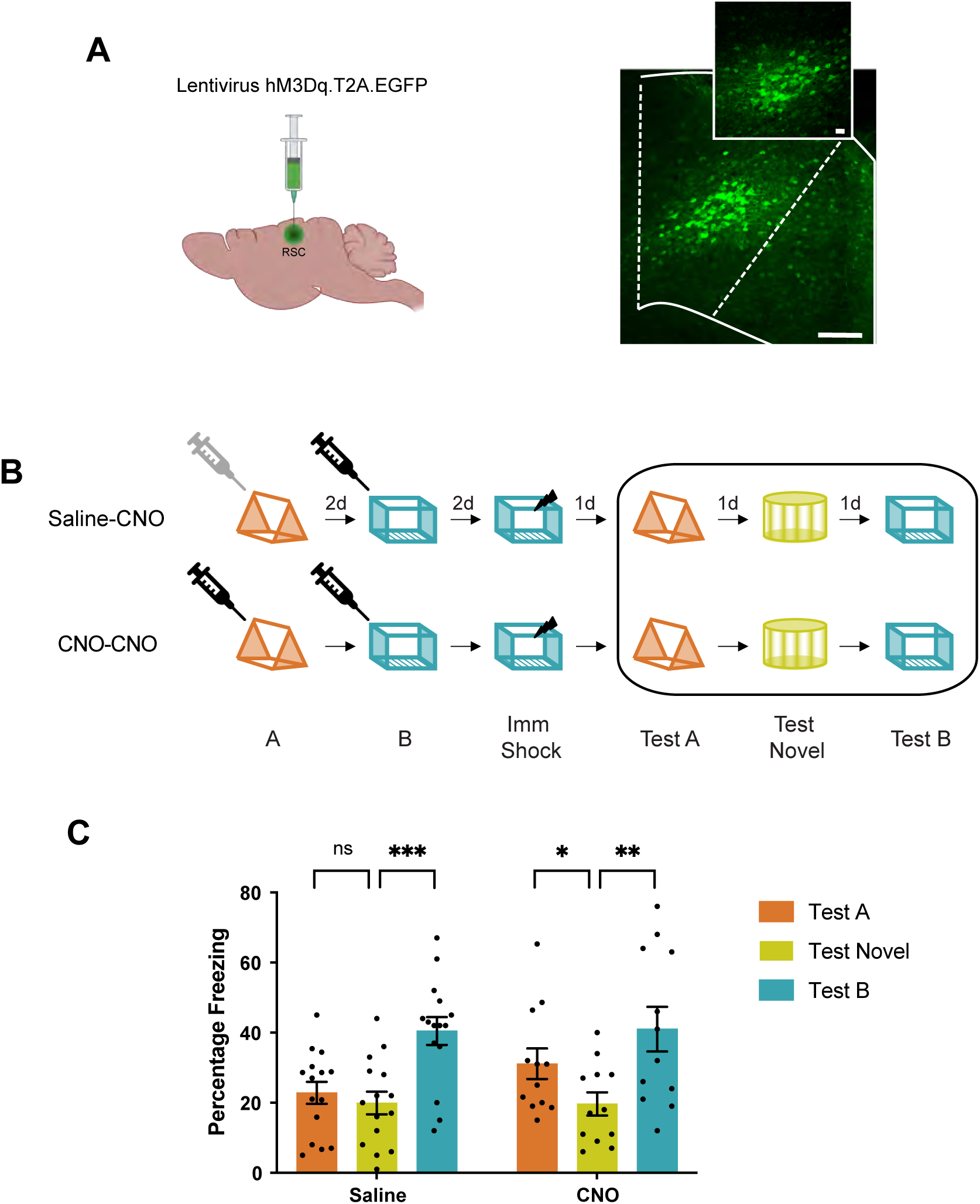
Chemogenetic manipulation of the neuronal ensemble overlap is sufficient to link two distinct contextual memories. (a) All mice received a bilateral injection of lentivirus DREADD hM3Dq-T2A-EGFP which infects a sparse population of RSC neurons. Representative images demonstrating hM3Dq-T2A-EGFP infection of RSC neurons from two mice on the left. Scale: 100 and 20 µm. (b) All mice explored two different contexts 2 days apart and were subsequently shocked in one of these contexts. Neuronal excitability was increased in a small subset of RSC neurons by administering a CNO (0.5mg/kg) injection 45 mins before each context exploration. The control mice only received the CNO injection before the second context exploration. (c) Control mice display low levels of freezing in a novel as well as the previously explored neutral (Context A) context but freeze more in the training context (Context B). In contrast, mice from the experimental group display memory linking: Both the previously explored contexts (Context A and B) elicit high freezing relative to the freezing in a novel context. (Two-way RM ANOVA, F_time_ (1.8, 44.9) = 28.45, P < 0.0001, Dunnett’s multiple comparisons test). The physical contexts presented were counterbalanced to minimize any effect of context similarity. Data represent mean ± s.e.m. and each data point, * *p* < 0.05, ** *p* < 0.01, *** *p* < 0.001.

**Figure S10.**
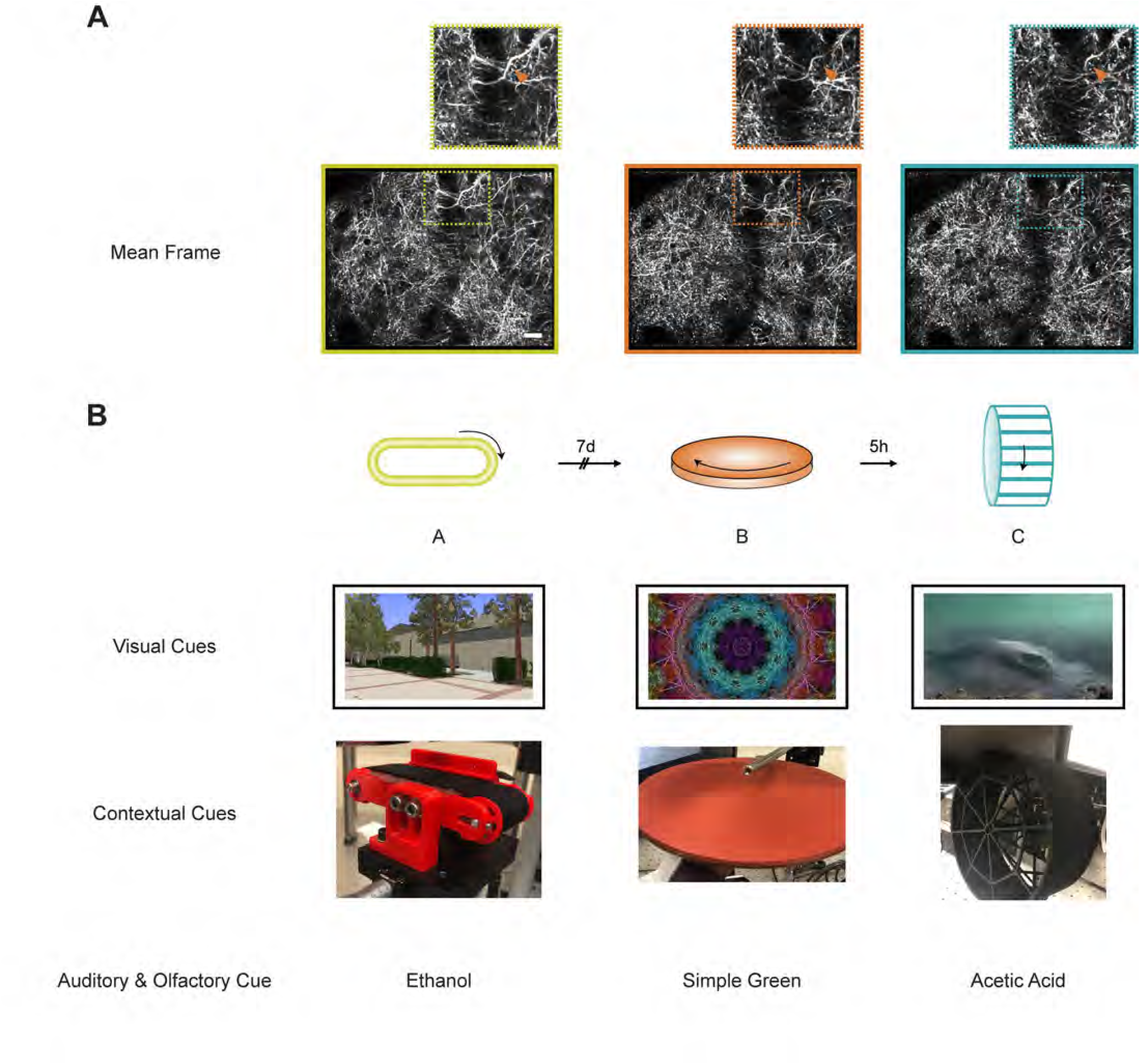
Experimental setup for functional dendritic co-allocation studies: (a) Bottom: Example maximum projection images from three imaging sessions from a mouse. Scale: 20 µm. Top: Inset demonstrates boxed region from each image below magnified to depict the same dendritic segment across sessions. (b) Schematics of 3 distinct contexts (different auditory, visual, and olfactory cues as well as running apparatus) used in the head-fixed experiments. Mice were exposed to 3 distinct contexts 7 days or 5 hours apart in a counterbalanced manner while RSC dendritic transients were imaged.

**Figure S11.**
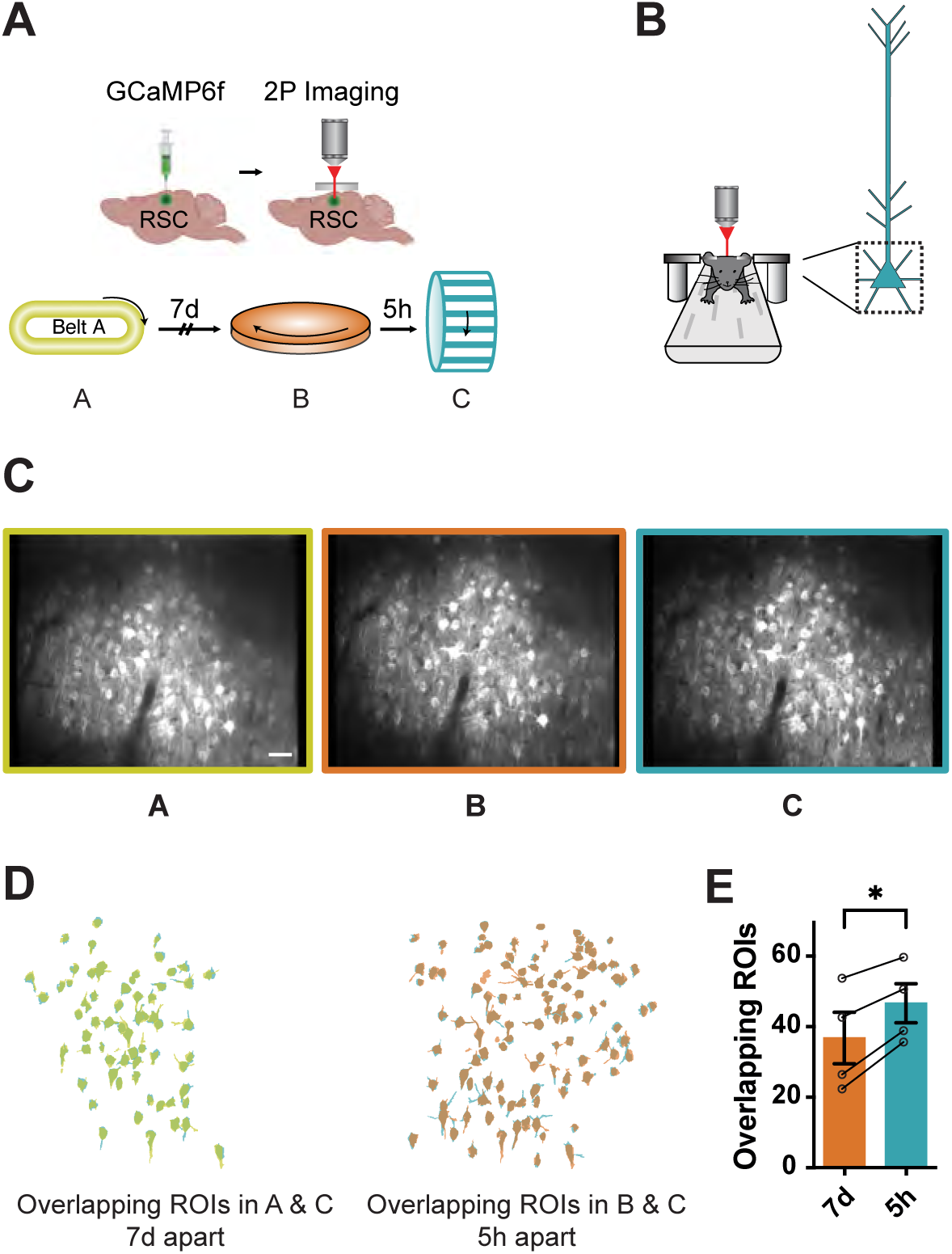
Overlapping neuronal ensemble encodes contexts experienced close in time in head-fixed mice. (a, b) Experimental setup: Head-fixed mice experienced three distinct contexts either 7 days or 5 hours apart while calcium transients from layer V RSC neurons were imaged. (c) Mean frames from three imaging sessions from a mouse. Scale: 40 µm. (d) Overlapping neuronal ROIs reactivated when contexts are separated by 7 days (left) or 5 hours (right) from one mouse. (e) The same neuronal ensemble is more likely to be activated in a head-fixed setting when context exposures are 5 hours (5h) apart vs 7 days (7d) apart. (Paired t-test; t = 5.6; *p* = 0.01; n=4 mice). Data represent mean ± s.e.m. and each data point.

**Figure S12.**
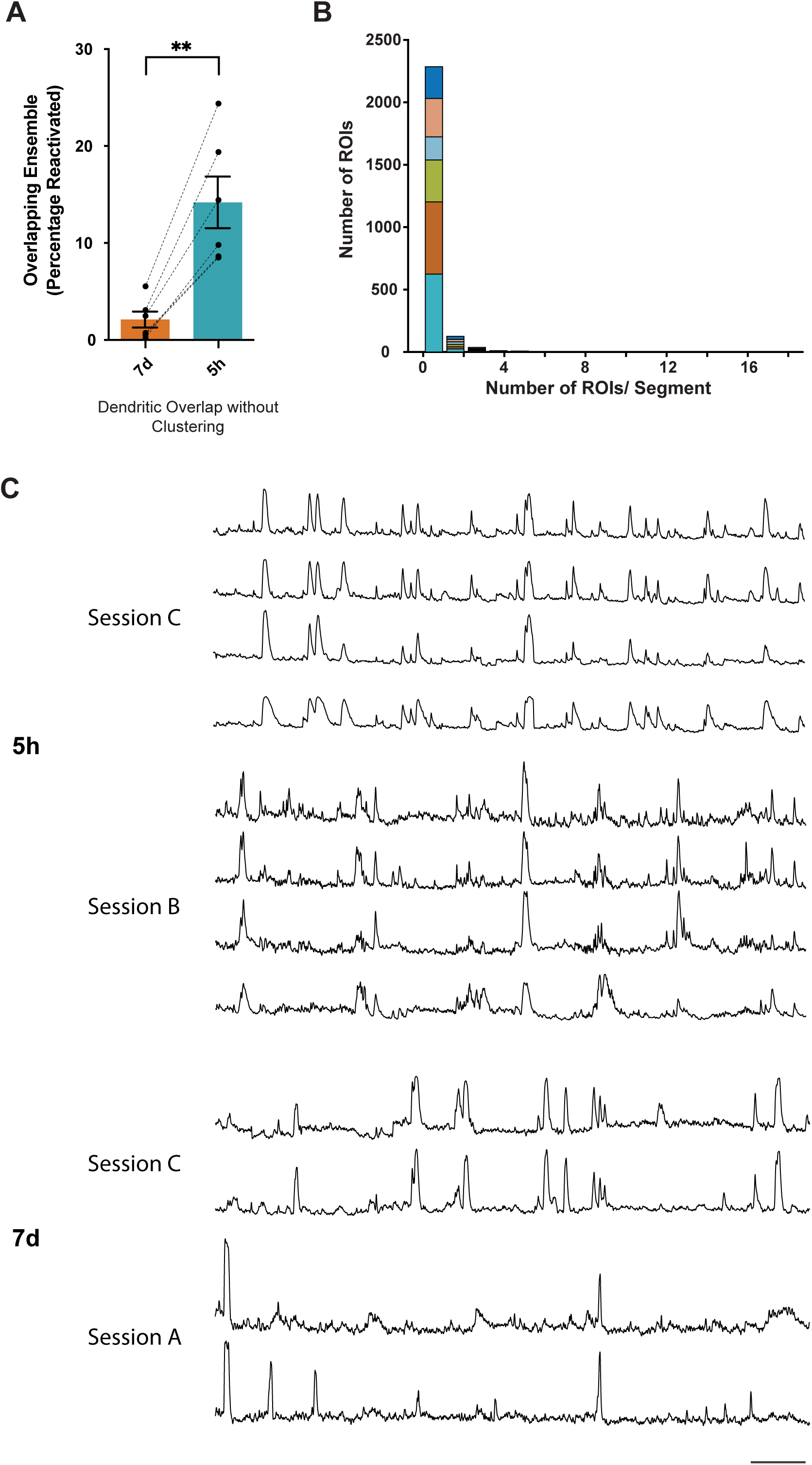
Differences in dendritic overlap are not driven by correlated activity within dendritic branches. (a) Dendritic overlap is higher when context exposures are 5 hours (5h) apart vs 7 days (7d) apart when dendritic ROIs are not clustered together based on correlated activity. (Paired t-test; t = 6.5; *p* < 0.01; n=6 mice). (b) Histogram of the number of ROIs per dendritic segment following clustering (1.15 ± .03 ROIs per cluster). Data from each mouse is depicted in a separate color. (c) For the reactivated dendritic segments (i.e., the clustered ROIs within reactivated segments), ROIs clustered based on their activity in session C (reference session) display high within-cluster correlated activity across sessions. Traces represent z-score of calcium transients. Scale: 30s.

**Figure S13.**
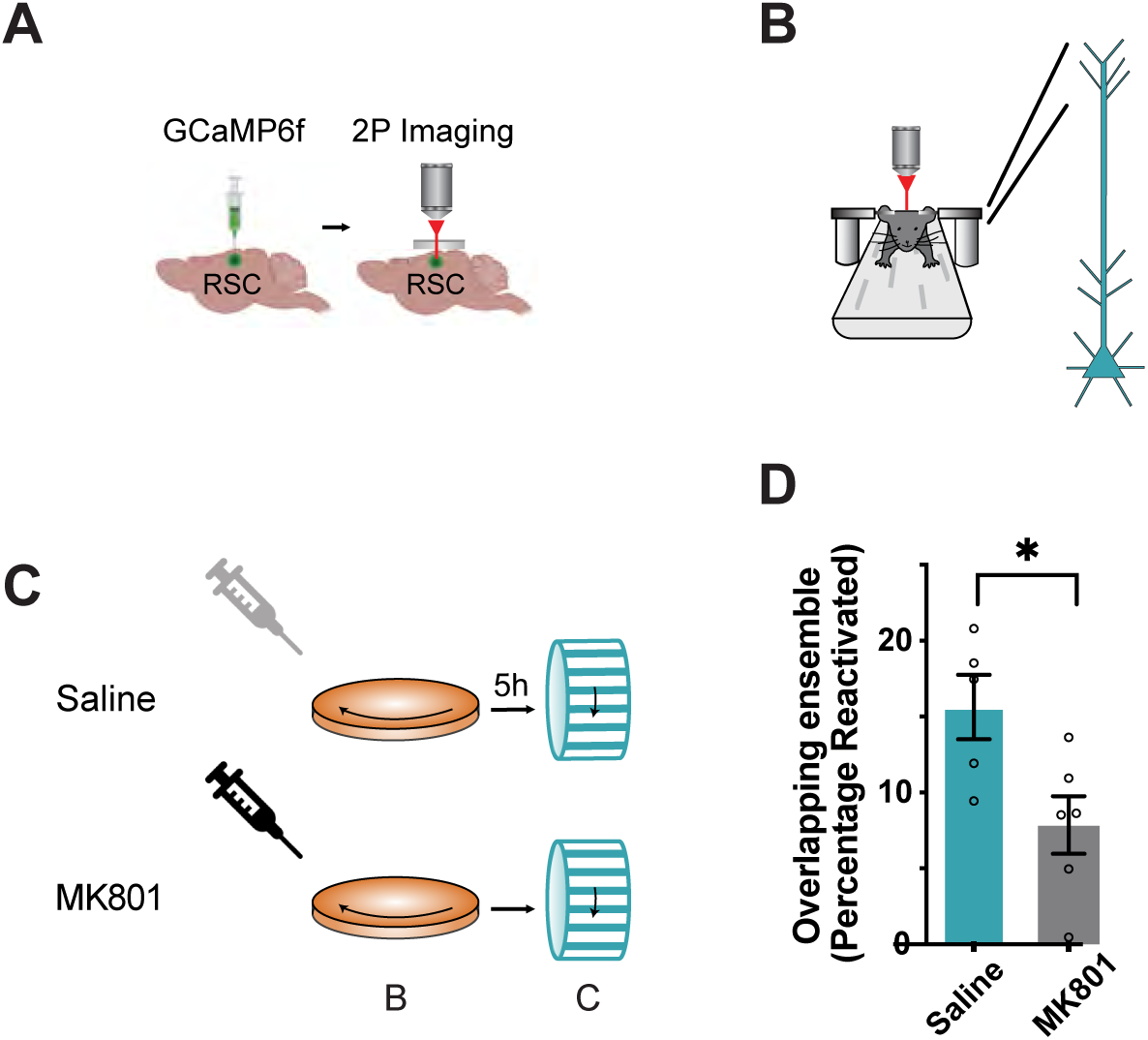
NMDA receptor activation is required for the reactivation of dendritic ensembles. (a-c) Dendritic overlap was measured as described in Figure 3. Mice were administered NMDA receptor antagonist, MK801, 30 minutes prior to the first context exposure. (d) NMDA receptor antagonist, MK801, impairs reactivation of dendritic ensembles following two context exposures 5 hours (5h) apart. (Paired t-test; t = 9.2; *p* < 0.0005; n=6 mice each).

**Figure S14.**
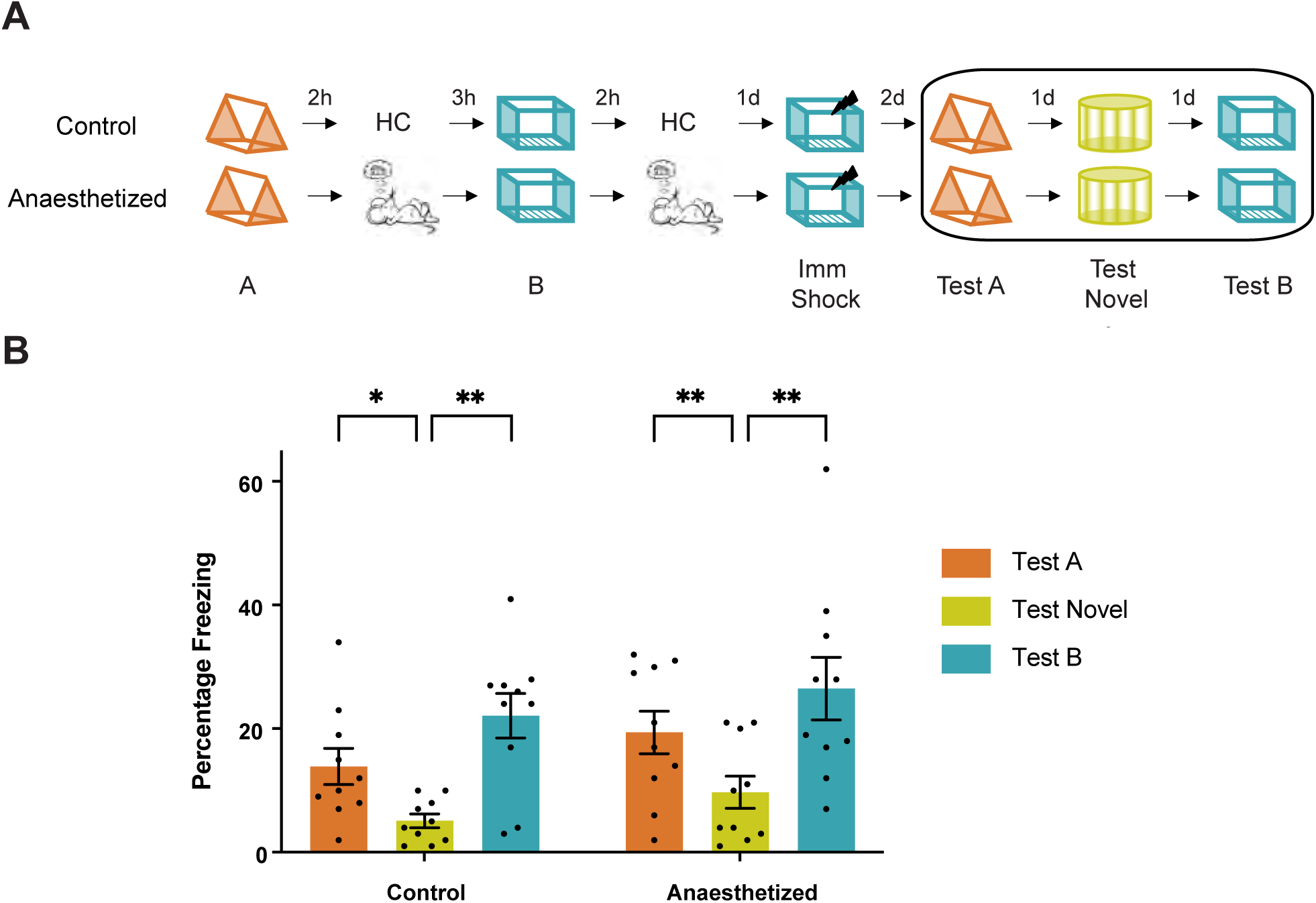
Memories of contexts can still be linked under conditions used during structural imaging. (a) Experimental setup: All mice were handled and habituated in a manner identical to the imaging experiments in Figure 4. Mice experienced two contexts (A and B) 5 hours apart. Two hours following each context exposure, mice were anesthetized for 40 minutes to mimic anesthesia during the imaging sessions to study spine dynamics. (b) The anesthetized mice can link the shock context (context B) to a neutral context (context A) 5 hours apart. For both groups of mice, freezing in the linked (context A), as well as training context (context B), is higher than freezing in a novel context. Therefore, prolonged anesthesia on the day of memory linking does not disrupt memory linking. (Two-way RM ANOVA, F_context_ (1.5, 36) = 27.8, P < 0.0001, Dunnett’s multiple comparisons test). Data represent mean± s.e.m. and each data point, * *p* < 0.05, ** *p* < 0.01.

**Figure S15.**
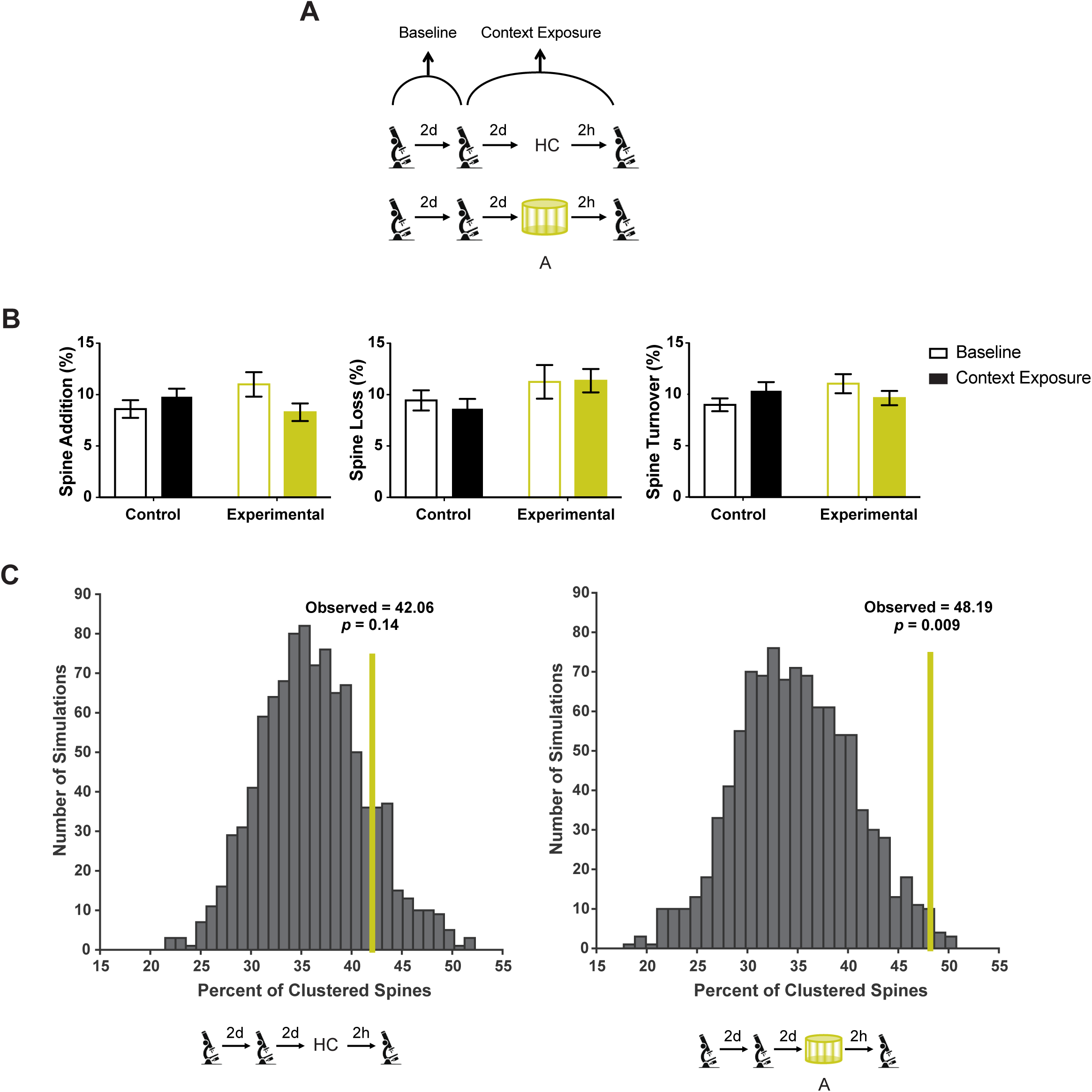
Spine dynamics within the RSC following context exposure. (a) Thy1-YFP mice were imaged every two days (baseline), and the same RSC dendrites were tracked to measure contextual exposure-related spine dynamics. Following two baseline imaging sessions, mice were left in the home cage or exposed to a novel context. (b) Spine addition, spine loss, and spine turnover is not altered within the RSC apical dendrites following context exposure (Two-way RM ANOVA; Sidak’s post hoc tests) (c) Clustered spine addition following context exposure is greater than chance: The histogram shows percent clustering from 1000 simulations of randomized new spine positions, where the percent of new spines within 5 µm of each other was calculated. Yellow line: Percentage spine clustering observed from the data. The percentage of clustered spines is more than that expected by chance for the experimental group (Right, n=6; *p* = 0.009) whereas the percentage of clustered spines is at chance levels for the control group (Left, n=5; p = 0.14). Control: n=44 dendrites (5 mice); Experimental: n=46 dendrites (6 mice). Data represent mean ± s.e.m.

**Figure S16.**
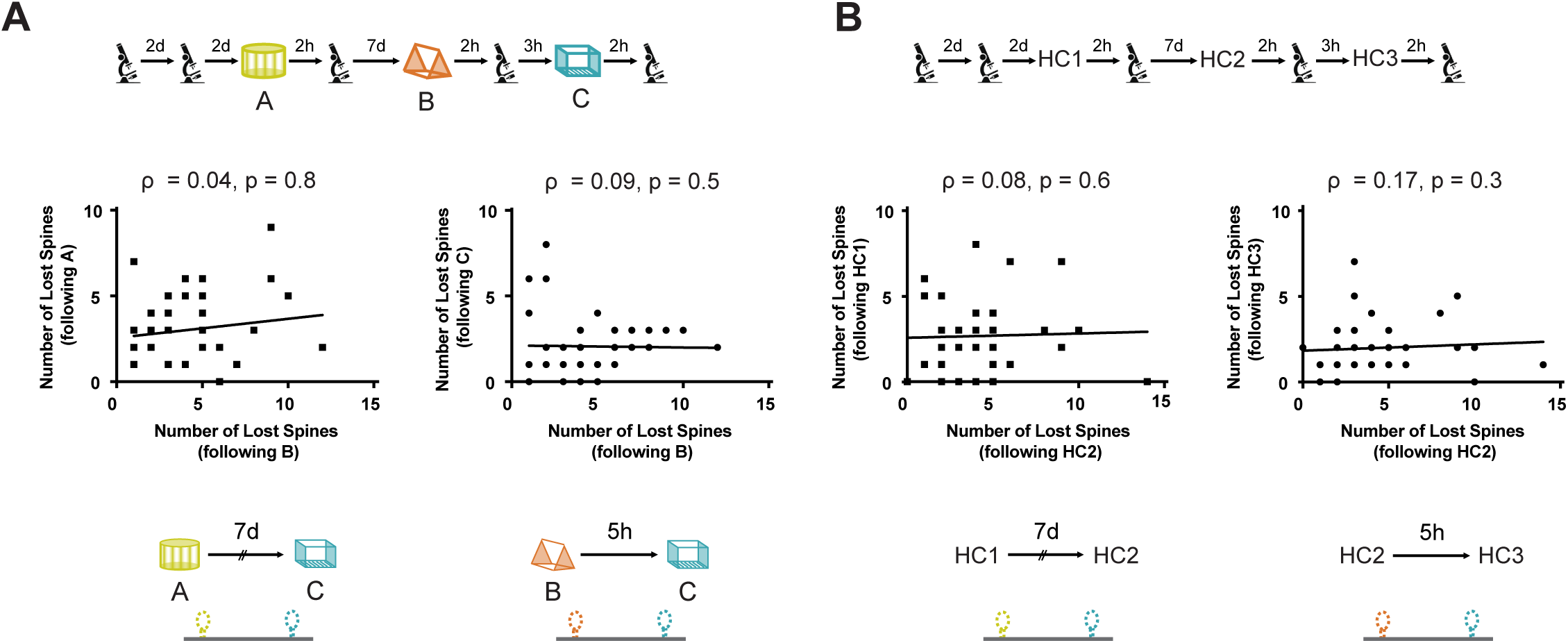
Spine loss during memory linking is not biased to the same dendritic segments. (a) Left: Number of spines lost from a dendritic segment following Context A and B exposure (7 days apart) are not correlated (ρ = 0.04, *p* = 0.8). Right: Number of spines lost from a dendritic segment following Context B and C (5 hours apart) exposure are not correlated (ρ = 0.09, *p* = 0.5). (b) For mice left in their home cages (HC), the number of spines lost from a dendritic segment are not correlated whether imaging sessions are separated by either 7 days (left, ρ = 0.08, *p* = 0.6) or 5 hours (right, ρ = 0.17, *p* = 0.3). Spearman’s correlation was used.

**Figure S17.**
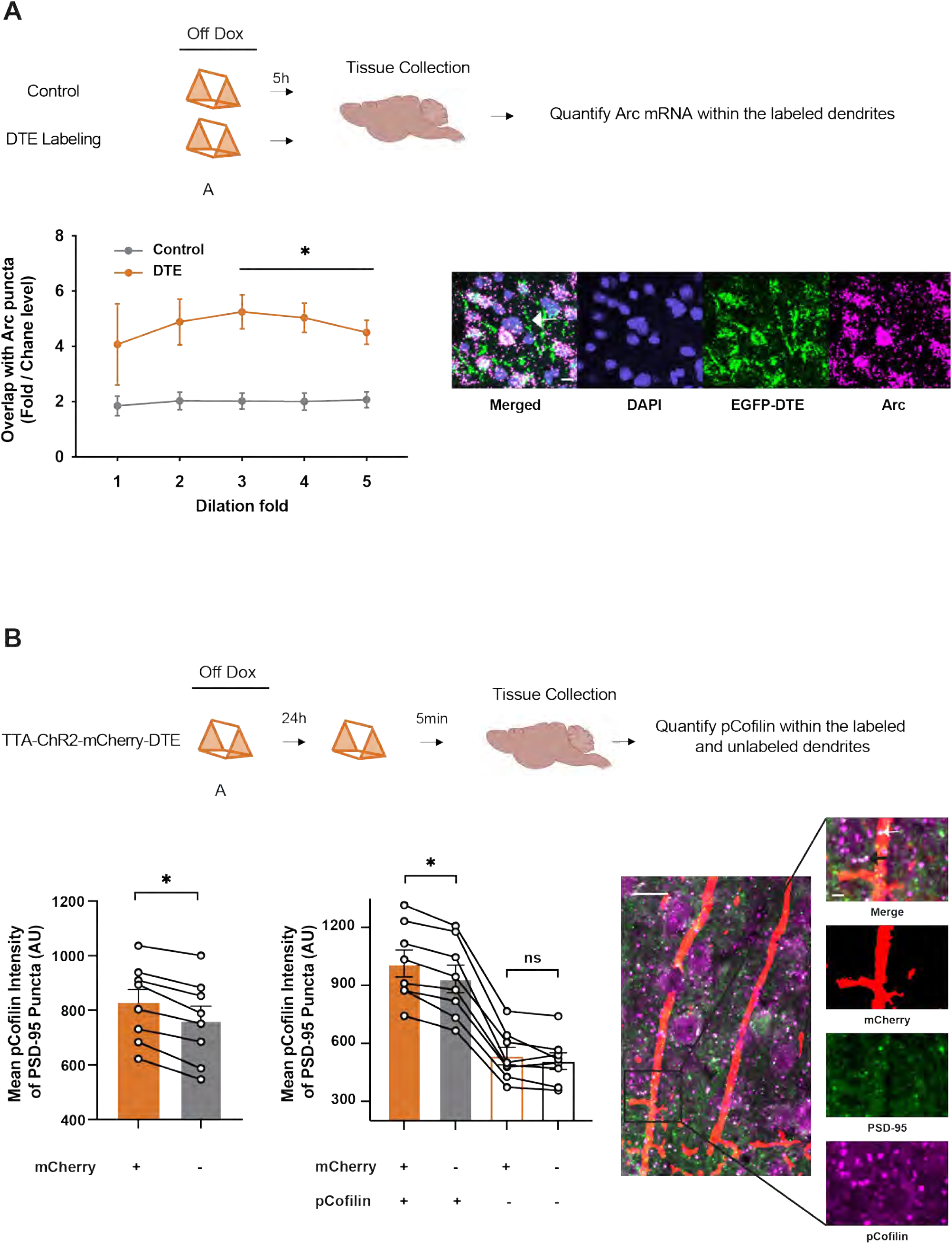
DTE-mediated targeting labels recently activated dendritic segments. (a) Dendritic segments labeled using DTE-mediated strategy are enriched in Arc mRNA. Top: Experimental Design; Mice in Control and DTE group were exposed to a novel context, brains were collected 5 hours later and processed for in situ hybridization to quantify mRNA for Arc and fluorescent reporter protein. Control group was designed to label a small but random subset of dendrites (CamKII-Cre::DIO-GFP in WT mice) and the DTE group used a low titer injection to label activated dendrites sparsely in an activity-dependent manner (cFos-tTa mice with TRE-Opsin-GFP-DTE). Bottom: Regions of interest (ROI) were manually delineated to specifically isolate the fluorescent signal within dendrites (to exclude somatic regions). GFP and Arc signals within these ROIs were automatically segmented. A 1-5 fold dilation of the GFP signal was applied, and the volume of overlap between the dilated GFP signal and the Arc signal was quantified to determine the extent of their colocalization. Arc mRNA was enriched in labeled dendrites in the DTE vs Control group (Control: n = 3, DTE: n = 5; Two-way RM ANOVA, F_Group_ (1, 6) = 10.08, P < 0.05, Sidak’s multiple comparisons test). Scale: 10 µm. (b) Dendritic segments labeled using DTE-mediated strategy are preferentially reactivated upon re-exposure to the original labeling context. Experimental Design (Top): cFos-tTa mice injected with TRE-ChR2-mCherry-DTE virus underwent a novel context exposure and a re-exposure to the same context 24 hours later. Bottom Left: PSD-95 puncta on DTE labeled dendrites displayed more pCofilin labeling (n = 4 mice, 8 slices; Wilcoxon test, p < 0.05). Bottom Middle: Similarly, PSD-95 puncta that were classified as positively labeled for pCofilin (pCofilin+ PSD-95+) displayed higher fluorescence intensity when present on mCherry-labeled dendrites than neighboring pCofilin+ PSD-95+ puncta (Two-way RM ANOVA, F_Group_ (3, 21) = 137.7, P < 0.0001, Sidak’s multiple comparisons test). Bottom right: Representative image depicting pCofilin-positive puncta (magenta) on PSD-95 (green) and mCherry-positive dendrite (red). White and black arrows represent pCofilin-positive, PSD-95-postive puncta on mCherry-positive dendrites and neighboring regions respectively. Scale: 10 µm, Inset Scale: 2 µm. Data represent mean± s.e.m. and each data point, * *p* < 0.05.

**Figure S18.**
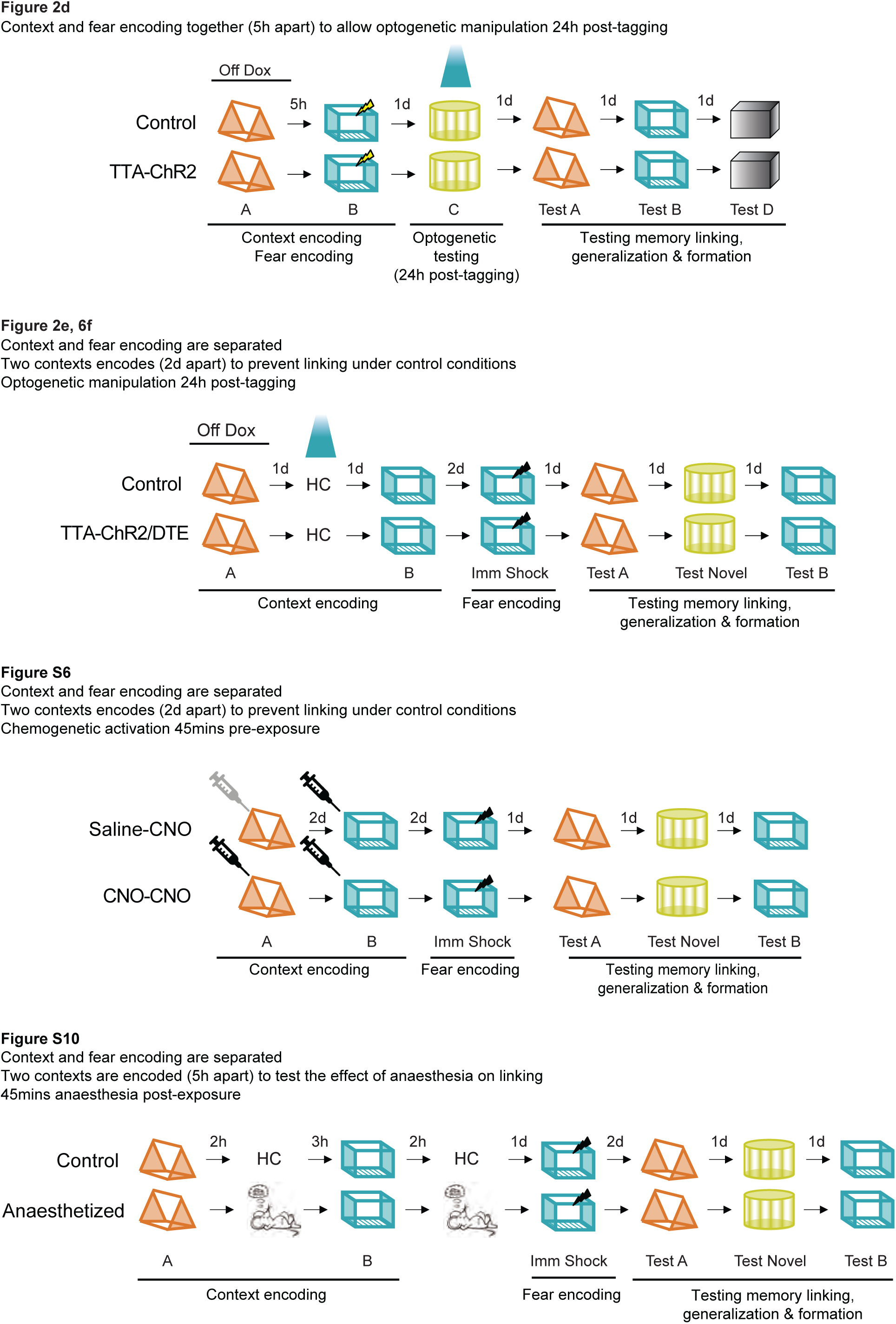
Rationale for behavioral experiments investigating memory linking.

**Supplementary Table 1.**

|  |  | Centroid Distance |  |  |  |  |  |  |
| --- | --- | --- | --- | --- | --- | --- | --- | --- |
|  |  | 3 | 4 | 5 | 6 | 7 | 8 | 9 |
| SFP correlation | 0.6 | 7d: 26.0 ± 4.8<br>5h: 50.6 ± 5.3<br>p = 0.0005 | 7d: 37.0 ± 5.4<br>5h: 59.7 ± 5.3<br>p = 0.002 | 7d: 46.8 ± 5.2<br>5h: 66.1 ± 4.1<br>p = 0.002 | 7d: 51.3 ± 5.0<br>5h: 70.2 ± 3.3<br>p = 0.0007 | 7d: 54.3 ± 4.6<br>5h: 72.6 ± 2.8<br>p = 0.0003 | 7d: 57.5 ± 4.3<br>5h: 74.5 ± 2.6<br>p = 0.0003 | 7d: 59.1 ± 4.2<br>5h: 75.3 ± 2.6<br>p = 0.0005 |
|  | 0.65 | 7d: 25.4 ± 4.6<br>5h: 50.1 ± 5.2<br>p = 0.0005 | 7d: 35.8 ± 5.3<br>5h: 58.8 ± 5.2<br>p = 0.001 | 7d: 44.7 ± 5.2<br>5h: 64.7 ± 4.1<br>p = 0.001 | 7d: 48.8 ± 5.0<br>5h: 68.6 ± 3.2<br>p = 0.0004 | 7d: 51.1 ± 4.7<br>5h: 70.5 ± 2.9<br>p = 0.0003 | 7d: 53.1 ± 4.5<br>5h: 72.0 ± 2.7<br>p = 0.0002 | 7d: 54.3 ± 4.5<br>5h: 72.6 ± 2.7<br>p = 0.0003 |
|  | 0.7 | 7d: 24.5 ± 4.5<br>5h: 49.3 ± 5.3<br>p = 0.0004 | 7d: 34.0 ± 5.1<br>5h: 57.9 ± 5.1<br>p = 0.0007 | 7d: 41.3 ± 5.1<br>5h: 63.3 ± 4.3<br>p = 0.0009 | 7d: 44.7 ± 4.9<br>5h: 66.4 ± 3.8<br>p = 0.0003 | 7d: 46.3 ± 4.7<br>5h: 67.9 ± 3.3<br>p = 0.0002 | 7d: 47.8 ± 4.7<br>5h: 69.1 ± 3.0<br>p = 0.0001 | 7d: 48.8 ± 4.6<br>5h: 69.5 ± 3.0<br>p = 0.0002 |
|  | 0.75 | 7d: 23.5 ± 4.3<br>5h: 47.4 ± 5.2<br>p = 0.0004 | 7d: 31.3 ± 4.9<br>5h: 55.5 ± 5.1<br>p = 0.0005 | 7d: 37.1 ± 5.0<br>5h: 59.8 ± 4.7<br>p = 0.0006 | 7d: 39.6 ± 4.8<br>5h: 62.3 ± 4.5<br>p = 0.0003 | 7d: 41.0 ± 4.7<br>5h: 63.6 ± 4.1<br>p = 0.0002 | 7d: 41.9 ± 4.7<br>5h: 64.6 ± 3.8<br>p = 0.0002 | 7d: 42.3 ± 4.7<br>5h: 64.9 ± 3.9<br>p = 0.0002 |
|  | 0.8 | 7d: 21.7 ± 4.1<br>5h: 45.0 ± 5.2<br>p = 0.0006 | 7d: 27.8 ± 4.7<br>5h: 52.1 ± 5.2<br>p = 0.0007 | 7d: 31.8 ± 4.8<br>5h: 55.8 ± 5.0<br>p = 0.0005 | 7d: 33.4 ± 4.7<br>5h: 57.7 ± 4.8<br>p = 0.0003 | 7d: 34.2 ± 4.7<br>5h: 58.3 ± 4.7<br>p = 0.0003 | 7d: 34.5 ± 4.7<br>5h: 58.8 ± 4.6<br>p = 0.0002 | 7d: 34.8 ± 4.7<br>5h: 58.9 ± 4.6<br>p = 0.0002 |
|  | 0.85 | 7d: 17.8 ± 4.0<br>5h: 39.9 ± 5.0<br>p = 0.0008 | 7d: 22.2 ± 4.6<br>5h: 44.8 ± 5.2<br>p = 0.001 | 7d: 24.2 ± 4.7<br>5h: 47.3 ± 5.0<br>p = 0.0009 | 7d: 24.9 ± 4.6<br>5h: 48.5 ± 4.9<br>p = 0.0006 | 7d: 25.3 ± 4.6<br>5h: 48.8 ± 4.8<br>p = 0.0006 | 7d: 25.5 ± 4.6<br>5h: 49.2 ± 4.8<br>p = 0.0005 | 7d: 25.7 ± 4.6<br>5h: 49.3 ± 4.7<br>p = 0.0005 |
|  | 0.9 | 7d: 11.9 ± 3.1<br>5h: 31.8 ± 4.5<br>p = 0.001 | 7d: 13.9 ± 3.3<br>5h: 34.5 ± 4.6<br>p = 0.0008 | 7d: 14.6 ± 3.4<br>5h: 35.7 ± 4.5<br>p = 0.0007 | 7d: 14.8 ± 3.4<br>5h: 36.2 ± 4.4<br>p = 0.0006 | 7d: 15.0 ± 3.4<br>5h: 36.4 ± 4.4<br>p = 0.0006 | 7d: 15.1 ± 3.4<br>5h: 36.6 ± 4.3<br>p = 0.0005 | 7d: 15.1 ± 3.4<br>5h: 36.7 ± 4.3<br>p = 0.0005 |
|  | 0.95 | 7d: 3.9 ± 1.1<br>5h: 15.3 ± 3.3<br>p = 0.003 | 7d: 4.2 ± 1.1<br>5h: 15.8 ± 3.5<br>p = 0.004 | 7d: 4.2 ± 1.1<br>5h: 16.1 ± 3.4<br>p = 0.003 | 7d: 4.3 ± 1.1<br>5h: 16.2 ± 3.4<br>p = 0.003 | 7d: 4.3 ± 1.1<br>5h: 16.2 ± 3.4<br>p = 0.003 | 7d: 4.3 ± 1.1<br>5h: 16.2 ± 3.4<br>p = 0.003 | 7d: 4.3 ± 1.1<br>5h: 16.4 ± 3.4<br>p = 0.003 |

**Supplementary Table 2.**

| Parameters of the computational model |  |  |
| --- | --- | --- |
| $N_{\text{pyr}}$ | Number of excitatory neurons | 400 |
| $N_{\text{inh}}$ | Number of inhibitory neurons | 50 dendrite-targeting (DT)<br>50 soma-targeting (ST) |
| $N_{\text{dend}}$ | Number of dendritic subunits per neuron | 10 for excitatory<br>1 for interneurons |
| $N_{\text{pyr} \rightarrow \text{ST}}$ | Synapses from excitatory neurons to soma-targeting(ST) interneurons | Count: 1000<br>Weight: 0.6 |
| $N_{\text{pyr} \rightarrow \text{DT}}$ | Synapses from excitatory neurons to dendrite-targeting (DT) | Count: 1000<br>Weight: 0.3 |
| $N_{\text{ST} \rightarrow \text{pyr}}$ | Synapses from ST interneurons to excitatory neurons | Count: 10000<br>Weight: 0.5 |
| $N_{\text{DT} \rightarrow \text{pyr}}$ | Synapses from DT interneurons to excitatory neurons | Count: 2000<br>Weight: 0.3 |
| $N_{\text{input} \rightarrow \text{pyr}}$ | Synapses from input afferents to pyramidal dendrites per encoded memory | Count: 23000<br>Initial Weight: 0.16 – 0.36 |
| $E_L$ | Leakage reversal potential | 0 mV |
| $g_E / g_I$ | Dendritic excitatory / inhibitory synaptic conductance | 22nS / 20nS |
| $g_{Ld} / g_L$ | Dendritic/somatic leak conductance | 10nS / 8nS |
| $g_{\text{Inh}}$ | Somatic inhibitory current scaling constant | 600nS |
| $\tau_{\text{Inh}}$ | Somatic inhibitory current time constant | 30msec |
| $E_E / E_I$ | Excitatory /inhibitory synapse reversal potential | +70mV / -10mV |
| $C$ | Membrane capacitance | 200pF |
| $\tau_{\text{dend}}$ | Dendritic membrane time constant | Inhibitory: 20msec<br>Excitatory: 25msec |
| $V_d$ | Dendritic Depolarization | -10mV < $V_d$ < 70mV |
| $g_{\text{ax}}$ | Axial conductance | 36nS |
| $\theta_{\text{soma}}$ | Voltage threshold for somatic spikes | 18mV |
| $\tau_{\text{adapt}}$ | Adaptation time constant of excitatory neurons | 200msec |
| $\beta_{\text{adapt}}$ | Adaptation reset constant | Baseline excitability: 9<br>High excitability: 6.5 |
| $a_{\text{adapt}}$ | Adaptation coupling parameter | 0.02 |
| $\Theta_{\text{PRP}}$ | Calcium threshold for somatic Plasticity-Related Protein (PRP) synthesis | 40.0 |
| $\Theta_{\text{dend}}$ | Calcium threshold for dendritic excitability | 2.0 |
| $\Theta_{\text{soma}}$ | Calcium threshold for somatic excitability | 40.0 |
| $\tau_H$ | Time constant of homeostatic synaptic scaling | 1440 hours |

## METHODS

### Animals

All experimental protocols were approved by the Chancellor’s Animal Research Committee of the University of California, Los Angeles, in accordance with NIH guidelines. cFos-tTa mice that express tetracycline transactivator (tTA) protein under the control of the c-fos (also known as Fos) promoter were maintained in a C57BL/6N background. Adult (3–8 months old) male and female Thy1-YFP-H mice (Jackson Laboratories, Stock No: 003782) were used for structural imaging experiments. C57BL/6N Tac mice were purchased from Taconic Farms (Germantown, NY) for all other experiments.

### Viral construct

pAAV-Syn-GCaMP6f-WPRE-SV40 was a gift from Douglas Kim & GENIE Project (Addgene viral prep # 100837-AAV1; http://n2t.net/addgene:100837; RRID: Addgene 100837). The lentivirus hM3Dq-T2A-EGFP vector was derived as previously described in Cai et al., 2016 ^12^. Finally, AAV1-TRE-hChR2-mCherry, AAV1-TRE-hChR2-mCherry-DTE and AAV1-TRE-mCherry-DTE were derived in our laboratory. Briefly, to construct a vector for TRE-driven hChR2 expression, a CamKIIa promoter from pAAV-CamKIIa-hChR2(H143R)-mCherry (Addgene #26975) was replaced with TRE promoter from pAAV-RAM-d2tTA∷TRE-NLS-mKate2-WPRE (Addgene #84474) using MluI/AgeI digestion. The DTE sequence of Arc mRNA was PCR-amplified from cDNA of 14 weeks Spraque-Dawley rat using primers as previously described ^18^ and inserted into the pAAV-TRE-hChR2-mCherry vector using EcoRI/HindIII. The pAAV-TRE-hChR2-mCherry and pAAV-TRE-hChR2-mCherry-DTE were subjected to AgeI/BsrGI digestion for construction of mCherry vectors, respectively. The mCherry was digested by AgeI/BsrGI from pmCherry-N1. AAV production was conducted as previously described in detail ^100^ with modifications.

### Surgery

Mice were anesthetized with 1.5 to 2.0% isoflurane for surgical procedures and placed into a stereotactic frame (David Kopf Instruments, Tujunga, CA) on a heating pad. Artificial tears were applied to the eyes to prevent drying. Subcutaneous saline injections were administered throughout each surgical procedure to prevent dehydration. In addition, carprofen (5 mg kg−1) and dexamethasone (0.2 mg kg−1) were administered both during surgery and for 2-7 days post-surgery. A midline incision was made down the scalp, and a craniotomy was performed with a dental drill. Water with amoxicillin was administered for two weeks.

### Miniscope experiments

#### Surgeries

Mice were unilaterally microinjected with 500 nl of AAV1-Syn-GCaMP6f-WPRE-SV40 virus at 20-120 nl/min into the RSC using the stereotactic coordinates: −2.3 mm posterior to bregma, 0.5 mm lateral to the midline, and −0.8 mm ventral to the skull surface. Immediately afterward, the microendoscope (a gradient refractive index or GRIN lens) was implanted above the injection site. For this procedure, a 2.0 mm diameter circular craniotomy was centered above the virus injection site. The microendoscope (0.25 pitch, 0.50 NA, 2.0 mm in diameter, and 4.79 mm in length, Grintech Gmbh) was slowly lowered with a stereotaxic arm above the craniotomy 450 μm ventral to the surface of the skull. Next, a skull screw was used to anchor the microendoscope to the skull. Both the microendoscope and skull screw were fixed with cyanoacrylate and dental cement. Kwik-Sil (World Precision Instruments) was used to cover the microendoscope. Three weeks later, a small aluminum baseplate was cemented onto the animal’s head atop the previously placed dental cement. The microscope was placed on top of the baseplate and locked in a position such that the field of view contained cells and visible landmarks, such as blood vessels, and these appeared sharp and in focus. Finally, a plastic headcap was fit into the baseplate and secured with magnets. Independent experiments confirmed that GCaMP6f expression was limited to RSC neurons (Figure 1b).

#### Miniscope Behavior

Customized UCLA Miniscopes V3 with a 20mm achromatic doublet lens and modified housing were used to allow imaging 300 μm below the surface of the GRIN lens allowing imaging of RSC neurons. Using head-mounted miniature microscopes (UCLA Miniscopes V3)(*4*, *38*), we imaged GCaMP6f-mediated calcium dynamics in RSC neurons of GRIN lens implanted mice that explored distinct contexts. Before imaging sessions, mice were handled and habituated to the experimental conditions, including carrying the Miniscope while it was tethered to the implanted GRIN lens. Mice were exposed to each context (with distinct visual, auditory, and olfactory cues) for 10 mins during which calcium transients were recorded (Figure 1c for representative calcium transients). Context A was separated from Context B by 7 days, and Context B and Context C were separated by 5 hours (Figure 1e). The actual contexts used were counterbalanced and comprised of rectangular plastic containers (15 ± 1 by 11 ± 1 inches) that were covered with various visual cues.

#### Miniscope Analysis

Calcium imaging data were registered to remove small movement artifacts using NormCorre ^101^. This was followed by automated segmentation, demixing, and denoising of calcium signals using constrained non-negative matrix factorization for endoscopic data (CNMFe) ^102^. We used a modified version of the Miniscope analysis package developed by Guillaume Etter (Sylvain Williams Lab, McGill University) for data analysis ^103^. Recordings from multiple sessions of the same mouse were aligned using an amplitude-based registration algorithm used for within-session registration, except the algorithm was only applied to the mean frame from each session. Once regions of interest (ROIs-putative neurons) from two sessions were registered, ROIs across two sessions were matched to each other using a distance (between ROI centroids) and correlation (between ROIs spatial footprints - SFPs) measure. The neuronal ensemble overlap was calculated as the percentage of ROIs activated in both contexts divided by the average number of ROIs identified in each imaging session. Neurons were matched across days based on distance (< 4 pixels) and correlation (> 0.9) thresholds. These results (Figure 1e) were consistent and robust for a range of distance (3-9 pixels) and correlation (0.6-0.95) thresholds used to match segments across days (data not shown).

In a parallel approach, we aligned and concatenated the imaging data from the three context exposures into a single video file (followed by motion correction and segmentation as described above) and analyzed the data such that we were able to detect and track the activity of ROIs across all different sessions as well as investigate their modulation during context exploration. The raw data from CNMFe extracted putative neurons was deconvolved into spike probabilities using the foopsi thresholded method (OASIS toolbox). Finally, the spike probabilities from single frames were binarized between 1 (active) and 0 (inactive). For each neuron, the firing rate (number of active frames per second) for each session was estimated. Population Vector Correlations (PVCs) were calculated as the Pearson correlation between the average firing rate (per session) of each neuron across two imaging sessions (Figure S2).

Naïve Bayes (NB) Binary Classifier. The activity of each neuron during each 10 min session was resampled into various time bin sizes (0.5-60 second bins, step size 0.5s; Figure S2). Each resampled data with a specific bin size was used as trials from each session. The classifier was trained on 90% of the data and we used the information contained in the probability of activity from each neuron to test the remaining 10% data (10-fold cross-validation strategy) as belonging to the two given sessions. The area under the receiver operating characteristic curve (AUC) was calculated for the first context (A for 7d or B for 5h; Figure 1g) using the Wilcoxon-Mann-Whitney statistic. The quality of the classification is defined by AUC, which ranges from 0 to 1. AUC = ~0.5 means sorting at chance levels by the classifier.

Pairwise correlations (PWC) maps for each session were calculated by binning neuronal activity into 100ms bins to compute the Pearson correlation for each pair of neurons (Figure S5a,b). PWC stability was calculated as the Pearson correlation between PWC maps from different sessions, excluding the main diagonal (correlation between each neuron with itself) and cell pairs below the main diagonal (such that each cell pair was represented only once). Since artificially high correlations can arise due to sub-optimal demixing of calcium signals from nearby ROIs, we computed the PWC analysis while ignoring the PWCs from nearby cell pairs. We defined nearby cells as cell pairs where spatial footprints (SFP) had any overlap or where the centroid to centroid distance was shorter or equal to 20 pixels (~40 µm). To control for the different number of neurons detected for different mice, we calculated PWC stability between 2 sessions by randomly subsampling a group of 10 cells, computing the PWC map for each of the sessions using these cells, and computing the Pearson correlation between the two PWC maps. This process was repeated 1000 times and the final PWC stability was defined as the average of these 1000 values. The absolute PWC per imaging session and PWC stability across sessions follows the same trend whether the analyses were done with or without nearby cells or with subsampling of 50 cells instead of 10 cells (t=3.61, *p* = 0.006). For brevity, we only present analyses that excluded the nearby neurons.

### Optogenetic experiments

Adult male and female (3-8 months) cFos-tTa transgenic and their wild-type littermates maintained on doxycycline chow (for 1 month or more) were bilaterally microinjected with 500 nl of AAV1-TRE-hChR2-mCherry, AAV1-TRE-hChR2-mCherry-DTE, AAV1-TRE-mCherry-DTE virus at 20-50 nl min^−1^ into the RSC using the stereotactic coordinates: −2.3 mm posterior to bregma, 0.5 mm lateral to the midline, and −0.8 mm ventral to the skull surface. For Figure S8, wild-type mice were injected with a cocktail of CamKII-Cre (Addgene 105558-AAV, diluted 1:10^3^) with DIO-hChR2 (Addgene 35509-AAV9 – Experimental) or DIO-GFP (Control). Following viral injections, bilateral optogenetic cannulae (Doric Lenses Inc.; DFC_200/240-0.22_0.5mm_GS1.0_FLT) were implanted over the injection site at −0.45 mm ventral to the skull surface.

### Chemogenetic experiments

Adult (3-5 months old) C57Bl/6NTac male mice were bilaterally microinjected with 1000 nl of lentivirus hM3Dq.T2A.EGFP at 20-100 nl/min into the RSC using the stereotactic coordinates: −1.95 and −2.65 mm posterior to bregma, 0.5 mm lateral to the midline, and −0.8 mm ventral to the skull surface. Following viral injections, mice were allowed to recover from surgeries for 3 weeks before being handled (3 days) and habituated (3 days) for a modified two-day memory linking experiment. To ensure that the same RSC neurons are recruited for encoding these different contexts, we transiently increased the intrinsic excitability of a small subset of RSC neurons by administering a clozapine N-oxide (CNO, 0.5mg/kg) injection 45 mins before each context exploration ^12^. The control mice only received the CNO injection before the second context exploration. Following this, the mice underwent the memory linking paradigm described below.

### Memory linking studies

Linking of context memories was carried out as previously described ^12^. Briefly, mice were handled for 3 days (2–5 min/day) and then habituated to transportation and experimental room/s for 3-5 days (2-5 mins/ day). In the memory linking task, mice explored 2 distinct contexts (A and then B, for 10 mins each) separated by 5h (Figure 2d and S14) for linking under control conditions or 2 days (Figures 2e, 6f, S8 and S9) to ensure a robust lack of linking under control conditions^48^. The actual contexts presented were counterbalanced to minimize any effect of context similarity. <u>For</u> Figure 2d: The context exposure in chamber B also included a 2s, 0.75mA footshock that was delivered 58 seconds before the end of context exposure. This was done to shorten the window of time between the encoding of the first contextual memory (for activity-dependent tagging), subsequent linking, and optogenetic manipulation to 24 hours post-tagging. All optogenetic manipulations were performed 24 hours post-tagging to ensure sufficient expression of the tagged opsin (Figure S18).

#### Manipulation to link normally independent memories

For experiments where we extended the time window for memory linking by manipulating neuronal (Figure 2e) or dendritic overlap (Figure 6f), the two context exposures were separated by 2 days. We did this for two reasons. First, two contexts explored 5 hours apart are linked but when contexts are explored 2 days apart, they are not linked^12,48^. Second, Channelrhodopsin expression under the TetTag system peaks at 24 hours ^43^. This allowed us to extend the window of memory linking (to 2 days) by using the transient expression of Channelrhodopsin on a day after the first context exposure.

Immediate shock: <u>For</u> Figures 2e and 6f: Two days following the last context exposure (in B), mice were placed in context B again for an immediate foot shock (10 second baseline, 2 second shock, 0.7-0.75mA, 28-58 second post-shock period). <u>For</u> Figure S14: To compensate for the lower freezing seen in C57Bl/6N Jackson mice (the genetic background of the Thy1-YFP mice), the immediate shock protocol was modified to a 10 second baseline, two shocks for 2 seconds each, 0.75mA, 15 seconds apart.

Testing: During the testing phase, mice were tested in the designated contexts (5 minutes each) on three separate days to minimize any effects of testing animals in one context on subsequent tests in another context. The order of testing was also chosen to control for any gradual increase or decrease in freezing. The actual contexts were counterbalanced. Freezing was assessed via an automated scoring system (Med Associates) with 30 frames per second sampling rate; the mice needed to freeze continuously for at least one second before freezing could be counted.

### Tagging of RSC ensemble

Mice were allowed to recover from surgeries for 3-5 weeks before being handled (3 days) and habituated (3-5 days) for behavioral exposure as well as optogenetic manipulation. The day after the last day of habituation, mice were taken off doxycycline chow (40 mg kg^−1)^ and placed on regular chow and tTA expression was allowed for 3 days before behavioral tagging for the memory linking experiments. The activity-dependent tag was shut off by administration of high dox chow (200 mg kg^−1^) 90 minutes after behavioral tagging. For experiment in Figures 2d and S7 a subset of animals was also administered doxycycline intraperitoneally (i.p, 50 ug/ gram of body weight; 2 hour post tagging episode) to ensure that the tagging window is closed even in the absence of immediate feeding. The dose and timing were chosen because of its effectiveness in initiating tagging with the Tet-On system ^104^. Doxycycline is also detectable at near peak levels 2 hours post injection in the brain tissue ^105^. Our behavioral results remain the same with and without doxycycline administration i.p. and hence we have combined data from these two sets of experiments in Figure 2d. Set 1 (without i.p. doxycycline): Animals were placed back on high dox chow 90 mins post context A exposure. Control, n = 4; TTA-ChR2, n = 6; Two-way RM ANOVA, F_Interaction_ (1, 8) = 5.4, p < 0.05; Sidak’s multiple comparisons test, p < 0.005). Set 2 (with i.p. doxycycline): Animals were placed back on high dox chow and injected with doxycycline i.p. 2 hours post context A exposure. Control, n = 12; TTA-ChR2, n = 8; Two-way RM ANOVA, F_Interaction_ (1, 18) = 5.4, *p* < 0.05; Sidak’s multiple comparisons test, p < 0.005). Combined: Control, n = 16; TTA-ChR2, n = 14; Two-way RM ANOVA, F_Interaction_ (1, 28) = 12.8, *p* < 0.005; Sidak’s multiple comparisons test, p < 0.0001).

### Optogenetic Manipulations

All optogenetic manipulations were performed 24 hours following the tagging event to ensure sufficient expression of the opsins.

#### Reactivation of tagged ensembles in home cage

For Figures 2e and 6f, ensembles tagged during the first context exposure were reactivated in the home cage using a 473nm laser (5 ms pulses, 5 Hz) for 10 minutes. *Testing:* For Figure 2c-d, mice were placed in an open field and freezing behavior was recorded using a digital camera. Following a 3-minute baseline period, the tagged RSC ensemble was reactivated using a 473nm laser (5 ms pulses, 5 Hz) for one minute followed by a one-minute interval with no stimulation. This pattern of stimulation was repeated three times, and the time spent freezing during the three epochs was averaged.

### Immunostaining

Mice were transcardially perfused with 0.1 M phosphate buffer followed by 4% PFA (4% paraformaldehyde in 0.1 M phosphate buffer) and after perfusion, brains were kept in the fixation solution overnight at 4 °C, then transferred to 30% sucrose solution for 48 h, sectioned (40 µm thickness) on a cryostat and stained while free-floating. For staining synaptic proteins, tissue was sectioned at 15 µm thickness.

The sections were blocked for 1 h at room temperature in 0.3% Triton-X in PBS (PBST) and 10% normal goat serum (Vector Laboratories, S-1000) solution. Primary and secondary antibodies were diluted in the same blocking solution. The primary antibody (guinea pig anti-RFP: SySy 390004, chicken anti-RFP: SySy 409006, anti-PSD95: SySy 124308, anti-phospho-Cofilin; Millipore C8992) incubation was overnight (~18 h) at 4 °C, and the secondary antibody (Alexa Fluor 488, 568, 647: Invitrogen) incubation was 2 h at room temperature, both with constant shaking. Brain slices were incubated with 4’,6-diaminodino-2-phenylindole (DAPI, Invitrogen, 1:1000) for 10 min, washed with PBST two times and PBS once before mounting onto slides. Immunostaining images were acquired with a Nikon A1 Laser Scanning Confocal Microscope (LSCM) and analyzed with automatic spot-detection algorithm (Imaris 9.2, Bitplane AG) and manually verified.

### In situ hybridization

Control mice were WT C57BL/6N mice that were injected with a cocktail of CamKII-Cre (Addgene 105558-AAV1, diluted 1:10^3^) and DIO-GFP viruses to label a sparse and random subset of RSC neurons. DTE group comprised of cFos-tTa mice injected with TRE-Opsin-GFP-DTE (10^11^ gc/ml) to sparsely label dendrites in an activity-dependent manner. Mice brains were dissected and fast-frozen in optimal cutting temperature compound (OCT) using dry ice without paraformaldehyde (PFA) fixation. Frozen sections were sliced (15µm). In situ hybridization was performed using the RNAscope™ Multiplex Fluorescent Reagent Kit v2 (ACD, 323100) according to the manufacturer’s instructions. Probe-GFP (ACD, 409011) and Probe-Mm-Arc (316921) were used for mRNA labeling.

The images were acquired using NIS-Elements AR (Nikon, v.4.40.00) with a Nikon A1 Laser Scanning Confocal Microscope (LSCM). Analysis of the confocal images was conducted using NIS-Elements AR Analysis software (Nikon, v.4.40.00). To perform the analysis, regions of interest (ROI) were manually delineated to specifically isolate the GFP signal within dendrites (excluding the soma). The GFP and Arc signals within these ROIs were automatically segmented using thresholding techniques. A 1-5 fold dilation of the GFP signal was applied, and the volume of overlap between the dilated GFP signal and the Arc signal was quantified to determine the extent of their colocalization as follows:

**Chance level = (GFP volume / ROI total volume) * (Arc Volume / ROI total volume)**

**Arc overlap possibility = (GFP and Arc overlap volume / ROI total volume) / Chance level**

### Structural 2p imaging

#### Methods

Adult (3-8 months old) male and female Thy1-YFP-H mice were used for structural imaging experiments. Mice underwent window implantation surgeries as previously described ^19^. Briefly, a square region of the skull 2-3 mm in width was marked using stereotactic coordinates (RSC: center at bregma −2.3 mm AP). The skull was thinned using a dental drill and removed. After cleaning the surgical site with saline, a custom cut sterilized coverslip (square, 2×2mm unilateral or 3×3mm bilateral) was placed on the dural surface and fastened with adhesive and dental acrylics to expose a square window of approximately 2 mm. Next, an aluminum bar with a threaded hole was attached to stabilize the mice during imaging sessions. Mice were allowed to recover for two or more weeks before the first imaging session. Following recovery from surgery, mice were handled and habituated as per the memory linking paradigm. In addition, mice were also habituated for transportation to the imaging room as well as anesthesia. After handling/habituation (1-2 days later), mice underwent the first home cage baseline imaging session. Two days later mice underwent the second baseline imaging session. Two days following the last baseline imaging session, a subset of mice was exposed to a novel context ‘A’ for 10 minutes while another subgroup remained in the home cage (control group). After 7 days, mice were exposed to a novel context (B, 10 minutes) which was followed by a third novel context exposure (C, 10 minutes) five hours later. Mice were imaged 2 hours after each context exposure.

Two-Photon imaging measuring spine dynamics. A custom-built two-photon laser scanning microscope was paired with a Spectra-Physics two-photon laser tuned to 920nm. A 40x 1.0 NA water immersion objective (Zeiss) was used to acquire images 2 hours after each behavioral session. Mice were lightly anesthetized with isoflurane and attached to the head mount using a small screw. During the first imaging session, segments of apical dendrites from layer V pyramidal cells were imaged. These segments were acquired within 200 μm from the cortical surface, likely representing dendrites located in layers I and II/III. Imaged segments were generally oriented in the x,y plane of imaging with minimal z-projection. 512×512 pixel images were acquired at 0.5 μm intervals to fully capture the segment of dendrite, and image stacks generally consisting of 30-40 slices. If a segment of dendrite was larger than could be acquired in one 512×512 pixel stack, additional image stacks were sequentially acquired through the x,y,z plane of the dendrite in question so that its full extent could be visualized. The same segments were repeatedly imaged across experimental days by locating their position via a coordinate system established during the first imaging session.

#### Image and Data Analysis

Dendritic spines were analyzed and counted by established criteria. Specifically, the Spine Analysis software included in ScanImage was used to open all imaging days for a given segment of dendrite. A segment was classified as the entire visible length of a piece of a dendrite, and segments were often followed across several images. The presence, gain, and loss of spines was quantified across days for each segment, and all segments were examined for a given animal. Importantly, all images were coded following the completion of the experiment so that the experimenter was blind to the training status of all mice while analyzing and counting spines. Dependence between new spines added to a dendritic segment following various imaging sessions was calculated using Spearman’s correlation and mutual information. Spearman’s rho (ρ) was used as the spine addition/loss data did not follow a normal distribution. Bonferroni method was used to correct for multiple comparisons. For mutual information analysis, statistical significance was calculated by comparing the observed value to the z-score of the chance distribution. A distribution of chance values was calculated by randomly permuting the number of spines added during the second imaging session (10,000x).

Clustering ratios were calculated as the number of clustered spines divided by the total number of new spines gained between two time points. Clustered spines were defined as a new spine that was less than 5 µm from another new spine. For the resampling analysis of clustering, the number of new spines added per segment of dendrite was used to pick an equivalent number of random positions along the same segment (regardless of whether a spine was recorded on that spot on a previous imaging sessions) and assess whether these positions were within 5 μm of each other. When this was completed for all dendrites for a given animal, the percent of clustered spines was calculated as the number of randomly selected new spine positions within 5 μm of each other divided by the total number of stably added new spines for that animal. In turn, each animal’s resampled clustering percentage was calculated, and then these values were averaged together. This completed one resampling event, and this process was then repeated for a total of 1000 resampling events, which then yielded the full distribution of random sampling (Figure S15).

#### Cross Clustering across exposures

The number of clustered spines added following a context exposure were randomly distributed on the dendritic segments from that mouse (10,000x). The percentage of clustered spines added to a dendritic segment following the first context exposure, that were added to a segment that also gained clustered spines following the subsequent context exposure, were measured and compared to the shuffled distribution obtained from the above analysis. Distance between two newly formed spines following each imaging session was calculated for spine pairs that were the nearest neighbors. If no new spine was added or the no newly formed spines persisted during the final imaging session (reference session), these dendrites were not considered during the analysis. Our results remain the same when dendrites with non-persistent or no newly added spines are included in the analysis (5h: 32.1%, average distance between nearest neighbors = 18.1 ± 2.2μm; 7d: 11.1%, average distance between nearest neighbors = 30.9 ± 2.3μm; p < 0.0001). In this case, the length of the dendritic segment was considered the average distance between nearest neighbors.

Resampling Analysis (Figure 5 e-g): Dendritic branches (n = 40) from each condition were subsampled (10,000X) to obtain cumulative frequency distribution for Spearman Correlations, Mutual Information, and spine clustering probability for each condition. Insets demonstrate the difference between observed measurements for each variable from context exposure and HC groups imaged at the 5h interval. P values were calculated as: (Number of measurements where the difference between experimental vs control group < 0/10,000).

### Functional two-photon Imaging

Mice underwent bilateral injection of diluted GCaMP6f (final concentration ~10^11^ vg/mL) in the RSC to achieve semi-sparse infection of layer V RSC neurons ^106^ using the stereotactic coordinates: −2.3 mm posterior to bregma, 0.5 mm lateral to the midline, and −0.8 mm ventral to the skull surface. All dendritic imaging experiments were completed within 25 days of virus injection to prevent viral overexpression. A square 3mm x 3mm craniotomy spanning the midline and hence revealing both RSCs was then made over the injection. After cleaning the surgical site with saline, the coverslip was placed on the dural surface and fastened with adhesive and dental acrylics to expose a square window. Next, an aluminum bar with a threaded hole was attached to stabilize the mice during imaging sessions. Two to three weeks following the surgery, mice underwent handling (3 days) and habituation (3 days) to acclimate to the treadmill and head-fixation. Neuronal and dendritic calcium activity was imaged in head-fixed mice that were free to run on a head-fixed setup.

We recorded dendritic signals during context exposure as well as those evoked spontaneously using a resonant-scanning two-photon microscope (Neurolabware) controlled by Scanbox acquisition software. Distinct contexts were created by immobilizing the mice either on a running wheel, a treadmill, or a horizontal disc (Figure S10), in addition to distinct auditory, olfactory and visual cues associated with each context. Visual stimuli were presented on a large LCD monitor directly in front of the animal and 18 cm from the eye. Visual stimuli consisted of non-repeating natural movies with intermittent gray screens (9s on, 14s off). Spontaneous response data was collected with a blank gray screen in the absence of auditory and olfactory cues. A Coherent Discovery laser (Coherent Inc.) was used for GCaMP excitation, fixed at a wavelength of 920 nm. The objective used was a 16x water-immersion lens (Nikon, 0.8NA, 3 mm working distance). Image sequences were captured at 15.5 Hz at a depth of 30-50 µm below the brain surface for apical tuft dendrites and 320-450 µm for layer V RSC neurons in separate animals.

Collected data were processed using the Suite2P analysis pipeline ^107^. Recorded frames were aligned using a non-rigid motion correction algorithm. Following alignment, any frames with significant motion in the z-axis were dropped from the original video and the data were reanalyzed. Regions of interest (representing dendritic segments) were segmented in a semi-automated manner using a Suite 2p based classifier. Dendritic segments were matched across imaging sessions using an open-source algorithm (https://github.com/ransona/ROIMatchPub, matching criteria: correlation: 0.4). The percentage of reactivated dendrites was defined as the number of matched segments normalized to the average number of dendritic segments detected in each imaging session.

### Hierarchical clustering of dendritic ROIs

To account for global dendritic calcium events and/ or back-propagating action potentials that invade more than one dendritic ROI from the same neuron, we merged any dendritic ROI with highly correlated calcium transients into a single dendritic segment. We adapted a hierarchical clustering method ^108^ previously used to assign axonal boutons to the same source with some variations. Briefly, we generated a sparse activity matrix by thresholding calcium transients from each ROI such that only frames with activity three standard deviations above the mean activity were retained. The time course of calcium transients for each ROI was then cross-correlated with all other ROIs during the same session to generate a matrix of Pearson correlation coefficients between all ROI pairs. This matrix was thresholded in two ways to obtain a sparse matrix. Only those correlation coefficients that were either larger than 0.7 or exceeded 2.5 x standard deviations above the mean value of all the coefficients between this ROI and all others were used. If neither of these conditions were met for a given ROI pair, the associated correlation coefficient was set to 0. The cosine similarity between every ROI pair was then computed from the thresholded matrix of Pearson correlation coefficients.

Next, we classified ROIs with similar activities into clusters using agglomerative hierarchical clustering based on the pairwise distance, computed as ‘1 – cosine similarity’ and the weighted-pair group method with arithmetic means (WPGMA) algorithm. To choose a distance cutoff at which ROIs were considered in the same cluster (i.e., same dendritic segment), we generated a correlation matrix using a shuffled distribution for each animal. The time course of calcium activity from each ROI from each mouse was circularly shuffled by a random amount. This procedure essentially ensures uncorrelated activity in all ROIs and the cutoff value that yielded at least one inaccurate cluster in less than 5 percent of the trails (500 trails) was used as the cutoff for that animal (mean cutoff value = 0.13 ± 0.01). To ensure that clustering criteria were not lenient, we also used singular cutoff values −0.15 or 0.3 – to cluster less correlated ROIs. These criteria when used for all animals resulted in similar results (*p* < 0.001). The clustering method yielded ROI clusters with highly correlated within-cluster activity across sessions (reference and comparison sessions for reactivated ROIs; Figure S12c).

Analysis of correlated dendritic activity: Dendritic activity/events were estimated via non-negative temporal deconvolution of the Suite-2p extracted signal using Vanilla algorithm ^106,109^. Event probabilities were then binarized to reflect active frames and number of active frames were used to calculate an event rate. To account for variations in the number of reactivated ROIs in imaging sessions 5 hours and 7 days apart, we randomly subsampled 30 reactivated ROI pairs for each comparison (500x) to generate a probability distribution (compared using a ks test).

### Slice Preparation

Adult cFos-tTa mice were injected with 500 nl of AAV1-TRE-hChR2-mCherry or AAV1-TRE-hChR2-mCherry-DTE. After three or more weeks, mice were taken off doxycycline for three days, allowed to explore a novel context for 10 min and 24 hours later were deeply anaesthetized with isoflurane and decapitated. The brain was rapidly dissected out and transferred to oxygenated (95% O_2_ / 5% CO_2_), ice-cold cutting solution containing (in mM): 92 choline, 2.5 KCl, 1.2 NaH_2_PO_4_, 30 NaHCO_3_, 20 HEPES, 25 glucose, 2 Thiourea, 5 Na-ascorbate, 3 Na-pyruvate, 5 N-acetyl-L-cysteine, 0.5 CaCl_2_ and 10 MgSO_4_. Coronal slices (300 µm thick) containing the retrosplenial cortex were cut using a Leica VT1200 vibrating blade microtome, transferred to a submerged holding chamber containing oxygenated cutting solution and allowed to recover for 15 min at 34°C. Following recovery, the slices were transferred to an oxygenated solution containing (in mM): 92 HEPES, 2.5 KCl, 1.2 NaH_2_PO_4_, 30 NaHCO_3_, 20 HEPES, 25 glucose, 2 Thiourea, 5 Na-ascorbate, 3 Na-pyruvate, 5 N-acetyl-L-cysteine, 2 CaCl_2_ and 2 MgCl_2_ and allowed to recover further for 1hr. Following incubation, slices were transferred to a superfused recording chamber and constantly perfused with oxygenated aCSF containing (in mM): 115 NaCl, 10 glucose, 25.5 NaHCO_3_, 1.05 NaH_2_PO_4_, 3.3 KCl, 2 CaCl_2_ and 1 MgCl_2_ and maintained at 28°C. For two mice (TTA-ChR2 and TTA-ChR2-DTE, n = 1 each), brains were sliced in recording solution.

### Whole-cell patch recordings

Whole cell current-clamp recordings were performed on pyramidal neurons in the RSC using pipettes (3-5MΩ resistance) pulled from thin-walled Borosilicate glass using a Sutter P97 Flaming/Brown micropipette puller and filled with an internal solution containing (in mM) 110 K-Gluconate, 20 KCl, 2 MgCl_2_, 10 HEPES, 10 Na_2_-ATP and 0.3 Na_2_-GTP and 10 Na-phosphocreatine. All recordings were obtained using a MultiClamp 700B amplifier controlled by the pClamp 10 software and digitized using the Digidata 1440A system. Signals were filtered at 10kHz and digitized at 20kHz. Neurons were included in the study only if the initial resting membrane potential (Vm) ≤ −55 mV, access resistance (Ra) was <25MΩ and were rejected if the Ra changed by >20% of its initial value. For all recordings, neurons were held at −60 mV. The stable resting membrane potential of neurons were measured and averaged over a 60s duration with 0mA current injection immediately after breaking in. Input resistance was measured as the slope of the steady-state voltage response to increasing current injections (−50pA to 50pA, Δ=10pA). To investigate the response of the neurons following optogenetic stimulation, 473nm LED (5 ms pulses, 5 Hz) was delivered through Cool LED pE-300 and neuronal response was calculated. Only neurons with a visible response to optogenetic stimulation were included in analysis (n = 12 from > 50 RSC neurons). All mCherry-positive RSC neurons from TTA-ChR2 mice resulted in APs following stimulation. The recordings were analyzed using Clampfit and Matlab.

### Computational modeling

We adapted a previously published model network of memory allocation ^51^. The model network consists of populations of excitatory and inhibitory neurons, which are modeled as 2-level integrators to account for dendrites. Increases in dendritic and somatic excitability are assumed to follow the time course of CREB activation. CREB activation, in turn, is assumed to be triggered when calcium accumulation exceeds a threshold.

Neurons consist of a somatic spiking unit connected to multiple independent dendritic subunits. Dendrites and soma are modeled using simplified integrate-and-fire model dynamics, where the somatic unit includes adaptation current, and dendrites and soma are coupled to it via axial resistance. Inhibitory neurons provide feedback inhibition and are separated into 2 equal sub-populations, soma-targeting and dendrite-targeting. The voltage of each dendritic subunit is as follows:

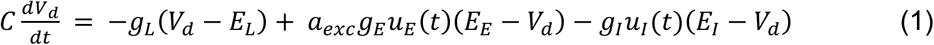

where *V_d_* is the dendritic membrane potential, *C* is the membrane capacitance, *E_E_* is the reversal potential for excitatory receptors, *E_I_* is the reversal potential for inhibitory receptors, *E_L_* is the resting potential (0mV), *a_exc_* is the dendritic excitability level parameter, *g_L_* is the leak conductance, *g_E_*, *g_I_* are the maximal excitatory and inhibitory synaptic conductances. *u_I_(t) and u_E_(t)* are the instantaneous activations of excitatory and inhibitory synapses on the dendrite respectively:

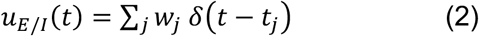

where *w_j_* is the weight of synapse *j* and *t_j_* are the timings of incoming spikes.

Somatic voltage is modeled as an Integrate-and-Fire model with adaptation as follows:

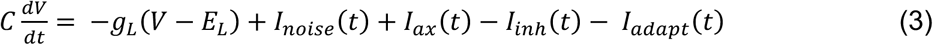

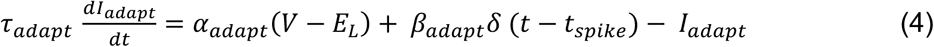

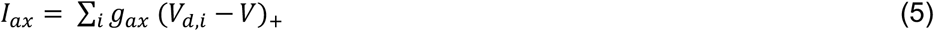

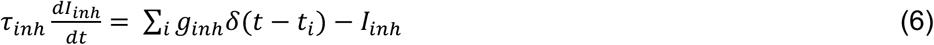

where *V* is the somatic voltage, *I_noise_* is uniform noise current (max amplitude 500 pA), *I_ax_* is the excitatory axial current, *I_inh_* is the filtered inhibitory current from somatically-targeting interneurons, *I_adapt_* is the adaptation current, *τ_adapt_* is the adaptation time constant, *α_adapt_* the adaptation coupling parameter, *β_adapt_* is the amount by which adaptation current increases every time the neuron spikes, *g_ax_* is the axial resistance, *τ_inh_* is the time constant of inhibitory current and *g_inh_* the inhibitory current scaling constant. Somatic spiking occurs when the somatic voltage reaches the spike threshold *θ_soma_*. On an incoming spike, synaptic and dendritic branch calcium is increased by an amount *ΔCα(V_d_)* that depends on the instantaneous depolarization *V_d_* of the dendritic branch to account for the Magnesium blocking of NMDA receptor:

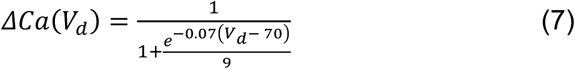

Synaptic inputs representing memories to be encoded are initially allocated randomly to the dendritic subunits of excitatory neurons with initial weight ranging uniformly randomly between 0.16 and 0.36. In addition, feedback synapses between excitatory and inhibitory populations are allocated at random, with separate distributions for soma-targeting and dendrite-targeting interneurons (see Supplementary Table 2). In the case of dendritic separation between memories (Figure 7g) the memories were allocated at random dendrites but were not allowed to overlap in the same dendrite.

The total calcium influx during memory encoding determines synaptic plasticity, changes in excitability and the levels of plasticity-related proteins (PRPs) after encoding. In order to replicate the experimental observation that the number of new synapses correlates positively with the existing potentiated synapses within the same dendrite, we introduce stochasticity in synaptic tagging as described below. Synapses that receive calcium influx above zero during stimulation are selected for potentiation or depression. If the synapse resides on a neuron that is highly activated (spiking with frequency > 10 Hz during stimulus presentation) then the synapse is tagged for potentiation with probability *p_LTP_= 0.29 + X_dend_*N_s_/2,* otherwise the synapse is tagged for depression. *N_s_* is the number of preexisting potentiated synapses in the same dendrite and *X_dend_* is the excitability of the dendrite (see below). Synaptic tags decay exponentially with time constant of 1 hour.

When the somatic calcium level of a neuron exceeds *Θ_PRP_* at time *T*, the level of plasticity-related proteins of the neuron is elevated according to the following function:

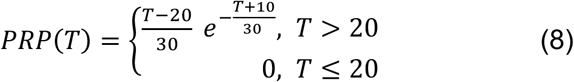

where *T* is the time since the stimulus in minutes. When synaptic tags and PRPs have values above 0.1, the weights *w* of synapses are updated by *Δw = 0.15*PRP(t)*(synaptic tag)*, where t is time in seconds. Synaptic weights are bounded in the interval [0, 1]. We note that, because the delay between memories in all simulations is 5 hours or more, the PRP protein levels are always sufficient for full synaptic tag consolidation, and thus there was no competition for PRPs that could affect the linking of memories.

#### Excitability within the Linking model

Dendritic and somatic excitability are assumed to be mediated via the dynamics of CREB activation after learning. The increased excitability after learning is triggered when calcium accumulation exceeds a threshold. The time course of increased excitability is modeled as a sum of two sigmoids and exceeds 24 hours ^110^ as shown below

When the total dendritic and somatic calcium are above thresholds *Θ_dend_* and *Θ_soma_* respectively, the excitability level *Χ* of the dendrite/soma is increased by the amount given by:

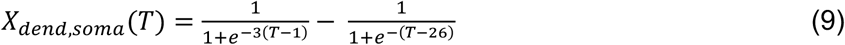

where *T* is the time in hours since the stimulus. The excitability level parameter *a_exc_* is increased by 10% when *X_dend_* > 0.1, while the adaptation reset parameter *β_adapt_* is increased by 28% when *X_soma_* > 0.1. For the simulations of the Linking model without dendritic mechanisms, the dendritic excitability was set at *X_dend_= 0* and the probability *p_LTP_* was kept constant at *0.32*.

Synaptic weights are additionally subject to a homeostatic synaptic scaling rule, which adjusts synaptic weights *w_j_* to normalize the total synaptic input to each neuron with time constant *τ_H_*:

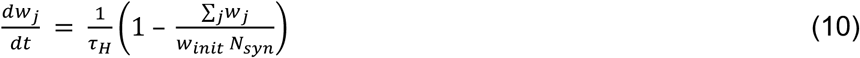

where *w_init_* is 0.3 and *N_syn_* is the total number of incoming synapses to the neuron.

#### Stimulation protocol

For every memory being encoded, the synaptic inputs which represent the memory are stimulated for 4 seconds with firing rate 35Hz in order to drive the initially weak synapses. After the first memory encoding, a delay period is simulated for 5 hours, 2 days or 7 days, and then the second memory is encoded. Memories are recalled by being stimulated again after 2 days.

For the clustering statistics, a branch was considered to contain overlapping clusters if it contained at least 3 potentiated synapses from each memory. The parameters used in the model are listed in Supplementary Table 2. The model was written in C++ and data analysis was done with Python/numpy. The source code for the simulation, data analysis and scripts to reproduce the data and figures are available at https://dendrites.gr/wp-content/uploads/2022/08/rsc_model2.zip. Chance levels for neuronal overlap was calculated as previously described ^12^: Chance Overlap = [(Neuronal ensemble encoding A x Neuronal ensemble encoding B)/100]. Percent above chance overlap = (Observed Overlap – Chance Overlap)/Chance Overlap.

### Quantification and Statistical Analyses

The investigator who collected and analyzed the data including behavior, imaging and staining was blinded to the mouse genotypes and treatment conditions. Error bars in the figures indicate the SEM. All statistical analyses were performed using GraphPad Prism 9 or Matlab. For behavior and imaging experiments, n designates the number of mice unless otherwise mentioned. Statistical significance for behavioral manipulations was assessed using parametric tests (Student’s t test, or one- or two-way ANOVA) followed by the indicated post-hoc tests (GraphPad Prism 9 recommended post-hoc tests) as data followed a Gaussian distribution. The level of significance was set at *p* < 0.05 unless Bonferroni’s correction for multiple comparisons was used.

## Acknowledgements

We thank P. Riazi, J. Lin, E. Rubin, G. Padda, Y. Cai, M. Zhou, D. J. Cai, D. Aharoni, C.E. Yaeger, A. F. de Sousa, A. Chowdhury and Y. Shen for advice and technical support. We thank Dean Buonomono, Mayank R. Mehta, Peyman Golshani and James M. Otis for helpful discussions. This work was supported by grants from the NIMH (R01 MH113071), NIA (R01 AG013622), and from the Dr. Miriam and Sheldon G. Adelson Medical Research Foundation to AJS. The computational modeling work was supported by the European Commission (H2020-FETOPEN-2018-2019-2020-01, FET-Open Challenging Current Thinking, GA-863245), the NIH (R01MH124867-01) and the Einstein Foundation Berlin. We acknowledge resources from the Campus Microscopy and Imaging Facility (CMIF) and the OSU Comprehensive Cancer Center (OSUCCC) Microscopy Shared Resource (MSR), The Ohio State University (RRID:SCR_025078). This facility is supported in part by grant P30 CA016058, National Cancer Institute, Bethesda, MD.

## Author contribution

MS and AJS did experimental design, data acquisition and analyses, drafting and revising the article; JT helped with experimental design for two-photon calcium imaging experiments. SM, IDM, AP helped with behavioral data acquisition; SK, JL, WDH made the TRE-dependent constructs; DAF helped with Miniscope data analysis and interpretation; AL and SH helped with two-photon data analyses and interpretation; GK and PP designed, implemented, and analyzed the computational modeling experiments. All authors read and edited the manuscript.

## Competing interests

The authors declare no competing interests.

## Notes

http://modeldb.yale.edu/267175

### Competing Interest Statement

The authors have declared no competing interest.

### Summary of Updates

Extensive revision with the addition of new data. Revision to existing figures as well as addition of several figures

